# Microglia Mediate Early Corticostriatal Synapse Loss and Cognitive Dysfunction in Huntington’s Disease Through Complement-Dependent Mechanisms

**DOI:** 10.1101/2021.12.03.471180

**Authors:** D.K. Wilton, K. Mastro, M.D. Heller, F.W. Gergits, C R. Willing, A. Frouin, A. Daggett, X. Gu, A.Y. Kim, R. Faull, S. Jayadev, T Yednock, X.W. Yang, B. Stevens

## Abstract

Huntington’s disease (HD) is a devastating monogenic neurodegenerative disease characterized by early, selective pathology in the basal ganglia despite the ubiquitous expression of mutant huntingtin. The molecular mechanisms underlying this region-specific neuronal degeneration and how these relate to the development of early cognitive phenotypes are poorly understood. Here, we show that there is selective loss of synaptic connections between the cortex and striatum in postmortem tissue from HD patients that is associated with the increased activation and localization of complement proteins, innate immune molecules, to markers of these synaptic elements. We also find that levels of these secreted innate immune molecules are elevated in the CSF of premanifest HD patients and correlate with established measures of disease burden.

In preclinical genetic models of HD we show that complement proteins mediate the selective elimination of corticostriatal synapses at an early stage in disease pathogenesis marking them for removal by microglia, the brain’s resident macrophage population. This process requires mutant huntingtin to be expressed in both cortical and striatal neurons and inhibition of this complement-dependent elimination mechanism through administration of a therapeutically relevant C1q function blocking antibody or genetic ablation of a complement receptor on microglia, prevented synapse loss, increased excitatory input to the striatum and rescued the early development of visual discrimination learning and cognitive flexibility deficits in these models. Together, our findings implicate microglia and the complement cascade in the selective, early degeneration of corticostriatal synapses and the development of cognitive deficits in presymptomatic HD, and also provide new preclinical data to support complement as a therapeutic target for early intervention.

## Introduction

Huntington’s disease (HD), is the most common autosomal dominant neurodegenerative disease. It is characterized by progressive motor, cognitive and psychiatric symptoms, with onset of the manifest phase defined as the point when patients develop a movement disorder that meets clinical thresholds. Typically this occurs around 45 years of age (but can be highly variable), with the disease course usually characterized by inexorable progression, ultimately resulting in death from disease related complications^1^. Currently, there are no therapies that modify disease onset or progression.

HD is caused by a CAG repeat expansion mutation encoding a polyglutamine (PolyQ) tract in the *Huntingtin* (*HTT*) gene^2,3^. Based on this genetic finding, multiple transgenic animal models have been generated to interrogate the underlying biology of the disease^4,5^; however, the molecular mechanisms that drive the early and selective degeneration of basal ganglia circuits and how these relate to early cognitive phenotypes remain poorly understood^6-23^. The corticostriatal pathway, which connects intratelencephalic (IT) and pyramidal tract (PT) neurons in the cortex with medium spiny neurons (MSNs) and cholinergic interneurons (ChIs) in the striatum, is affected at very early stages of disease progression^19,24^. Electrophysiological recordings in mouse models and structural and functional imaging of HD patients in the premanifest stage of the disease reveal altered white matter structure and functional connectivity in this pathway which in patients correlates with a more rapid cognitive decline^18,25-29^. At later stages of the disease, reductions in synaptic marker levels suggest that corticostriatal synapses are lost; but it is unknown whether this synapse loss occurs prior to the onset of motor and cognitive deficits or why corticostriatal synapses are selectively vulnerable. Moreover, the molecular mechanisms that mediate synapse loss in HD remain unclear^22^.

Studies in mouse models of Alzheimer’s disease and frontotemporal dementia have demonstrated a link between synaptic loss and components of the classical complement cascade, a group of secreted ‘eat me’ signals that mediate the recognition and engulfment of synaptic elements by microglia during development and in disease contexts^30-36^. While this mechanism of synaptic elimination has never been explored in HD, transcriptional profiling, brain imaging studies and analysis of patient-derived serum and cerebrospinal fluid (CSF) have all indicated an altered neuro-immune state in premanifest HD patients^37-44^. In mouse models of HD, microglia have also been found to display changes in morphology, deficits in motility and an altered transcriptional profile during the symptomatic phase of the disease^45-53^. Separately, transcriptomic studies have identified an increase in the expression of complement proteins and their regulators in the basal ganglion of postmortem tissue from HD patients^54-57^, suggesting complement proteins and microglia are dysregulated in HD. However, neither complement nor microglia have been studied at early stages of the disease, and it is unknown whether they play any roles in the pathogenesis of synapse loss or the development of early cognitive deficits in HD.

In this study, by integrating findings from postmortem HD brain samples and two preclinical HD mouse models, we provide evidence that microglia and complement coordinate to selectively target corticostriatal synapses for early elimination in the dorsal striatum; a process that requires mutant HTT (mHTT) expression in both cortical and striatal neurons. Inhibition of this synaptic elimination through administration of a therapeutic C1q function blocking antibody (ANX-M1; Annexon Biosciences) or genetic ablation of microglial complement receptor 3 (CR3/ITGAM) reduces loss of corticostriatal synapses and improves visual discrimination learning and cognitive flexibility impairments at early stages of disease progression in HD models. We further show that aspects of this pathological synapse elimination mechanism may be operating in premanifest HD patients, as complement protein levels in the CSF of HD patients correlate with an established predictor of both pathological severity and disease onset.

## Results

### Progressive and selective loss of corticostriatal synapses in HD patients is associated with complement activation and changes in microglia

To test whether there is specific loss of corticostriatal synapses in HD patients, we assessed glutamatergic excitatory synapses in postmortem tissue from the caudate nucleus (part of the striatal structure) and cerebellum of control individuals (no documented evidence of neurodegenerative disease; see methods and **supplemental table 2**) and HD patients with different Vonsattel grades of striatal HD neuropathology^58-61^. Immunohistochemical (IHC) analysis revealed a progressive and significant loss of corticostriatal synapses, as denoted by the colocalization of the excitatory postsynaptic marker Homer1 and the presynaptic corticostriatal marker VGLUT1, in the disease-affected caudate nucleus of the HD tissue relative to that seen in tissue from control individuals **(Fig 1a**,**b)**. Conversely, we observed no difference in the numbers of VGLUT1 positive glutamatergic synapses in the cerebellum, which is less affected in HD **(Fig. 1c)**.

**Figure 1:**
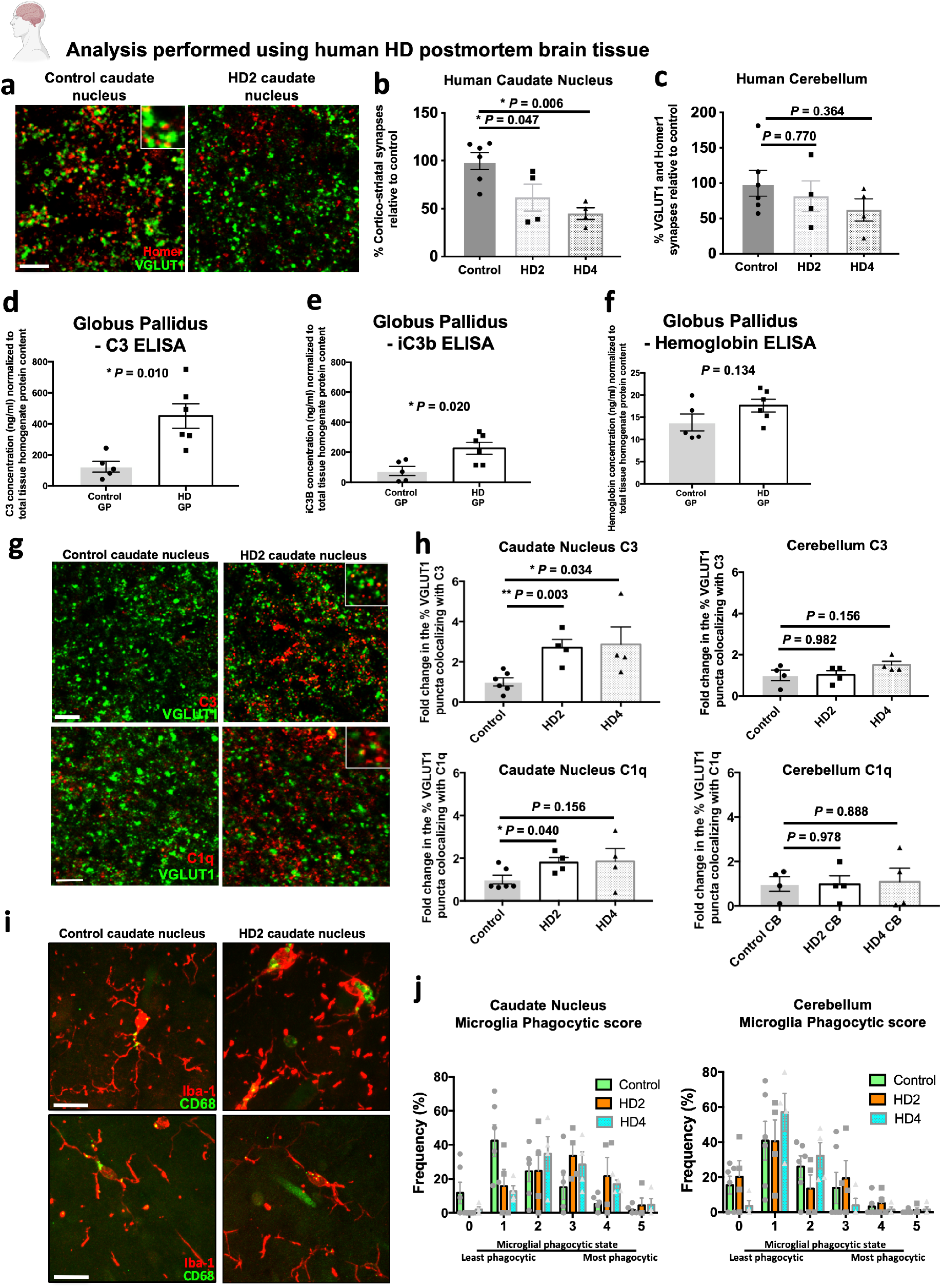
Loss of specific synaptic populations, activation and association of complement proteins with synaptic elements and adoption of a more phagocytic microglial state are evident in postmortem brain tissue from HD patients. (a) Representative confocal images showing staining for the corticostriatal specific presynaptic marker VGLUT1 and the postsynaptic density protein Homer1 in the caudate nucleus of postmortem tissue from a control (no documented evidence of neurodegenerative disease; see methods and supplemental table 2) individual and the caudate nucleus from an HD patient, which has been assessed to be Vonsattel grade 2. Note the reduced number of both synaptic markers in the HD tissue. Scale bar = 5 μm (b) Bar chart showing quantification of corticostriatal synapses (colocalized VGLUT1 and Homer1 puncta) in the caudate nucleus of control, Vonsattel grade 2 HD and Vonsattel grade 4 HD tissue. In the HD tissue there is a decrease in the percentage of this synaptic population at both Vonsattel grades relative to that seen in control individuals, n=6 control, n=4 HD with Vonsattel 2 tissue grade and 4 HD with Vonsattel 4 tissue grade. One way anova p=0.0058; Tukey’s multiple comparisons test, control vs HD2 p=0.0474; control vs HD4 p=0.0061; HD2 vs HD4 p=0.537. (c) In contrast the same analysis carried out in the molecular layer of the folia of the cerebellum (an area currently thought to be less impacted by the disease) showed no change in the population of VGLUT1 positive glutamatergic synapses relative to the numbers seen in control individuals, n=6 control individuals, 4 HD Vonsattel grade 2 and 4 HD Vonsattel grade 4. One way anova p=0.393; Tukey’s multiple comparisons test, control vs HD2 p=0.770; control vs HD4 p=0.365; HD2 vs HD4 p=0.791. (d) Bar chart showing ELISA measurements of the concentration of complement component C3 in proteins extracted from the globus pallidus (GP) of postmortem tissue from manifest HD patients and control (no documented evidence of neurodegenerative disease; see methods and supplemental table 2) individuals after normalization for total tissue homogenate protein content, n=5 control GP and 6 HD GP. ANCOVA controlling for the effect of age F(2,10)=8.74, p=0.010 (e) Bar chart showing ELISA measurements of the concentration of complement component iC3b in proteins extracted from the caudate nucleus of postmortem tissue from manifest HD patients and control individuals after normalization for total tissue homogenate protein content, n=5 control GP and 6 HD GP. ANCOVA controlling for the effect of age F(2,10)=6.71, p=0.020. (f) Bar chart showing ELISA measurements of the level of hemoglobin in proteins extracted from the globus pallidus (GP) of postmortem tissue from manifest HD patients and control individuals after normalization for the total tissue homogenate protein content, n=5 Control GP and 6 HD GP. Unpaired t-test p=0.134. (g) Representative confocal images showing staining for VGLUT1 together with either complement protein C3 or C1q in the caudate nucleus of postmortem tissue from a control (no documented evidence of neurodegenerative disease; see methods and supplemental table 2) individual and the caudate nucleus from an HD patient which has been assessed to be Vonsattel grade 2. Scale bar = 5 μm. Note the increased deposition of both complement components in the HD tissue. Insets show examples of colocalization of both complement proteins with presynaptic marker VGLUT1. (h) Bar charts showing quantification of the percentage of VGLUT1 positive glutamatergic synapses associating with C3 and C1q puncta in the caudate nucleus and cerebellum of postmortem tissue from HD patients assessed to be either Vonsattel grade 2 or 4 relative to that seen in tissue from control individuals (no documented evidence of neurodegenerative disease; see methods and supplemental table 2). In the caudate nucleus there is a significant increase in the percentage of VGLUT1 presynaptic terminals that colocalize with complement component C3 in both Vonsattel grade 2 and grade 4 HD tissue, n=6 control individuals, n=4 HD individuals with Vonsattel tissue grade 2 and n=4 HD individuals with Vonsattel grade 4. One way anova p = 0.029. Unpaired t-test control vs HD2 p=0.003; control vs HD4 p= 0.034. This is also the case for C1q and VGLUT1 with an increased % of VGLUT1 puncta colocalizing with this complement protein in the HD tissue relative to that seen in tissue from control individuals, n=6 control individuals, n=4 HD individuals with Vonsattel tissue grade 2 and n=4 HD individuals with Vonsattel tissue grade 4. One way anova, p= 0.181. Unpaired t-test, control vs HD2 p=0.040, control vs HD4 p=0.156. In contrast the same analysis carried out in the molecular layer of the folia of the cerebellum showed no significant increase in the percentage of VGLUT1 presynaptic terminals colocalizing with C3 or C1q in the HD tissue relative to that seen in tissue from control individuals. For C3, n=4 control individuals, n=4 HD individuals with Vonsattel tissue grade 2 and n=4 HD individuals with Vonsattel tissue grade 4. One way anova, p=0.215. Unpaired t-test, control vs HD2 p=0.982; control vs HD4 p= 0.156. For C1q, n=4 control individuals, n=4 HD individuals with Vonsattel tissue grade 2 and n=4 HD individuals with Vonsattel tissue grade 4. One way anova, p=0.981. Unpaired t-test, control vs HD2 p=0.978; control vs HD4 p=0.888. (i) Representative confocal images showing staining for Iba-1 and CD68 in the caudate nucleus and cerebellum of postmortem tissue from a control clinically normal individual and the caudate nucleus from an HD patient (Vonsattel grade 2). Scale bar = 20 μm. Note the shorter and thicker process of the microglia in the caudate of the HD tissue and the increased level of lysosomal marker CD68. In contrast microglia in the cerebellum of tissue from controls and HD patients display similar morphology and lysosomal levels. (j) Bar charts show quantification of microglia phagocytic state based on both changes in morphology and CD68 levels with 5 being the most phagocytic and 0 the least. Microglia in the caudate of the HD tissue show a significant shift towards a more phagocytic state relative to those in the control tissue, n=6 control individuals, n=4 HD individuals with Vonsattel tissue grade 2 and n=4 HD individuals with Vonsattel tissue grade 4. Two way anova, interaction of score and genotype p=0.019. This shift is not observed when the analysis is carried out in the molecular layer of the folia of the cerebellum, n=6 control individuals, n=4 HD individuals with Vonsattel tissue grade 2 and n=4 HD individuals with Vonsattel tissue grade 4. Two-way anova, interaction of score and genotype p=0.367. For bar charts, bars depict the mean. All error bars represent SEM. Stars depict level of significance with *=p<0.05, **p=<0.01 and ***p<0.0001.

To determine whether complement proteins could contribute to this loss of corticostriatal synapses we first developed or optimized ELISA assays to measure the levels of C3 and its activated cleavage component iC3b (a cognate ligand for microglial CR3), which functions downstream in the complement pathway as an opsonin (marking substances for removal by phagocytic cells)^62,63^. Examining extracts from the globus pallidus, a component of the basal ganglion structure^64-66^, these assays showed that levels of both C3 and iC3b were higher in HD tissue relative to age-matched controls despite both sets of extracts displaying similar levels of blood contamination **(Fig. 1d**,**e**,**f)**.

We next tested whether complement proteins associate with corticostriatal synapses in HD brain samples. Immunohistochemical analysis of postmortem tissue from the caudate nucleus and cerebellum of symptomatic HD (Vonsattel grade 2 and 4) patients and age-matched controls revealed increased association of both complement component C1q, an initiator of the classical complement cascade (expressed by microglia **Extended data Fig. 1a**) and C3 (expressed by both astrocytes and microglia **Extended data Fig. 1b**,**c**) with corticostriatal synapses in the disease-affected caudate nucleus of HD brains relative to that seen in age-matched controls. There was, however, no increased association of these complement proteins with glutamatergic synapses in the cerebellum (a less disease-affected region) which correlates with the relative preservation of these structures in this region **(Fig. 1g**,**h; Extended data Fig. 1k)**.

Microglia in the HD tissue set displayed a region-specific shift towards a more phagocytic phenotype relative to that seen in the age matched controls, adopting a more amoeboid morphology and possessing higher levels of the lysosomal marker CD68 **(Fig. 1i**,**j)**. Also, consistent with prior transcriptomic studies we found that protein levels of the microglia-specific complement receptor 3 (CR3) were elevated in the HD globus pallidus relative to that seen in extracts from age matched controls; however, with the current sample size this difference was not statistically significant **(Extended data Fig. 1d)**.

To test whether complement-mediated microglial synaptic elimination could occur in premanifest HD, we performed quantitative RT-PCR on RNA extracted from two rare samples of caudate nucleus from premanifest HD patients and found that transcript levels of C3 and CR3 were elevated relative to controls **(Extended data Fig. 1e**,**f)**, consistent with a previous study that used unbiased transcriptomic profiling in the BA9 region of premanifest HD patients^55^. Together these results demonstrate that corticostriatal synapses are selectively and progressively lost in postmortem tissue from HD patients and that this loss is accompanied by an increase in complement protein levels, activation and synaptic localization of complement proteins as well as phenotypic changes in microglia that suggest complement-mediated synaptic elimination might be contributing to this synaptic pathology.

### Early and specific loss of corticostriatal synapses in HD mouse models

To further explore the mechanisms underlying this synaptic pathology and determine whether corticostriatal synapses are selectively vulnerable early in disease, we quantified corticostriatal synapses together with the thalamostriatal synapse population (the other significant source of excitatory input to the striatum)^67-69^, in the dorsolateral striatum of the zQ175 knock-in and BACHD human genomic transgenic mouse models of HD^70,71^. Both models develop a variety of electrophysiological abnormalities and have a similar time course of striatal and cortical atrophy, with zQ175 mice showing relatively mild motor deficits at around 7 months (mo) of age^17,19,22,70-76^. Consistent with previous functional studies that suggested a reduction of glutamatergic inputs onto MSNs, we found ∼50% fewer corticostriatal synapses in the dorsolateral striatum of 7 mo zQ175 mice (using both super-resolution and confocal imaging approaches)^72,74,77,78^ **(Fig. 2a**,**b**,**c)**. Interestingly, this loss was not observed in the dentate gyrus of the hippocampus, a less disease affected brain region **(Fig. 2d)** and was replicated in the BACHD model **(Extended data Fig. 2a**,**b)**. Concordantly, immunoblot analysis found reduced levels of synaptic markers in the striatum but not in the less disease-affected cerebellum in 7 mo zQ175 mice **(Extended data Fig. 2c**,**d**,**e**,**f**,**g**,**h**,**i)**.

**Figure 2:**
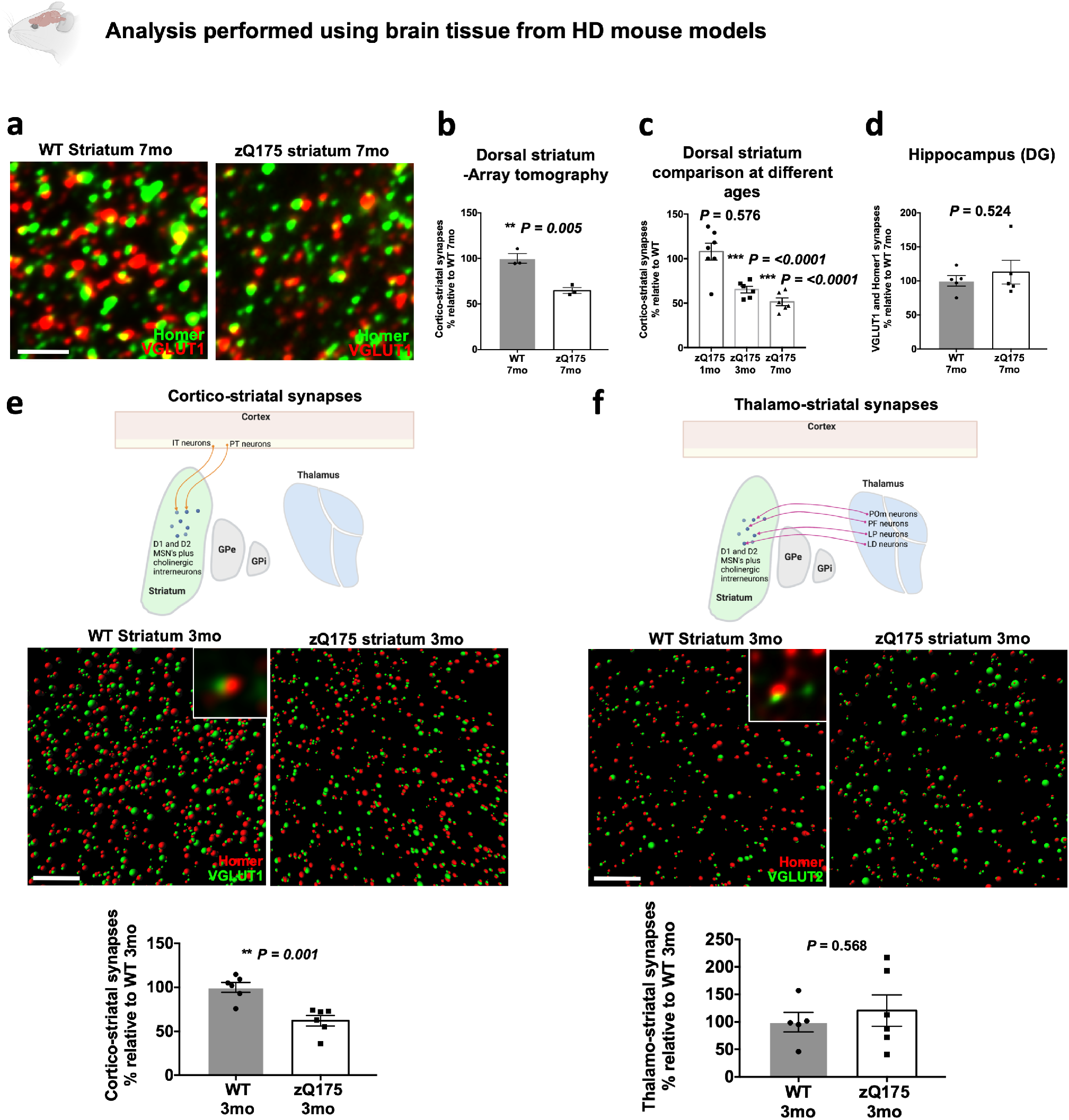
Early and selective loss of corticostriatal synapses in Huntington’s disease. **(a)** Representative fluorescence photomicrographs of array tomography projections from the dorsolateral striatum of 7 mo zQ175 and WT littermates stained with antibodies to Homer1 and VGLUT1. Note the significant reduction in colocalized pre and postsynaptic puncta (yellow overlap) in the image from the zQ175 striatum. Scale bar = 3 μm **(b)** Imaris and Matlab quantification of synapse numbers in the array tomography projections. There is a significant reduction in the number of corticostriatal synapses in the dorsal striatum of the zQ175 mice relative to those seen in their WT littermates, n=3 WT mice and 3 zQ175. Unpaired t-test p=0.0048. **(c)** Quantification of corticostriatal synapses (imaged with confocal microscopy) in the dorsolateral striatum of zQ175 mice at different ages expressed as a % of WT numbers at the same age. Note that at 1 mo synapse numbers are comparable to WT but there is a significant reduction at both 3 and 7 mo, n=7 WT mice and 7 zQ175 mice at 1 mo; n=6 WT and 6 zQ175 mice at 3 mo; n=6 WT and 6 zQ175 mice at 7 mo. Unpaired t-test relative to WT at each age at 1 mo p=0.576, at 3 mo p=<0.0001, at 7 mo p=<0.0001. **d)** Quantification of VGLUT1 labeled glutamatergic synapses in the molecular layer of the dentate gyrus of the hippocampus shows no significant difference between zQ175 mice and WT littermates at 7 mo of age, n=5 WT mice and 5 zQ175 mice. Unpaired t-test p=0.524. **(e)** Imaris processed structured illumination images (SIM) show a significant reduction in the number of corticostriatal synapses in the dorsolateral striatum of 3 mo zQ175 mice. Pictograms show synapses defined as pre and postsynaptic spheres (rendered around immunofluorescent puncta) whose centers are less than 0.3 μm apart. Inset shows a representative example of colocalized pre and postsynaptic puncta. Scale bar = 5 μm. Bar chart shows Matlab quantification of corticostriatal synapses, n=6 WT mice and 6 zQ175 mice. Unpaired t-test p=0.001. **(f)** Imaris processed structured illumination images showing no difference in the numbers of thalamostriatal synapses at 3 mo as denoted by staining with the presynaptic marker VGLUT2. Scale bar = 5 μm. Bar chart shows quantification of these images, n=5 WT and 6 zQ75 mice. Unpaired t-test p=0.568. For bar charts, bars depict the mean. All error bars represent SEM. Stars depict level of significance with *=p<0.05, **p=<0.01 and ***p<0.0001.

To test whether synapse loss occurs before the onset of motor and cognitive deficits, we repeated the analysis in 3 mo zQ175 mice and still saw a significant reduction of corticostriatal synapses relative to WT littermates **(Fig. 2c**,**e)**. Importantly, this reduction of synapses did not appear to result from a developmental failure in synapse formation as, consistent with previous findings, no difference could be detected at 1 mo of age in these mice **(Fig. 2c)**^79^.

The striatum receives excitatory inputs from both the cortex and thalamus, which can be distinguished by staining for the presynaptic vesicular proteins VGLUT1 and VGLUT2, respectively^67,68,80-84^. Using antibodies to these markers, we found that corticostriatal synapses but not thalamostriatal synapses were lost in 3 mo zQ175 mice **(Fig. 2e**,**f)**. Only when the mice were old enough to display motor deficits were both synaptic populations reduced, in line with previous reports in other HD models^70,85-87^ **(Extended data Fig. 2j**,**k)**. One group has reported fewer thalamostriatal synapses in a different region of the striatum on postnatal day 21 (P21) and P35 (although they also used a different combination of synaptic markers which have been shown by others to be dependent on synaptic maturity for their localization at synaptic contacts^79,88^). This result presumably reflects a developmental failure of synapse formation or elimination, but our data showing no differences at 3 mo suggests any such deficits are resolved or that there are differences in the synaptic populations being surveyed.

Consistent with the selective vulnerability of corticostriatal synapses, immunoblot analysis of two rare samples of caudate nucleus from premanifest HD patients also showed a reduction in levels of the corticostriatal marker VGLUT1 and the postsynaptic marker PSD95 but not in the thalamostriatal marker VGLUT2 **(Extended data Fig. 2l)**, further suggesting selective vulnerability of the corticostriatal connection in HD. Taken together, these results show that corticostriatal synapse loss is an early event in the pathogenesis of mouse models of HD, occurring before the onset of motor and cognitive deficits.

### Complement components are upregulated and specifically localize to vulnerable corticostriatal synapses in HD mouse models

Emerging research implicates classical complement cascade activation in synaptic elimination both during normal development and in disease and injury contexts^30-36^. To test whether complement proteins could mediate the selective loss of corticostriatal synapses, we investigated whether levels of C1q and complement protein C3, were elevated in disease-vulnerable brain regions. IHC analysis revealed that, similar to our findings in the post-mortem caudate nucleus of HD patients levels of C1q and C3 were both elevated in the striatum and motor cortex of 7 mo zQ175 mice, but not in the less disease-affected hippocampus (dentate gyrus) at this time ^17,89^ **(Fig. 3a**,**b**,**d**,**e)**. As determined by *in situ* hybridization, and in line with other studies, we found C1q to be predominantly expressed by microglia in the striatum of zQ175 mice and C3 to be expressed by Acta2 expressing ependymal cells lining the wall of the lateral ventricle **(Extended data Fig. 3h**,**i)**^90,91^. Both complement proteins were also significantly elevated at 3 mo in the dorsolateral striatum of zQ175 mice, correlating with the earliest time point that we observed fewer corticostriatal synapses **(Fig. 3c**,**f)**. These findings were replicated in the BACHD model **(Extended data Fig. 3a**,**b)**.

**Figure 3:**
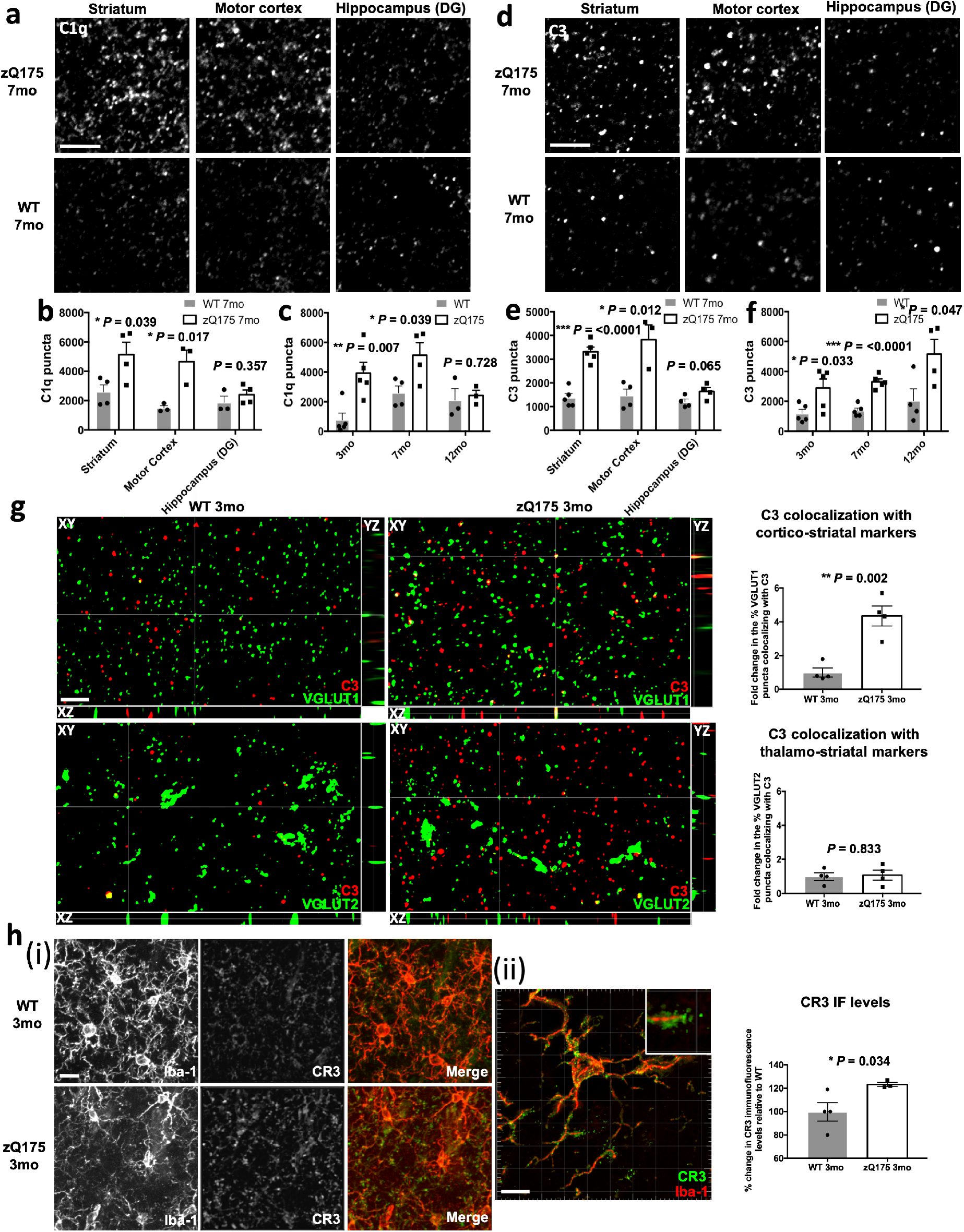
Complement proteins associate with specific synaptic connections and microglia increase their expression of complement receptors in HD mouse models. **(a)** Representative confocal images showing C1q staining in disease affected (dorsolateral striatum and motor cortex) and less affected regions (dentate gyrus of the hippocampus) of zQ175 mice and WT littermate controls. Note the increased number of C1q puncta in the striatum and motor cortex but not the dentate gyrus of the zQ175 mice. Scale bar = 5 μm. **(b)** Bar charts showing quantification of C1q puncta in these different brain regions at 7 mo, striatum n=4 WT and 4 zQ175 mice, motor cortex n=3 WT and 3 zQ175 mice and hippocampus n=3 WT and 4 zQ175 mice. Unpaired t-test for comparisons of WT and zQ175 in each brain region, striatum p=0.039, motor cortex p=0.017, hippocampus (DG) p=0.357. and 4 zQ175 mice. Unpaired t-test for comparisons of WT and zQ175 in each brain region, striatum p=0.039, motor cortex p=0.017, hippocampus (DG) p=0.357. **(c)** Bar charts showing quantification of C1q puncta in the dorsolateral striatum at different ages in zQ175 mice and WT littermates, 3 mo n=5 WT and 5 zQ175 mice, 7 mo n=4 WT and 4 zQ175 mice (same data represented in (b)) and 12 mo n=3 WT and 3 zQ175 mice. Unpaired t-test for comparisons of WT and zQ175 at each age, 3 mo p=0.007, 7 mo p=0.039 and 12 mo p=0.728. **(d)** Representative confocal images showing C3 staining in disease affected and less affected regions of zQ175 mice and WT littermate controls. Note that as with C1q, C3 puncta are increased in the striatum and motor cortex but not the hippocampus (DG). Scale bar = 5 μm. **(e)** Bar charts showing quantification of C3 puncta in the striatum, motor cortex and hippocampus (DG) of 7 mo zQ175 mice and WT littermate controls, striatum n=5 WT and 5 zQ175 mice, motor cortex n=4 WT and 3 zQ175 mice, hippocampus (DG) n=4 WT and 4 zQ175 mice. Unpaired t-test for comparisons of WT and zQ175 in each brain region, striatum p=<0.0001, motor cortex p=0.012 and hippocampus (DG) p=0.065. **(f)** Bar charts showing quantification of C3 puncta in the dorsolateral striatum at different ages in zQ175 mice and WT littermates, 3 mo n=5 WT and 5 zQ175 mice, 7 mo n=5 WT and 5 zQ175 mice (same data represented in **(e)**) and 12 mo n=4 WT and 4 zQ175 mice. Unpaired t-test for comparisons of WT and zQ175 at each age, 3 mo p=0.033, 7 mo p=<0.0001 and 12 mo p=0.047. **(g)** Orthogonal views of structured illumination images showing C3 and VGLUT1 staining or C3 and VGLUT2 staining in 3 mo zQ175 or WT dorsolateral striatum. Note the increased number of C3 puncta in the zQ175 tissue, which colocalize with VGLUT1 puncta in both the XY, YZ and XZ planes. Scale bar = 5 μm. Bar charts show Matlab quantification of colocalized C3 and VGLUT puncta from Imaris processed images. There is a significant increase in the % of VGLUT1 puncta colocalized with C3 in tissue from zQ175 mice but not in the % of VGLUT2 puncta colocalized with C3, n=4 WT and 4 zQ175 mice. Unpaired t-tests for comparisons of WT and zQ175, VGLUT1 and C3 p=0.002; VGLUT2 and C3 p=0.833. **(h) (i)** Representative confocal images of CR3 and Iba1 staining in the dorsal striatum of 3 mo zQ175 and WT mice. While levels of the microglia (and macrophage) marker Iba1 do not change (Extended data Fig. 5g), CR3 levels are increased in the processes of microglia from zQ175 mice. Scale bar = 10 μm. **(ii)** Structured illumination image showing that CR3 is localized to the processes of microglia in the dorsal striatum of zQ175 mice. Scale bar = 10 μm. Bar chart shows quantification of average CR3 fluorescence intensity in the dorsal striatum (as assessed from confocal images). There is a significant increase in CR3 levels in tissue from zQ175 mice, n=4 WT and 3 zQ175 mice. Unpaired t-test p=0.034. For bar charts, bars depict the mean. All error bars represent SEM. Stars depict level of significance with *=p<0.05, **p=<0.01 and ***p<0.0001

To assess whether C1q and C3 preferentially localize to specific subsets of synapses, we co-stained sections with antibodies to both complement proteins as well as markers of corticostriatal and thalamostriatal synapses using protocols similar to those outlined in previous studies^30,92^. We found that a significantly higher percentage of glutamatergic synapses co-localized with complement components C1q and C3 in the dorsolateral striatum but not in the hippocampus of 7 mo zQ175 and BACHD mice relative to WT littermate controls **(Extended data Fig. 3c**,**e)**. These data are consistent with unbiased proteomic assessments that have uncovered an enrichment of C1q in isolated synaptic fractions from the striatum of 6 mo zQ175 mice^93^. Strikingly, at 3 mo, when there is a selective loss of corticostriatal synapses in zQ175 mice, we observed an increased percentage of this synaptic population colocalizing with both C3 and C1q but no increased association of complement proteins with neighboring thalamostriatal synapses **(Fig. 3g and Extended data Fig. 3g)**. To demonstrate that this increase was not simply a result of a random association, we repeated this analysis with the C3 channel rotated 90 degrees, to simulate a chance level of colocalization, and saw that in zQ175 mice the association of complement proteins with corticostriatal markers (relative to that seen in WT littermates) was now significantly reduced **(Extended data Fig. 3f)**.

Complement receptor 3 (CR3) is expressed exclusively by myeloid cells in the brain and binds to various cleaved forms of complement component C3 that opsonize damaged cells or pathogens. This interaction prompts engulfment of the opsonized material by the myeloid cell. In the dorsal striatum of 3 mo zQ175 mice, CR3 levels were significantly increased and localized to the processes of microglia **(Fig. 3h)**. Collectively, these results show that complement proteins localize specifically to vulnerable corticostriatal synaptic connections before the onset of motor and cognitive deficits, and that the levels of their receptors are elevated on microglial cells, thereby demonstrating that they are present at the right time and place to mediate selective elimination of corticostriatal synapses.

### Microglia engulf corticostriatal projections and synaptic elements in HD mouse models

To test whether microglia mediate the engulfment of synaptic elements we first investigated whether there were region specific changes in microglia phenotypes consistent with the adoption of a more phagocytic state in HD models^94,95^. Using an established combination of markers that incorporates both changes in cell morphology (Iba1) and CD68 lysosomal protein levels^30,96^, we found a significant shift towards a more phagocytic microglial phenotype in the striatum and motor cortex (another disease affected brain region^97,98^) of 3 mo and 7 mo zQ175 mice (as has been observed in other HD models before the onset of motor phenotypes^51^) that was similar to what we saw in the postmortem HD tissue set. However, this was not seen in the less disease-affected hippocampus **(Extended data Fig. 4a**,**b)**. Similar changes in morphology were seen using Sholl analysis and a shift in phagocytic state was also observed in the striatum of BACHD mice **(Extended data Fig. 4c**,**d**,**e)**. To establish that the observed cells were not invading monocytes, we co-stained with Iba1 and the microglia-identity markers Tmem119 and P2ry12 **(Extended data Fig. 5a**,**b**,**c)**^99,100^. Like complement proteins, phagocytic microglia are thus present at the right time and place to play a role in synaptic elimination.

To directly test whether microglia engulfed more corticostriatal inputs in HD mice we labeled cortical neurons with a unilateral stereotactic injection of pAAV2-hsyn-EGFP into the motor cortex of postnatal day (PD) 1 zQ175 and wildtype (WT) littermate mice. Mice were subsequently allowed to recover and aged to 4 mo before being sacrificed (schematized in **Fig. 4a**). Microglial engulfment in the ipsilateral dorsal striatum was then assayed in fixed brain sections in which microglia were labeled with Iba1 (as described in Schafer et al., 2012 and Schafer et al., 2014^96,101^). We found that microglia in the striatum of 4 mo zQ175 mice contained a greater volume of the labeled corticostriatal projections, compared to WT littermate controls **(Fig. 4b, Extended data Fig. 5h)**. Importantly, no differences in the number of Iba1 stained cells or the transcripts for microglia-specific markers were detected in zQ175 striatum relative to WT littermates at 4 and 7 mo **(Extended data Fig. 5d**,**e**,**f**,**g)**, which could have confounded interpretation of this data.

**Figure 4:**
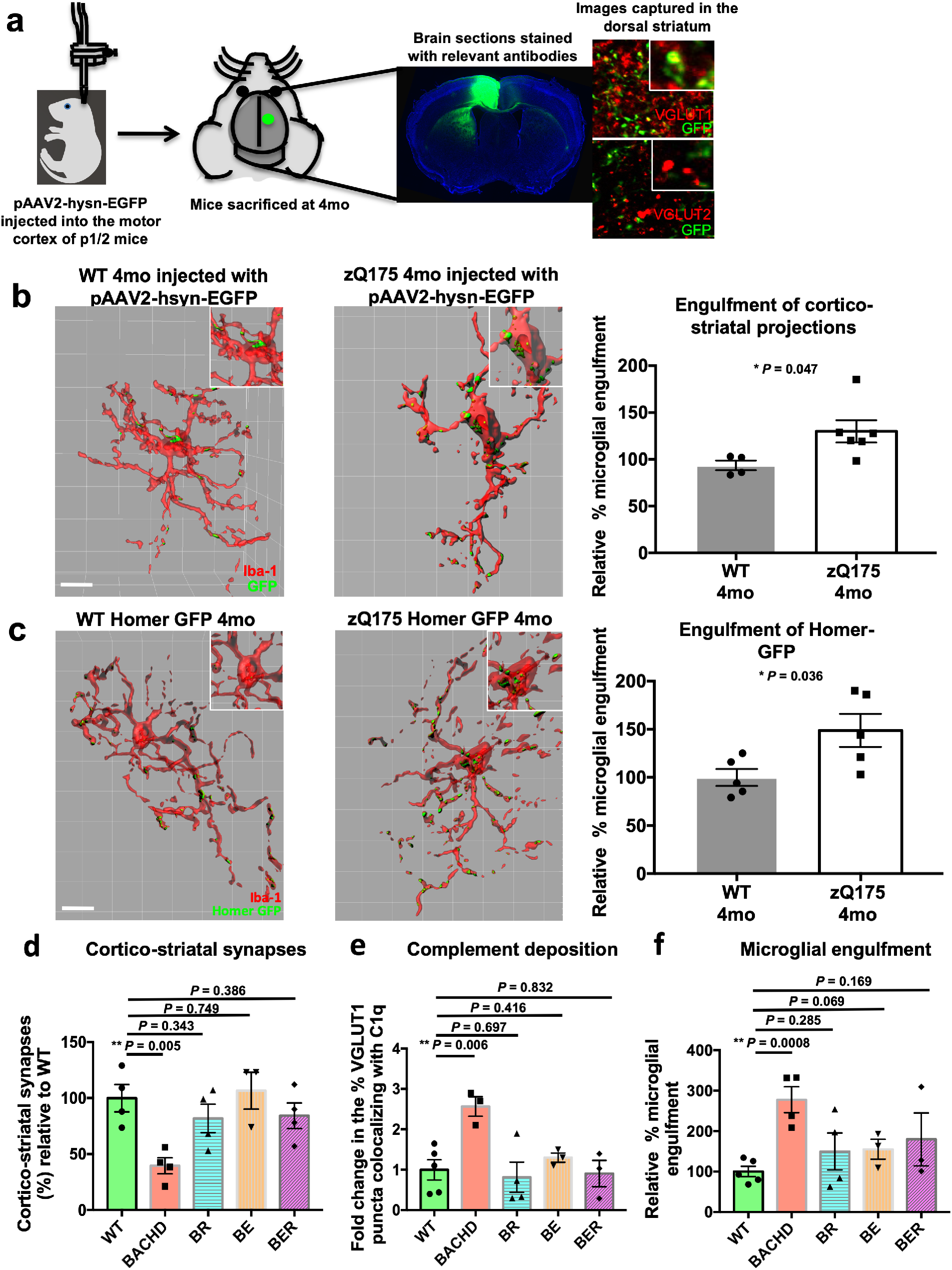
Microglia in the striatum of HD mice engulf more corticostriatal projections and synaptic material in a manner that is dependent on expression of neuronal mHTT. **(a)** Schematic showing how the corticostriatal projections were labelled with an injection of the pAAV2-hsyn-EGFP construct into the motor cortex of neonatal mice. The confocal image shows a coronal section from a 4 mo zQ175 injected mouse with strong GFP expression in the motor cortex and GFP labelled corticostriatal projections extending into the dorsal striatum. Panels to the right show colocalization of GFP signal in the dorsal striatum with VGLUT1, a marker of the corticostriatal synapse but not VGLUT2, a marker of the thalamostriatal synapse. **(b)** Representative surface rendered images of microglia (stained with Iba1) and engulfed GFP labeled corticostriatal projections in the dorsal striatum of 4 mo zQ175 and WT mice both of which received an injection of a pAAV2-hsyn-EGFP construct into their motor cortex at P1/2. Scale bar = 10 μm. Note the increased number of inputs inside the microglia from the zQ175 mouse (seen more clearly in the enlarged insets). Bar chart shows quantification of the relative % microglia engulfment (the volume of engulfed inputs expressed as a percentage of the total volume of the microglia) in 4 mo zQ175 mice relative to that seen in WT littermate controls. All engulfment values were normalized to the total number of inputs in the field, n=4 WT and 6 zQ175 mice. Unpaired t-test p=0.047. **(c)** Representative surface rendered images of microglia (stained with Iba1) and engulfed Homer-GFP in the dorsal striatum of 4 mo zQ175 Homer-GFP mice and WT Homer-GFP littermates. Scale bar = 10 μm. Note the increased amount of engulfed Homer-GFP inside the microglia from the zQ175 mice. Bar charts show quantification of the relative % microglia engulfment of Homer-GFP in 4 mo zQ175 Homer-GFP mice relative to that seen in WT Homer-GFP littermate controls, n=5 WT Homer-GFP and 5 zQ175 Homer-GFP mice. Unpaired t-test, p=0.036. **(d)** Bar chart showing quantification of the % of corticostriatal synapses in the dorsal striatum of BACHD, BR (deletion of mHTT in striatal neurons), BE (deletion of mHTT in cortical neurons) and BER (deletion of mHTT in striatal and cortical neurons) mice relative to those seen in WT littermates. Only BACHD mice expressing mHTT in both striatal and cortical neurons show a significant loss of this synaptic population, n=4 WT mice, 4 BACHD mice, 4 BR mice, 3 BE mice and 4 BER mice. Unpaired t-test with comparison to WT, BACHD p=0.005, BR p=0.343, BE p=0.749, BER p=0.386. **(e)** Bar chart shows the % of VGLUT1 puncta colocalized with C1q in the dorsal striatum of BACHD, BR, BE and BER mice and WT littermate controls. Again only BACHD mice expressing mHTT in both striatal and cortical neurons show a significant increase in complement deposition onto corticostriatal markers relative to that seen in WT mice, n=5 WT mice, 3 BACHD mice, 4 BR mice, 3 BE mice and 3 BER mice. Unpaired t-test with comparison to WT, BACHD p=0.006, BR p=0.697, BE p=0.416, BER p=0.832. **(f)** Quantification of the relative % microglial engulfment of corticostriatal projections in the dorsal striatum of BACHD, BR, BE, BER and WT mice. In line with the loss of corticostriatal synapses and increased complement deposition, BACHD mice show a significant increase in microglial engulfment relative to WT littermates, n=5 WT mice, 4 BACHD mice, 4 BR mice, 3 BE mice, and 3 BER mice. Unpaired t-test with comparison to WT, BACHD p=0.0008, BR p=0.285, BE 0.069, BER p=0.169. For bar charts, bars depict the mean. All error bars represent SEM. Stars depict level of significance with *=p<0.05, **p=<0.01 and ***p<0.0001

As this viral labeling method does not distinguish between the engulfment of presynaptic bouton structures, which connect cortical and striatal neurons, and axonal material, which might come from exosomes released from neurons or axosome shedding, we used a transgenic method to specifically assay engulfment of synaptic elements. zQ175 mice were crossed with mice in which the postsynaptic marker homer1c is fused to a GFP tag (hereafter referred to as homer-GFP)^102^. Engulfment analysis revealed that microglia in the dorsal striatum of zQ175 mice engulfed significantly more homer-GFP than those of their WT littermate controls **(Fig. 4c, Extended data Fig. 5i)**. Interestingly, in line with this result, a recent ultrastructural study found that microglia process interaction with synaptic clefts was also increased prior to the onset of motor deficits in a different HD mouse model^51^. To ensure that expression of the transgene had not affected pathological synapse loss, we quantified homer-GFP puncta and Homer1 immunoreactive puncta at 7 mo of age and saw the expected reduction of the markers’ levels in the zQ175 homer-GFP mice relative to homer-GFP littermates **(Extended data Fig. 6a**,**b)**. Taken together, these results show that microglia adopt a more phagocytic phenotype and accumulate significantly more engulfed synaptic material in the dorsal striatum of zQ175 mice.

Astrocyte dysfunction is linked to corticostriatal circuit abnormalities and motor and cognitive dysfunctions in HD mouse models and astrocytes can also phagocytose synaptic material during developmental refinement^103-106^. To test whether astrocytes engulf corticostriatal synapses in HD models, we stained sections from Homer-GFP and zQ175 Homer-GFP mice with the astrocyte marker S100β and performed engulfment analysis. We found that while astrocytes engulf synaptic elements in the striatum, the average volume of engulfed material was not altered in zQ175 mice relative to that seen in WT littermates **(Extended data Fig. 5k** and orthogonal views of engulfed material in **Extended data Fig. 5j)**, demonstrating that engulfment of synaptic material by astrocytes is unlikely to play a major role in the early synaptic loss we observe.

### Microglial engulfment and synapse loss are driven by mutant Huntingtin expression in both cortical neurons and MSN’s in the striatum

Mutant huntingtin (mHTT) is expressed ubiquitously, however previous work has demonstrated that specific HD pathologies can be driven by mHTT expression in different cell types^10,19,49,107-112^. To determine if the elimination of corticostriatal synapses requires mHTT in both cortical and striatal neurons, we quantified synapse density, complement deposition and microglial engulfment in BACHD mice in which mHTT had been selectively ablated from the cortex (using Emx1-cre, “BE” mice), striatum (using RGS9-cre “BR” mice) or both (using both Emx-1 and RGS9-cre, “BER” mice)^19^ **(Extended data Fig. 6g)**. We found that genetic deletion of mHTT from either the cortex, striatum or both completely rescued the loss of synapses normally seen in the BACHD model **(Fig. 4d)**. This is consistent with a previous study showing that both synaptic marker loss and corticostriatal synaptic transmission deficits are ameliorated by genetically deleting mHTT in either striatal or cortical neurons^19^. In addition, deposition of complement component C1q, as well as microglial engulfment, was reduced to WT levels **(Fig. 4e**,**f)**. Thus, mHTT expression in both MSNs and cortical neurons is required to initiate the complement deposition, microglial engulfment of synaptic elements, and synapse loss we observe in the BACHD model.

### Early loss of corticostriatal synapses is rescued by inhibiting activation of the classical complement pathway or genetically ablating microglial complement receptor 3

To test whether increased synaptic complement deposition and engulfment of corticostriatal inputs by microglia mediates the selective elimination of these synapses in zQ175 mice, we blocked these processes using several approaches. To prevent microglial interaction with bound complement proteins, we genetically ablated CR3 in zQ175 mice **(Extended data Fig. 8a)**. This increased the number of corticostriatal synapses present in the dorsolateral striatum by 60% relative to zQ175 littermates expressing CR3 (Fig. 5g). Importantly, this increase was not due to persistent effects related to differences in developmental pruning, as has been observed previously at the retinogeniculate synapse in CR3KO mice, or other aspects of altered biology, as CR3 ablation on a WT background did not alter corticostriatal synapse number at 4mo **(Extended data Fig. 8b)**.

**Figure 5:**
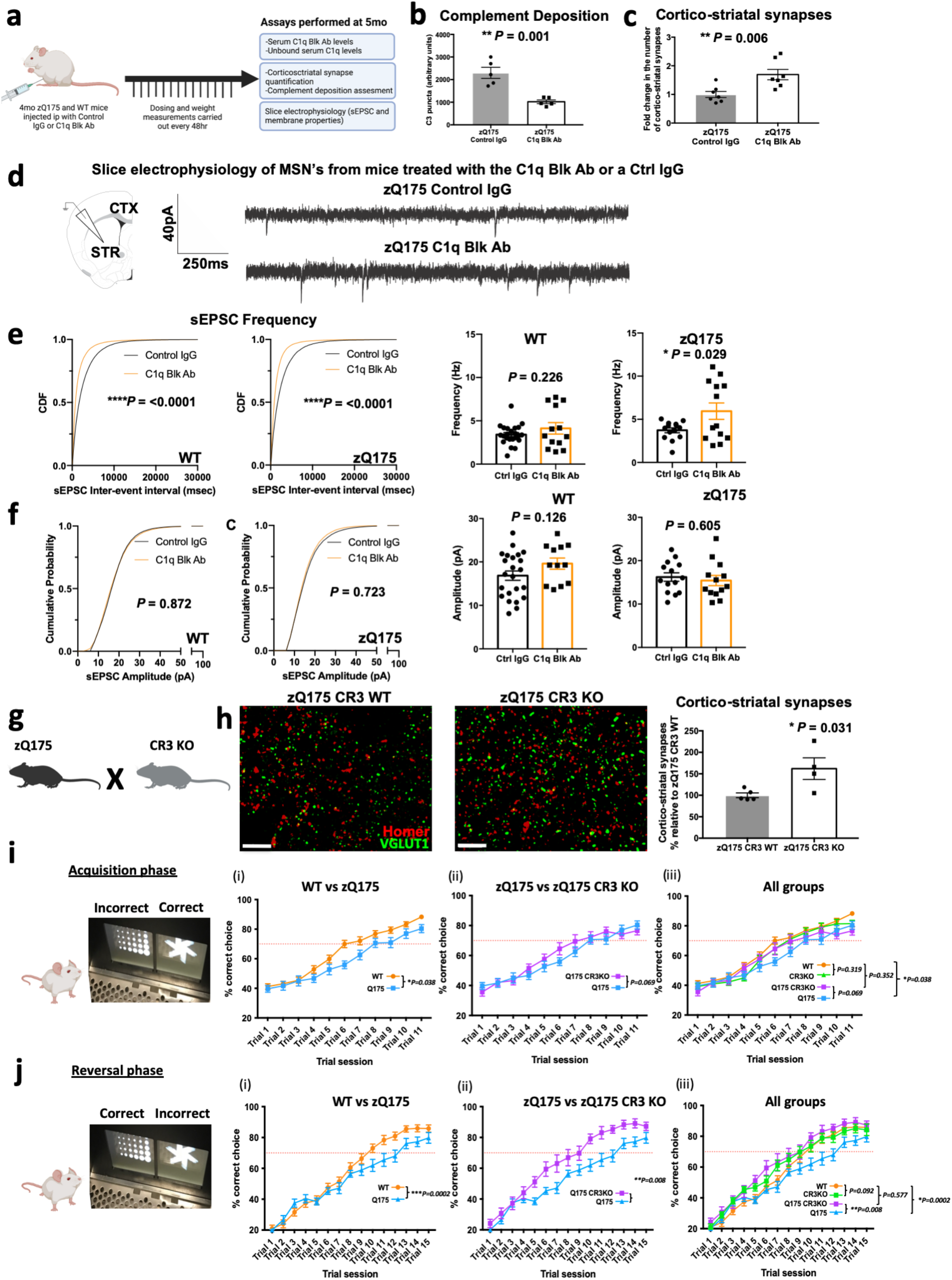
Blocking complement deposition or microglial recognition of complement opsonized structures with either genetic or antibody mediated strategies can reduce synaptic loss and prevent the development of cognitive deficits in HD mice. **(a)** Schematic showing the experimental paradigm in which 4mo zQ175 mice and WT littermates were treated with the C1q Blk Ab or a control IgG for 1 mo before being sacrificed. Relevant tissues and fluids were collected at 5mo for interrogation (see methods). **(b)** Bar chart shows quantification of C3 puncta in the dorsolateral striatum of zQ175 mice treated with the C1q function blocking antibody or a control IgG. Note that in zQ175 mice treated with control IgG there is increased deposition of complement C3 protein relative to that seen in zQ175 mice treated with the C1q function blocking antibody, n= 5 zQ175 mice treated with control IgG and 5 zQ175 mice treated with C1q function blocking antibody. Unpaired t-test p=0.001. **(c)** Bar charts show quantification of colocalized pre and post synaptic puncta denoting corticostriatal synapses in the dorsolateral striatum of zQ175 mice treated with the C1q function blocking antibody or a control IgG. There are significantly more corticostriatal synapses in the dorsolateral striatum of the zQ175 mice treated with the C1q function blocking antibody relative to those treated with the control IgG, n=7 zQ175 mice treated with control IgG and 6 zQ175 mice with the C1q function blocking antibody. Unpaired t-test p= 0.006. **(d)** Diagram depicting the strategy for carrying out electrophysiology recordings from coronal sections of treated mice and representative traces of sEPCS recorded from MSN’s in striatal slices from 5 mo zQ175 mice which had been treated for 1 mo with control IgG or the C1q function blocking antibody. **(e)** Cumulative distribution plots of interspike intervals (ISI) obtained from whole-cell voltage clamp recordings of medium spiny neuron spontaneous excitatory postsynaptic currents. Recordings were carried out in slices from WT and zQ175 mice after a 1 mo treatment with control IgG or the C1q function blocking antibody (Black = Control IgG, Orange = C1q function blocking antibody). Bar charts show average frequency (Hz) per cell recorded across conditions. n=13-23 cells per condition from 4-7 animals. Kolmogorov-Smirnov test for the cumulative distribution plots for both WT and zQ175 p=<0.0001; Unpaired t test for the bar charts, for WT p=0.226; for zQ175 p=0.029. **(f)** Cumulative distribution plots of amplitude obtained from whole-cell voltage clamp recordings of medium spiny neuron spontaneous excitatory postsynaptic currents. Recordings were carried out in slices from WT and zQ175 mice after a 1 mo treatment with control IgG or the C1q function blocking antibody (Black = Control IgG, Orange = C1q function blocking antibody). Bar charts show average amplitude (pA) per cell recorded across conditions. n=13-23 cells per condition from 4-7 animals. Kolmogorov-Smirnov test for the cumulative distribution plots, for WT p=0.872; for zQ175 p=0.723; Unpaired t test for the bar charts, for WT p=0.126; for zQ175 p=0.605. **(g)** Diagram showing the two transgenic mouse lines that were crossed together to generate the genotypes employed in subsequent assessments of corticostriatal synapse number as well as visual discrimination learning and reversal performance. **(h)** Representative structured illumination images of Homer1 and VGLUT1 staining in the dorsolateral striatum of 4 mo zQ175 CR3 WT and zQ175 CR3 KO mice. Note the increased number of both pre and postsynaptic puncta in the zQ175 CR3 KO. Scale bar = 5 μm. Bar chart shows Imaris and Matlab quantification of colocalized puncta denoting corticostriatal synapses. There are significantly more corticostriatal synapses in the zQ175 CR3 KO’s than in the zQ175 littermates expressing CR3, n=5 zQ175 CR3 WT mice and 4 zQ175 CR3 KO mice. Unpaired t-test p=0.031. **(i)** Following completion of shaping tasks (see methods), the visual discrimination performance of 4 mo WT, zQ175, CR3 KO and zQ175 CR3 KO mice was assessed using the Bussey-Sakisda operant touchscreen platform. Line charts show performance over 11 trial sessions (60 trials per session) of (i) WT vs zQ175 mice, (ii) zQ175 vs zQ175 CR3 KO mice and (iii) all groups. Note that WT mice reached the criterion of 70% correct choice selection in a session of 60 trials more rapidly than their zQ175 littermates and that this impairment in visual learning is prevented in zQ175 mice in which complement receptor 3 has been genetically ablated, n= 29 WT mice, 18 zQ175 mice, 24 CR3KO mice and 13 zQ175 CR3KO mice. Two way anova: for WT vs zQ175 p=0.038 for the combination of genotype x trial session as a significant source of variation and p=0.01 for genotype as a significant source of variation; for zQ175 vs zQ175 CR3KO p=0.069 for the combination of genotype x trial session as a significant source of variation and p=0.568 for genotype as a significant source of variation; for WT vs zQ175 CR3KO p=0.352 for the combination of genotype and trial as a significant source of variation and p=0.068 for genotype as a significant source of variation; for WT vs CR3 KO p=0.319 for the combination of genotype x trial session as a significant source of variation and p=0.593 for genotype as a significant source of variation. **(j)** Following completion of the acquisition phase the visual presentations were switched so that the visual stimuli previously associated with reward provision was now the incorrect choice. Performance of WT, zQ175, CR3 KO and zQ175 CR3 KO mice in this reversal phase of the task was then assessed using the Bussey-Sakisda operant touchscreen platform. Line charts show performance over 15 trial sessions (60 trials per session) of (i) WT vs zQ175 mice, (ii) zQ175 vs zQ175 CR3 KO mice and (iii) all groups. Note that WT mice again reached the criterion of 70% correct choice selection in a session of 60 trials more rapidly than their zQ175 littermates and that this impairment in cognitive flexibility is abrogated in zQ175 mice in which complement receptor 3 has been genetically ablated, n= 29 WT mice, 18 zQ175 mice, 24 CR3KO mice and 13 zQ175 CR3KO mice. Two way anova: for WT vs zQ175 p=0.0002 for the combination of genotype x trial session as a significant source of variation and p=0.080 for genotype as a significant source of variation; for zQ175 vs zQ175 CR3KO p=0.008 for the combination of genotype x trial session as a significant source of variation and p=0.003 for genotype as a significant source of variation; for WT vs zQ175 CR3KO p=0.577 for the combination of genotype and trial as a significant source of variation and p=0.072 for genotype as a significant source of variation; for WT vs CR3 KO p=0.092 for the combination of genotype x trial session as a significant source of variation and p=0.304 for genotype as a significant source of variation. For bar charts, bars depict the mean and all error bars represent SEM. Stars depict level of significance with *=p<0.05, **p=<0.01 and ***p<0.0001.

To reduce complement deposition, we injected a monoclonal C1q function blocking antibody, also referred to as ANX-M1 in some publications (see methods), intraperitoneally (IP) into 4 mo zQ175 mice and WT littermates over a 1 mo period **(Fig. 5a)**. ANX-M1 binds to an epitope in the globular head domains of C1q, preventing C1q binding to its substrates and downstream activation of the proteases C1r and C1s^113-115^, which results in suppression of the classical complement cascade activation and reduced cleavage and subsequent deposition of C3^30,34,116^. ELISA assays confirmed that treated mice had high levels of ANX-M1 and significantly less unbound C1q in their serum relative to mice treated with a control IgG **(Extended data Fig. 7c**,**d)**. FITC-tagged versions of ANX-M1 could also be detected in the neuropil of these mice, demonstrating that it can cross the blood brain barrier **(Extended data Fig. 7p)**, a finding that is consistent with other studies in which rodents have received peripheral administration of antibodies^117-119^.

Treatment with the C1q function blocking antibody reduced complement component C3 deposition in the dorsolateral striatum of zQ175 mice by approximately 60% and increased the number of corticostriatal synapses by 70% relative to treatment with a control IgG **(Fig. 5b**,**c)**, but had no effect on baseline complement deposition or corticostriatal synapse numbers in WT mice **(Extended data Fig. 7g**,**h)**. It was also well tolerated, with no change in body weight during the course of the study **(Extended data Fig. 7e**,**f)**.

Blocking complement deposition in zQ175 mice led to a greater number of corticostriatal synaptic structures. In order to test whether these synapses were functional, we performed *ex vivo* electrophysiology using striatal brain slices from mice treated with the C1q function blocking antibody or the control IgG. In line with the reduced number of corticostriatal synapses we observe at 3mo and in keeping with what others have reported at older ages with this HD line^72,74,77,78^, untreated 5mo zQ175 mice showed a reduction in spontaneous excitatory postsynaptic current (sEPSC) frequency (as denoted by cumulative distribution plots of sEPSC interevent interval values; mean values per cell showed a reduction but this was not statistically significant with the current sample size), reflecting a decline in either the presynaptic release probability or the number of presynaptic inputs onto the cell in the zQ175 mice when compared to their WT littermates **(Extended data Fig. 7i)**. We also saw the expected increase in input resistance, a membrane property which is thought to reflect a state in which the cell responds more significantly to a given stimulus and subtle reductions in amplitude which reflect changes in the density of postsynaptic receptors at an individual synapse **(Extended data Fig. 7j**,**k**,**l)**.

Treatment with the C1q function blocking antibody significantly shifted the sEPSC frequency distribution for both zQ175 and WT controls and in zQ175 mice, increased the mean frequency per cell relative to that seen in mice treated with control IgG. These data demonstrate that cells from mice treated with the blocking antibody had a greater number of presynaptic inputs or a greater release probability **(Fig. 5d**,**e and Extended data Fig. 7m)**. In contrast, no treatment induced changes were observed in the amplitude response or the intrinsic membrane properties of these cells, such as capacitance and input resistance **(Fig. 5f and Extended data Fig. 7n**,**o)**. Taken together, these results demonstrate that blocking complement activity in zQ175 mice leads to more functional excitatory synapses on MSN’s and increased excitatory input to the striatum, thereby showing that strategies that target complement in this way can reduce the loss of corticostriatal synapses in HD mice.

### The development of early cognitive deficits is prevented by genetic ablation of microglial complement receptor 3

Cognitive deficits are a key clinical hallmark of HD and impaired performance in tasks related to visual discrimination and cognitive flexibility are some of the earliest quantifiable changes observed in premanifest HD patients, occurring up to 20 years prior to predicted onset of manifest disease and in the absence of detectable motor phenotypes^120-122^. Impairments in learning and memory tasks have also been observed in the zQ175 HD mouse model but these assessments have only been carried out in older mice after the development of striatal atrophy, significant transcriptional dysregulation and the appearance of a range of other pathological hallmarks^17,71-73,123-126^

To determine whether cognitive deficits can be observed at an earlier stage of disease progression in zQ175 mice, at time points corresponding to the loss of corticostriatal synapses, we used an operant touchscreen platform to monitor performance in a task of visual object discrimination, in which mice had to learn which image presentation was associated with the dispensing of a liquid reward. After completion of this task an assessment of cognitive flexibility was performed by reversing the object display associated with reward such that the previously incorrect choice now yielded a reward response and vice versa. Assessments in these tasks showed that 4mo zQ175 mice displayed a significantly impaired performance in visual discrimination learning compared to WT littermates but they were eventually able to achieve a threshold of 70% correct choice over 60 trial sessions on 3 consecutive trial days **(Fig. 5i(i))**. When the reward association was subsequently reversed it again took the zQ175 mice much longer to reach a 70% correct choice threshold indicating an impairment in cognitive flexibility **(Fig. 5j(i))**. These deficits were unlikely to be driven by differences in visual acuity, motor performance or motivation to complete the task as optomotor measurements, the speed and number of trials completed and performance in progressive ratio tasks showed no genotype dependent differences **(Extended data Fig. 8c**,**d**,**e**,**f**,**g**,**h**,**i**,**j)**.

To test whether strategies that block complement and microglia mediated synapse elimination rescue the cognitive flexibility impairments observed in the zQ175 mice, we also assessed the performance of zQ175 CR3KO mice in these tasks. Genetic ablation of CR3 prevented the impaired rate of visual discrimination learning observed in zQ175 mice with performance of zQ175 CR3KO mice not being significantly different to that of WT and CR3KO littermate controls **(Fig. 5i(ii and iii))**. Furthermore, in the reversal phase of the task zQ175 CR3KO mice showed significantly improved performance when contrasted to zQ175 littermates taking 240 fewer trials to achieve the 70% correct threshold **(Fig. 5j (ii and iii)**.

Taken together these results show that zQ175 mice display deficits related to visual processing and cognitive flexibility at 4mo of age at a time point where we also observe loss of corticostriatal synapses. These deficits do not appear to be driven by impairments in visual acuity, motor performance or motivation to complete the task. In addition, adopting strategies that reduce complement mediated synaptic elimination prevents the appearance of these deficits and returns performance to WT levels.

### Complement protein levels in the CSF are correlated with measures of disease burden in premanifest HD patients

To determine if changes in complement protein levels or activity can be detected in the cerebral spinal fluid (CSF) of premanifest HD patients and if so whether these changes are associated with established measures of disease burden we developed quantitative assays to measure complement cascade proteins and their activation fragments. CSF is a unique biological fluid containing proteins with enriched expression in the brain and spinal cord^127-130^ and modifications to its proteome reflect transcriptional changes in disease-associated regions of the Huntington’s disease brain^131^. It is also possible to safely obtain CSF from patients before the manifestation of clinical symptoms, enabling the observation of early pathological changes^132,133^.

Analysis of CSF samples from the HDClarity study group (see methods and Supplemental table 1 for clinical details) with ELISA assays that detect C3 and iC3b showed that both proteins were altered with disease stage (as defined by both clinical assessment and burden of pathology score (see methods)) in HD patients **(Fig. 6a**,**b)**. Levels of C3 and iC3b were both significantly increased in patients with early manifest HD relative to those with early premanifest HD. Importantly, for iC3b, this increase was also seen between early premanifest HD patients and late premanifest HD patients, a transition that is associated with reductions in brain and striatal volume, white matter microstructure damage, reduced functional activity of the basal ganglia while performing tasks, and multiple subtle differences in motor and cognitive performance as well as differences in neuropsychiatric assessments (see methods)^1,26,132,134,135^. This demonstrates that changes in the abundance of iC3b occur before the onset of motor symptoms.

**Figure 6:**
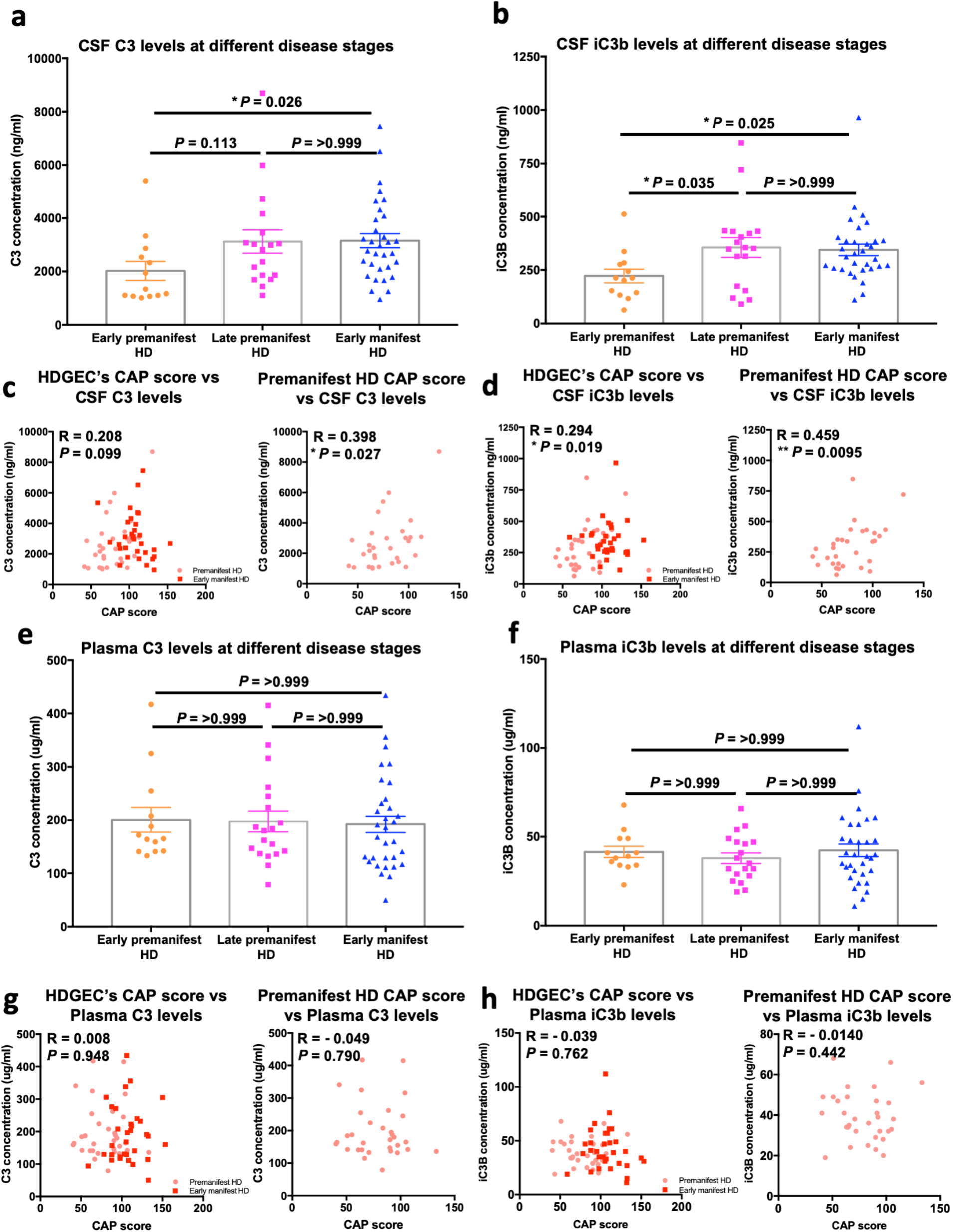
Association between disease stage and complement component levels in the CSF and plasma of Huntington’s disease patients. **(a)** Bar chart showing the concentration of complement component C3 in CSF samples from early premanifest HD patients (see methods for inclusion criteria), late premanifest HD patients (see methods for inclusion criteria) and early manifest HD patients (see methods for inclusion criteria) recruited into the HDClarity study. Each dot represents a sample from a separate individual and the bar denotes the mean for each group n=13 early premanifest HD, n=18 late premanifest HD and n=32 early manifest HD. Kurkasl-Wallis test (non-parametric ANOVA) p=0.028 with early premanifest versus late premanifest HD p=0.113, late premanifest versus early manifest HD p = >0.999 and early premanifest versus early manifest HD p=0.026 via Dunn’s multiple comparison test. **(b)** Bar chart showing the concentration of iC3b, the cleaved component of C3 that is formed following activation of the complement cascade, in the same CSF samples assayed in **(a)** n=13 early premanifest HD, n=18 late premanifest HD and n=32 early manifest HD. Kurkasl-Wallis test (non-parametric ANOVA) p=0.017 with early premanifest HD versus late premanifest HD p=0.035, late premanifest HD versus early manifest HD p =>0.999 and early premanifest HD versus early manifest HD p=0.025 via Dunn’s multiple comparison test **(c)** Graphs showing the association between CAP score and CSF C3 concentration for all samples from Huntington’s Disease gene expansion carriers (HDGECs) as well as those just from premanifest HD patients recruited into the HDClarity study. Each dot represents a sample from a separate individual n=63 HDGEC’s and n=31 premanifest HD. Spearman r for HDGEC’s p=0.099; for premanifest HD p=0.027. **(d)** Graphs showing the association between CAP score and CSF iC3b concentration for all samples from Huntington’s Disease gene expansion carriers (HDGECs) as well as those just from premanifest HD patients recruited into the HDClarity study. Each dot represents a sample from a separate individual n=63 HDGEC’s and n=31 premanifest HD. Spearman r for HDGEC’s p=0.019; for premanifest HD p=0.0095. **(e)** Bar chart showing the concentration of complement component C3 in plasma samples from early premanifest HD patients (see methods for inclusion criteria), late premanifest HD patients (see methods for inclusion criteria) and early manifest HD patients (see methods for inclusion criteria) recruited into the HDClarity study. Each dot represents a sample from a separate individual and the bar denotes the mean for each group n=13 early premanifest HD, n=19 late premanifest HD and n=32 early manifest HD. Kurkasl-Wallis test (non-parametric ANOVA) p=0.797 with early premanifest versus late premanifest HD p=>0.999, late-premanifest versus early manifest HD p=>0.999 and early premanifest versus early manifest HD p=>0.999 via Dunn’s multiple comparison test. **(f)** Bar chart showing the concentration of iC3b, the cleaved component of C3 that is formed following activation of the complement cascade, in the same plasma samples assayed in **(e)** n=13 early premanifest HD, n=19 late premanifest HD and n=32 early manifest HD. Kurkasl-Wallis test (non-parametric ANOVA) p=0.609 with early premanifest HD versus late premanifest HD p=>0.999, late premanifest HD versus early manifest HD p=>0.999 and early premanifest HD versus early manifest HD p=>0.999 via Dunn’s multiple comparison test **(g)** Graphs showing the association between CAP score and plasma C3 concentration for all samples from Huntington’s Disease gene expansion carriers (HDGECs) as well as those just from premanifest HD patients recruited into the HDClarity study. Each dot represents a sample from a separate individual n=64 HDGEC’s and n=32 premanifest HD. Spearman r for HDGEC’s p=0.948; for premanifest HD p=0.790. **(h)** Graphs showing the association between CAP score and plasma iC3b concentration for all samples from Huntington’s Disease gene expansion carriers (HDGECs) as well as those just from premanifest HD patients recruited into the HDClarity study. Each dot represents a sample from a separate individual n=64 HDGEC’s and n=32 premanifest HD. Spearman r for HDGEC’s p=0.762; for premanifest HD p=0.442. For bar charts, bars depict the mean. All error bars represent SEM. Stars depict level of significance with *=p<0.05, **p=<0.01 and ***p<0.0001

These findings were further confirmed by looking at the association between CSF concentrations of C3 and iC3b and the CAP score of HD patients. CAP score is thought to be a measure of disease burden and is derived from a mathematical formula that incorporates both the length of the individuals’ CAG expansions and their current age. Changes in this metric correlate with reductions in striatal volume, damage to white matter microstructure and, most importantly, the development of motor impairments (see methods)^1,136-143^ **(Extended data Fig. 9e**,**f**,**g)**. For both C3 and iC3b, there was a significant positive correlation between their CSF concentrations and CAP score in premanifest HD patients – a relationship that was maintained for iC3b after incorporating samples from early manifest HD patients **(Fig. 6c**,**d)**.

Both age and sex are potentially confounding variables. In line with previous studies, Spearman rank correlation models showed that, in our sample cohort, CSF C3 levels were elevated with aging in control (clinically normal; see methods and **Supplemental table 1**) individuals **(Extended data Fig. 9h)**^144-147^. iC3b levels, in contrast, were not **(Extended data Fig. 9i)**. To control for normal aging in our study, we adjusted the measured CSF concentrations of C3 and iC3b using a formula that incorporates the regression coefficient of aging in control individuals and the mean age of the control population in our sample cohort (see methods). While the significant association with CAP score in premanifest HD patients did not survive this adjustment, the difference in CSF iC3b levels between early and late premanifest HD patients was maintained **(Extended data Fig. 9k**,**l**,**m**,**n)**. However, the limited age range of the control samples and the fact that younger ages were significantly underrepresented **(Supplemental table 1)** means that further analysis of larger and more balanced cohorts of CSF samples from control individuals will be required to gain a better understanding of the effect of normal aging on CSF levels of complement proteins. The limited age range of the control samples is also the reason that no direct comparison of CSF complement levels was carried out between the control and HD patient cohorts. Gender also affects CSF C3 levels, with males having significantly more C3 in the HD patient samples; however, given the equal distribution of genders in all three patient groups, this did not alter the complement protein changes we observed with disease stage **(Extended data Fig. 10a**,**b**,**c)**.

A positive correlation between CSF iC3b levels and CAP score was also observed in a separate smaller cohort of CSF samples collected at the University of Washington under slightly different conditions (see methods) **(Extended data Fig. 11b)**, further demonstrating that iC3b increases with HD disease progression. We obtained longitudinal measurements from three premanifest HD patients in this cohort 1.5 years after the first collection **(Supplemental table 1)**, and in all three cases, the patients had increased iC3b levels in their CSF **(Extended data Fig. 11g**,**h**,**i)**, providing additional evidence that CSF iC3b concentration increases with disease progression. In contrast, C3 levels did not change in this CSF cohort, likely due to the smaller sample size and the weaker relationship between CAP score and C3 levels **(Fig.6c and Extended data Fig. 11a)**.

C3 and iC3b concentrations were unaltered with disease stage in paired plasma samples from the same patients and there was no correlation between plasma and CSF levels, demonstrating that complement changes in the CSF are localized and are not mirrored by systemic changes in other body fluids **(Fig. 6e**,**f, Extended data Fig. 10g**,**h**,**i)**. Plasma levels of C3 and iC3b also showed no correlation with CAP score and no significant difference in levels between genders **(Fig. 6g**,**h and Extended data Fig. 10d**,**e**,**f)**. This finding differs from what has previously been reported for downstream complement components such as C7 which have been found to be elevated in plasma samples from manifest HD patients relative to those from patients in the premanifest state^148^.

Interestingly, despite C1q being the initiating protein in the classical complement cascade, and thus acting to promote the activation/cleavage of C3 to iC3b, its levels were not associated with disease stage in the CSF of HD patients **(Extended data Fig. 9a)**. There was no correlation of CSF levels of C1q with CAP score, and unlike what was observed with C3, CSF levels of C1q showed no significant correlation with age in control individuals **(Extended data Fig. 9b**,**j)**. Its levels were also not different between males and females **(Extended data Fig. 10c)**. Furthermore, as seen with C3 and iC3b, plasma levels of C1q were unchanged with disease progression in samples from HD patients and not different between males and females **(Extended data Fig. 9c**,**d and Extended data Fig. 10f)**. This result is not unexpected because C1q, like C3, circulates at high levels and is present within the CNS even in individuals with no known evidence of injury or clinically defined disease^129,149^. Activation of C1q is also not regulated by its expression (although changes in expression can obviously augment the process), but rather by its binding to a target surface, in this case potentially through the recognition of changes occurring on the surface of synapses and other structures in the brain^113,150-152^. While local cellular expression of both C1q and C3 can increase at sites of pathology, relative blood or CSF levels may not change depending on the rate of local release and consumption. In contrast, the relative increase of iC3b we observe within the CSF with progression to more advanced disease stage is consistent with local activation of the complement cascade, reflecting ongoing activation and degradation of C3 within the CNS.

Taken together these results show that there are early changes in complement biology occurring specifically in the CSF of premanifest HD patients. Further work will be required to determine the factors prompting these changes and whether they correlate with clinical measures of disease progression as well as other known imaging and fluid markers.

## Discussion

Our study provides strong preclinical evidence that strategies to block complement-mediated microglial engulfment of synapses in HD may provide a therapeutic benefit in preventing the loss of corticostriatal synaptic connectivity and reducing cognitive behavioral deficits. We find that in two well-established HD mouse models expressing full-length mHTT, corticostriatal synapses are selectively lost at a very early disease stage, prior to the onset of robust motor and cognitive deficits. We provide multiple lines of evidence that complement and microglia mediate this corticostriatal synaptic loss, with C1q and C3 specifically associating with this synaptic population relatively early in disease progression and microglia engulfing more cortical inputs. Inhibition of the classical complement pathway with a therapeutically relevant C1q function blocking antibody and genetic ablation of complement receptor CR3, a gene exclusively almost exclusively expressed by microglia in the brain, both reduce early loss of coriticostriatal synapses in HD mouse models. In addition, ablation of microglial CR3^91^, prevented the development of deficits in visual discrimination learning and rescued impairments in cognitive flexibility observed at this age. While recent studies have demonstrated that microglia and complement coordinate to eliminate synapses in certain pathological contexts^30-32,34,36,153,154^, our study is the first to show that: (i) these mechanisms can selectively target synapses in a disease relevant neuronal circuit and (ii) that strategies that can block this process and preserve this synaptic population can also prevent the development of cognitive deficits.

An important question raised by these findings is what mechanisms mediate the selective association of complement proteins with corticostriatal synapses. Cortical and thalamic inputs form differing proportions of their structural connections onto different parts of striatal neuron dendrites^67-69^, and the receptor subunit composition and electrical properties of these two synaptic populations are distinct^155^. These structural differences may make thalamostriatal synapses more stable and less accessible for microglial engulfment. Equally, the different molecular compositions and pattern of electrical activity at these two synaptic populations could affect their relative levels of putative complement regulators such as CSMD1 and CLU, both of which have been found to be downregulated at the protein level in mouse models of HD^17^. CD47, a protective signal that inhibits microglial engulfment and whose localization at synapses is regulated by levels of neuronal activity^17,156^ may also be differentially distributed between these two synaptic populations. Finally, differences in astrocyte proximity to synaptic populations at early stages of the disease may also play a role in corticostriatal vulnerability. A recent study using a FRET-based proximity detector found that astrocyte association with corticostriatal inputs, but not with thalamostriatal inputs, is reduced in HD mice before synapse loss occurs^87^. Future studies to further delineate whether specific populations of corticostriatal synapses, such as those arising from intratelencephalic neuron or pyramidal neuron inputs, or those that form connections onto different types of medium spiny neuron, such as the D1 and D2 MSN’s, have a greater propensity for complement association and elimination at early stages of the disease will also be valuable in this regard.

Mutant HTT is expressed ubiquitously, but it elicits selective pathology in specific brain regions and synaptic circuits and causes cell-autonomous and non-cell autonomous pathological effects^10,19,49,107-110,112^. Our data suggest that expression of mHTT in both the presynaptic cortical and postsynaptic striatal neurons is required to trigger synaptic elimination and that mHTT expression in microglia is not sufficient to induce this process. Whether mHTT expression in microglia acts to potentiate this pathology, however, remains an open question. A recent study found that a 50% reduction of mHTT in microglia normalizes the expression of inflammatory cytokines but has no effect on the deterioration of motor performance or brain atrophy in the BACHD model^109^. While this doesn’t rule out the possibility that pathological events occurring early in the disease are impacted by reduced microglial mHTT levels, it does suggest that disease progression is not abrogated with this approach.

While our data shows that neuronal mHTT expression initiates complement-mediated synaptic elimination the mechanisms underlying this process remain undetermined. The corticostriatal pathway exhibits aberrant neuronal activity early in disease progression, before the onset of motor or cognitive deficits^21,22^, so it could be that mHTT expression in neurons leads to changes in corticostriatal synaptic transmission and neuronal firing that are detected by microglia. During normal developmental critical periods, microglia engulf retinal inputs in the visual thalamus in an activity-dependent manner^96^. Thus we could hypothesize that in disease contexts, aberrant neuronal activity, arising from mHTT expression in both cortical and striatal neurons^19,157^, promotes the deposition of a pro-engulfment molecular cue, such as complement, or the removal or redistribution of protective molecules, such as CD47, which then prompts microglial engulfment and removal of the synapse^96,156^. Understanding the link between mHTT-induced abnormalities in neuronal interactions and complement-mediated microglial engulfment will be crucial in shedding light on the upstream drivers of this synaptic elimination mechanism.

We found that strategies to block complement-mediated microglial engulfment of synapses in HD mouse models prevent at least some of the synaptic loss observed as well as inhibiting the appearance of early deficits in tasks of visual discrimination and cognitive flexibility. These results are consistent with prior studies where less specific methods were employed to demonstrate a selective role for this brain region in the performance of similar tasks^158^ as well as previous work showing that a none specific depletion of microglia can improve performance in some motor and cognitive tests^159^. However, it remains to be determined whether preservation of these synaptic structures has an impact on disease progression as well as the development of further deficits in motor and cognitive ability that have been observed previously in this HD model. It will thus be important to determine whether the synapses that remain after intervention are still functional and if there is a critical period after which it is not possible to intervene. Studies on developmental synaptic pruning have shown that complement knockout mice have excessive numbers of functional synapses in the retinogeniculate system^92^, suggesting that preserved corticostriatal synapses will still be able to transmit signals from cortical neurons to MSNs, and indeed, our slice electrophysiology data show that striatal neurons in HD mice treated with the C1q function blocking antibody have a greater number of excitatory inputs. However, a more in-depth analysis will be required to determine if these synapses still respond to different patterns of neuronal input in the same way and whether they are maintained as the disease progresses. An interesting finding from our slice electrophysiology data was the observation that the mean frequency data from the C1q function blocking antibody treated mice displayed a bimodal distribution perhaps suggesting that different medium spiny neuron populations (such as D1 and D2 MSN’s) react differently to the C1q function blocking antibody but further studies will be required to determine this. Notwithstanding the converging evidence of synaptic and behavioral benefit from genetic and pharmacological strategies to inhibit complement mediated synapse elimination in HD models, we are cognizant that our study is focused on relatively early stages of the disease. Thus, future investigations of the kinds described above are needed in these preclinical models to determine the optimal therapeutic window for intervention, which will be important for translating these findings into the clinic.

We provide several lines of evidence that complement and microglia mediated synaptic elimination mechanisms may also be operating in HD patients. In addition, in premanifest and early manifest HD patients C3 and iC3b levels in the CSF correlate with an established predictor of both pathological severity and time to clinically defined disease onset. Future longitudinal studies will be required to see whether changes in complement levels in HD patient CSF samples correlate with other aspects of clinical disease progression such as changes in the well-established motor and cognitive tests that comprise the Unified Huntington’s disease Rating Scale (UHDRS). If they do, even after accounting for known predictors of disease progression such as age and CAG repeat length, they could be used alongside other fluid and imaging markers^133,160-162^ as determinates of disease state enabling progression to be monitored and potentially predicting the development of clinical symptoms. Further, they would serve as important biomarkers of target engagement and pharmacodynamic impact for developing anti-complement therapies in the clinic in HD patients. To this end a Phase 2 study of an initial anti-C1q therapy (ANX005) is currently ongoing in HD patients (clinical trials.gov NCT04514367).

Our study provides the first in vivo evidence to validate targeting complement component C1q and complement receptor CR3 in preventing synapse loss and improving cognitive function in HD mouse models, and strengthens the rationale for developing complement-related therapeutics, including the anti-C1q antibody ANX005. Future work is needed to determine the extent to which these strategies will be successful in Huntington’s Disease patients.

## Supporting information

Key resources table

## Acknowledgements

We thank Christina Welsh, Emily Lehrman, Lasse Dissing-Olesen, Christina Usher, Matthew Johnson and Anna Kane, for their helpful discussions, reading and editing of this manuscript. We would also like to thank the imaging core at Boston Children’s Hospital and the Harvard Centre for Biological Imaging, including Tony Hill, and Doug Richardson respectively for technical support as well as the IDDRC animal behavioral and physiology core (funded by NIH U54 HD090255), where the visual discrimination and cognitive flexibility tasks were performed, specifically Nick Andrews, Sophie Griswold, Francesco Villa and Nathaniel Hodgson who showed us how to operate these instruments, helped to keep them maintained, provided advice on experimental design and carried out the Optomotor assessments. In addition, we would like to acknowledge Lee Barrett at the BCH Assay Development and Screening Facility and Ivy Pin-Fang at the BCH Human Neuron Core for providing advice and support in helping to automate the ELISA assays. Furthermore, we would like to acknowledge the efforts of Elaine Peskind and Elizabeth Colasurdo at the VA Puget Sound Health Care System, Seattle Division who contributed to the University of Washington CSF collection efforts, Annexon Biosciences (specifically Ted Yednock, Sethu Sankaranaryayanan, Yaisa Andrews-Zwilling and Poojan Suri) for their provision of the C1q function blocking antibody and a version of the C1q ELISA protocol, as well as the measurements of C1q blocking antibody and unbound C1q levels documented in Extended data figure 7c,d and also helpful discussions. In addition, we want to acknowledge Edward Wild at UCL for discussions and suggestions with regards to the CSF and plasma analysis. Finally, we would also like to thank Professor Richard H. Myers at Boston University for helpful discussions and for providing some of the fresh frozen postmortem human tissue. Some of the illustrations in the figures were created with BioRender.com. Work was supported by grants from the NIH (NIH;NINDS GRANT R01NS084298 to B.S. and X.W.Yang; P30-HD-18655 and U54 HD090255, IDDRC Imaging Core) and CHDI (project record A-9181). D.K.W. has been supported by an HD Human Biology fellowship from the Huntington’s Disease Society of America.

## Author Contributions

D.K.W. and B.S. designed experiments and wrote the manuscript; D.K.W., K.M., M.D.H., F.W.G., C R.W., A.F., A.D., X.G., and A.Y.K., performed experiments and analyzed data; K.M., F.W.G., A.F., S.J., T.Y., and X.W.Y provided comments on the manuscript; S.J. carried out clinical assessments of HD patients and collected CSF and serum samples; R.F. provided postmortem human tissue samples from HD patients and control (no clinical diagnosis of a neurodegenerative disease or chronic condition) individuals; T.Y. provided the C1q blocking antibody and participated in helpful discussions; X.W.Y provided expertise and feedback; X.W.Y and B.S. secured funding.

## Data Availability Statement

All of the data supporting the findings of this study can be found within the article, and its extended data and source data files. Extended data figure 1 has associated raw data for the immunoblots that can be located in Source data figures 1 and 2. Some of the data related to biological repository identifiers for the CSF and plasma samples are restricted from distribution as a result of guidelines stipulated in the material transfer agreement that were mandated by the foundation providing this material.

## Code Availability

Details of the custom MATLAB scripts and analysis pipelines used to identify the frequency, amplitude and membrane properties of sEPSCs in the slice electrophysiology experiments are documented in Merel et al., 2016^163^. All of the code related to the two-choice visual discrimination, reversal and progressive ratio tasks will be made freely available upon reasonable request.

## Availability of Biological Materials and Reagents

All unique biological materials with the exception of the C1q function blocking antibody, plasma, CSF and postmortem samples, whose distribution is limited by material transfer agreements, are available from the authors upon reasonable request or from standard commercial sources. Further information about this and requests should be directed to the corresponding author, Beth Stevens.

## Experimental Procedures

### Mice

zQ175 heterozygous mice^71,72^ in which a fragment extending upstream of *htt* exon 1 has been replaced with the human *Htt* sequence containing an expanded number of repeats of the CAG tract, and WT littermates were either bred in-house or obtained from The Jackson Laboratory (JAX) (JAX stock number 027410). BACHD (JAX stock number 008197), BR, BE, BER and WT littermates^19,70^ were obtained from JAX and the laboratory of William Yang at UCLA. C57BL/6J mice were obtained from JAX (JAX stock number 000664) CR3 KO mice (^164^; JAX stock number 003991) were bred in house and crossed with zQ175 mice to generate zQ175 heterozygous CR3 KO mice. Homer GFP mice^102^ were obtained from the laboratory of Shigeo Okabe at Tokyo medical and dental university, bred in house and crossed with zQ175 mice to generate zQ175 heterozygous Homer GFP mice. We used P1 or P2 and P300 mice for motor cortex injections and P120 mice for intraperitoneal (IP) injections. For all other experiments mice were used at the ages specified in the procedures listed below and displayed in relevant figure panels and legends. Unless otherwise stated, a mixture of male and female mice were employed in all experiments. Animals were group housed in Optimice cages and maintained in the temperature range and environmental conditions recommended by AAALAC. All experiments were approved by the institutional care and use committee of Boston Children’s Hospital and UCLA in accordance with National Institutes of Health (NIH) guidelines for the humane treatment of animals.

### Mouse Genotyping

DNA was extracted from ear clips using the Hot Shot method described under the Quick DNA purification section of the Jackson Laboratory website (https://www.jax.org/jax-mice-and-services/customer-support/technical-support/genotyping-resources/dna-isolation-protocols;^165^) or from tail clips using the Phenol/ chloroform method described on The Jackson Laboratory website (https://www.jax.org/jax-mice-and-services/customer-support/technical-support/genotyping-resources/dna-isolation-protocolsz). zQ175 heterozygous mice were genotyped for the CAG expansion using the protocol provided by The Jackson Laboratory and developed by Laragen, Inc and for the Neo cassette using the protocol provided by the Jackson laboratory. CR3KO and Homer GFP mice were genotyped using the below protocols for reagent setup and thermocycling. Primer details can also be found in the key resources table. In all cases the PCR product was visualized on 2% agarose gels with ethidium bromide.

### CR3 genotyping

#### Reagent setup

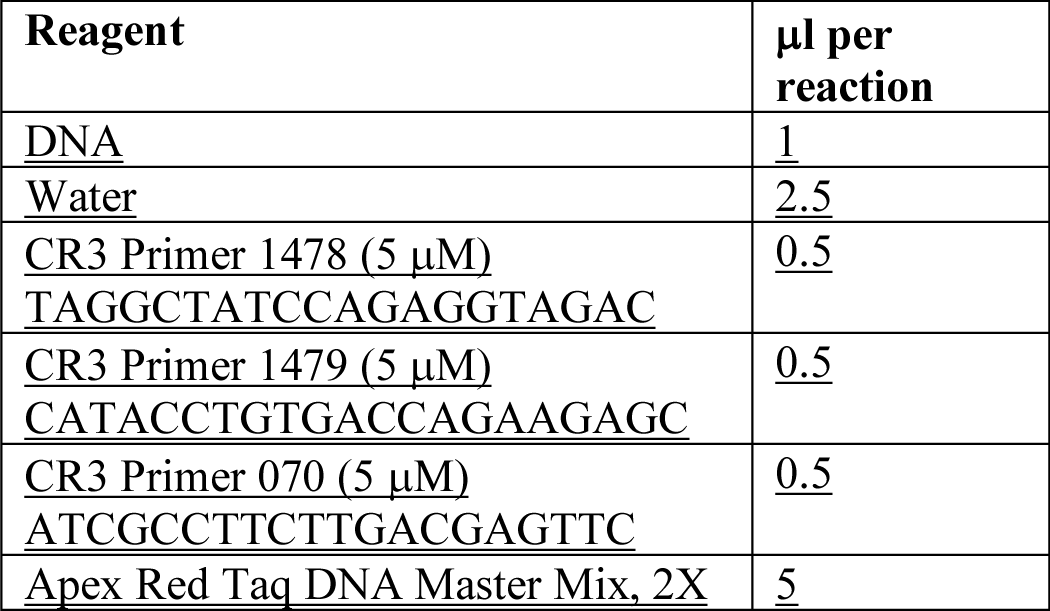

#### Thermocycling

| Step | Temperature | Time/cycles |
| --- | --- | --- |
| <u>1</u> | <u>94°C</u> | <u>3 min</u> |
| <u>2</u> | <u>94°C</u> | <u>30 sec</u> |
| <u>3</u> | <u>58°C</u> | <u>1 min</u> |
| <u>4</u> | <u>72°C</u> | <u>1 min</u> |
| <u>5</u> | <u>Go to Step 2</u> | <u>35 cycles</u> |
| <u>6</u> | <u>72°C</u> | <u>2 min</u> |
| <u>7</u> | <u>10°C</u> | <u>Forever</u> |

#### Expected band sizes

<u>CR3 KO 700bp</u>

<u>CR3 WT 350bp</u>

### Homer-GFP 370 bp

#### Reagent setup

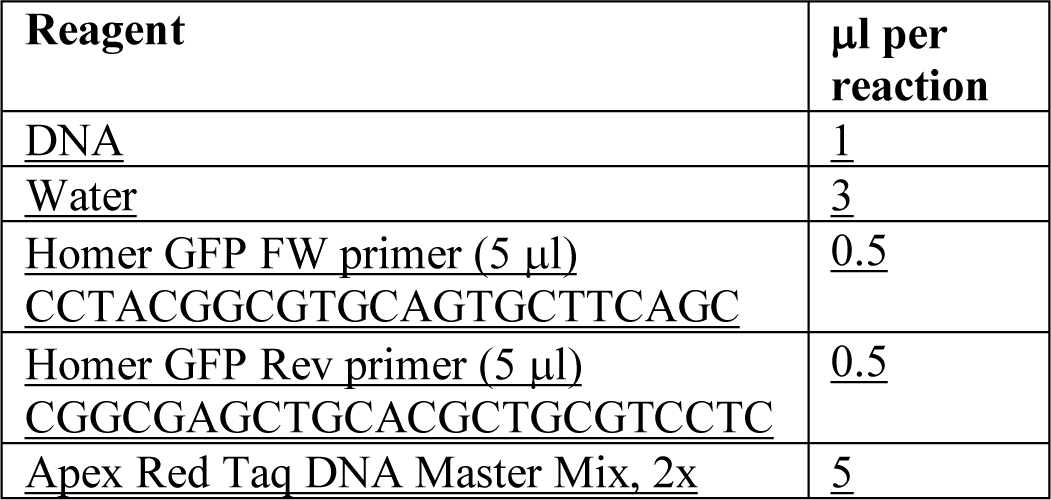

#### Thermocycling

| **Step** | **Temperature** | **Time/cycles** |
| --- | --- | --- |
| <u>1</u> | <u>95°C</u> | <u>5 min</u> |
| <u>2</u> | <u>95°C</u> | <u>30 sec</u> |
| <u>3</u> | <u>50°C</u> | <u>30 sec</u> |
| <u>4</u> | <u>72°C</u> | <u>1 min</u> |
| <u>5</u> | <u>Go to Step 2</u> | <u>30 cycles</u> |
| <u>6</u> | <u>72°C</u> | <u>5 min</u> |
| <u>7</u> | <u>10°C</u> | <u>Forever</u> |

#### Expected band sizes

Homer-GFP 370 bp

### CSF and Plasma samples (clinical details for all samples can be found in Supplemental table 1)

(i) HDClarity cohort: HDClarity is a multi-site cerebrospinal fluid collection initiative designed to facilitate therapeutic development for Huntington’s disease. It incorporates both longitudinal and cross sectional sampling (HDClarity clinical protocol at hdclarity.net/). Prior to sample collection appropriate ethical approval including ERB and IRB consent was obtained from all clinical sites in accordance with the rules and regulations in those countries. All participants were required to provide informed consent prior to undertaking study procedures and these informed consents were obtained by clinical site staff using approved processes.

#### Criteria for subject group placement

##### Healthy controls

Individuals who have no known family history of HD or have been tested for the huntingtin gene glutamine codon expansion and are not at risk.

##### Early premanifest HD

Individuals who do not have any clinical diagnostic motor features of HD, defined as Unified Huntington’s Disease Rating Scale (UHDRS) Diagnostic score <4 and who have a GAG expansion of ≥ 40 as well as a burden of pathology score (computed as (CAG-35.5) x age) of <250 *.

##### Late premanifest HD

Individuals who do not have any clinical diagnostic motor features of HD, defined as UHDRS score <4 and who have a CAG expansion of ≥ 40 as well as a burden of pathology score of >250 *.

##### Early manifest HD

Individuals who have a UHDRS score = 4 and a CAG expansion of ≥36 and have a UHDRS total functional capacity (TFC) score between 7 and 13 inclusive.

*Stratifying premanifest HD patients into two groups on the basis of their age and number of CAG repeats has been shown to reveal differences in total brain volume, subcortical grey matter volume, cortical thickness, white matter microstructure, resting-state cerebral blood flow, functional activity of the basal ganglia while performing a time discrimination task, multiple measures of motor and cognitive performance and differences in neuropsychiatric assessments^1,26,89,132,134,166^. Also, particularly pertinent to this study, this stratification has shown differences in the levels of a putative fluid biomarker of disease progression^133^.

The burden of pathology score used to separate early and late premanifest HD patients in this study yielded a comparable division to the that obtained by Tabrizi et al and Mason et al who divided their premanifest patients into 2 groups (termed PreHD-A and PreHD-B) using the group median for predicted years to onset as determined by the survival analysis formula described by Langbehn and co-workers^132,166,167^. In fact only 3 samples were differently allocated between the early and late premanifest groups when comparing these two stratification approaches and this did not affect the disease stage dependent differences in CSF levels of C3 and iC3b (data not shown). Stratifying early and late premanifest HD patients on the basis of a threshold number of years to onset of 10.8 (determined using the Langbehn formula) as employed by Byrne et al., (on the basis of the group median predicted years to onset for the sample cohort in their study) also did not affect the observed disease stage dependent differences in CSF levels of C3 and iC3b (data not shown)^133,162^. For our sample cohort the median estimated years to diagnosis for early premanifest HD patients when employing the Langbehn formula is 26, which contrasts to a value of 9.84 for the late premanifest HD patients. This is very similar to that reported for the Cambridge cohort in Mason et al., (22.1 and 10.8 respectively), in which differences in subcortical volumes were observed^166^.

#### Sample collection

For CSF samples, lumbar punctures were carried out according to the guidelines stipulated in the HDClarity clinical protocol and with the approval of the relevant institutional review boards. Briefly, patients were placed into the lateral decubitus position and CSF extracted from the subarachnoid space at the L3-4 or L4-5 interspace using a spinal needle. All samples were collected in a highly standardized manner with the collection time, method of extraction and storage all controlled (for further details please see the HDClarity clinical protocol at hdclarity.net/). In addition, patients underwent blood tests and clinical screening to account for any confounding variables such as underlying infections. CSF samples were also screened for blood cell contamination of >1000 erythrocytes per μl and were then aliquoted and maintained at -80°C until use (for further details please see the HDClarity clinical protocol at hdclarity.net/). In order to further examine the degree of blood contamination or ascertain whether there is any evidence of blood brain barrier breakdown in our samples the CSF concentration of albumin, a molecule which constitutes around 50% of the total protein in the plasma, and which has also been shown in many studies to be present at significantly lower levels in the CSF, was quantified (Extended data Fig. 10j,k) ^168-171^. Previous studies using albumin labelled with radioactive iodine have shown that the blood brain barrier has a relatively low permeability for this species relative to other plasma proteins that undergo receptor mediated transport and that it can reflect blood brain barrier disruption under certain experimentally defined paradigms^118,172^. We found that for all of the HDClarity samples CSF albumin concentrations were below the average level found in a previous study of a large patient cohort (over 1000 individuals whose serum:CSF albumin ratio was considered normal)^173^. Importantly there was also no disease stage specific changes in CSF albumin levels and no correlation of albumin levels with patient CAP score either when all HDGEC’s were considered or just when samples from premanifest patients were interrogated (Extended data Fig. 10j,k). These findings are consistent with a previous study which found no difference in the serum:CSF albumin ratio when comparing CSF samples from HD patients and control (clinically normal) individuals^174^ and suggest that any disease stage associated changes in the analytes we are assessing are unlikely to be driven by differences in the levels of blood contamination observed in these patient subject groups.

For plasma samples venous blood collection was carried out according to the guidelines stipulated in the HDClarity clinical protocol and with the approval of the relevant institutional review boards. Briefly, blood was drawn immediately after CSF sample collection was complete and collected in lithium heparin tubes. Tubes were then spun at 1300g for 10 min at 4°C and the plasma supernatant aliquoted and stored at -80°C. If the supernatant appeared pink after spinning implying that hemolysis had taken place the sample was discarded.

(ii) University of Washington cohort: This was a prospective single-site study with standardized longitudinal collection of CSF, blood and phenotypic data. The study was performed in compliance with the institutional review board protocol at the University of Washington. All participants were required to provide informed consent prior to undertaking study procedures and these informed consents were obtained by clinical site staff using approved processes.

#### Criteria for subject group placement

##### Healthy controls

Individuals who have no known family history of HD or have been tested for the *Huntingtin* gene glutamine codon expansion and are not at risk.

##### Premanifest HD

Individuals who do not have any clinical diagnostic motor features of HD, defined as Unified Huntington’s Disease Rating Scale (UHDRS) Diagnostic score <4 and who have a GAG expansion of ≥ 40 as well as a burden of pathology score (computed as (CAG-35.5) x age) of between 130 and 560.

#### Sample collection

For CSF samples, lumbar punctures were carried out according to the guidelines stipulated in the institutional review board protocol at the University of Washington. Briefly, patients were placed into the lateral decubitus position and CSF extracted from the subarachnoid space at the L3-4 or L4-5 interspace using a spinal needle. All samples were collected in a standardized manner with the method of extraction and storage controlled. CSF samples were screened for blood cell contamination of ≤ 10 erythrocytes and maintained at -80°C. They were thawed on ice immediately prior to use. To further examine the degree of blood contamination or ascertain whether there is any evidence of blood brain barrier breakdown in our samples the CSF concentration of albumin, a molecule which constitutes around 50% of the total protein in the plasma, and which has also been shown in many studies to be present at relatively lower levels in the CSF, was quantified^168-171^. Previous studies using albumin labelled with radioactive iodine have shown that the blood brain barrier has a relatively low permeability for this species relative to other plasma proteins that undergo receptor mediated transport and that it can reflect blood brain barrier disruption under certain experimentally defined paradigms^118,172^. We found that for all of the University of Washington samples CSF albumin concentrations were below the average level found in a previous study of a large patient cohort (over 1000 individuals whose serum:CSF albumin ratio was considered normal)^173^. Importantly there was also no correlation of albumin levels with patient CAP score (Extended data Fig. 11j). These findings are consistent with a previous study which found no difference in the serum:CSF albumin ratio when comparing CSF samples from HD patients and control (clinically normal) individuals^174^. and suggest that any disease stage associated changes in the analytes we are assessing are unlikely to be driven by differences in the levels of blood contamination observed in these patient subject groups.

For serum samples venous blood collection was carried out according to the guidelines stipulated in the institutional review board protocol at the University of Washington. Briefly, blood was drawn and allowed to clot before centrifuging at 1000-2000g for 10 min at 4°C. Serum supernatant was then aliquoted and stored at -80°C.

### Postmortem human tissue (clinical details for all samples can be found in Supplemental table 2)

Fresh frozen tissue from the globus pallidus and caudate putamen of Huntington’s disease patients and control individuals was a kind gift from Professor Richard H. Myers at Boston University and the Los Angeles VA Human brain and spinal fluid resource center. Fixed tissue sections were obtained through a collaboration with Dr Richard Faull and Marika Esez at the New Zealand Human Brain Bank (the University of Auckland). For all fixed samples brain tissue was perfused with PBS containing 1% sodium nitrite followed by a 15% formalin solution and post fixation for 12-24hr. Samples were subsequently cryopreserved in 30% sucrose and snap frozen before being sectioned. The Huntington’s tissue assessed in this manuscript was classified as having a neuropathological score of either Vonsattel grade 2 or grade 4. Grade 2 is classified as displaying mild to moderate striatal atrophy with the medial outline of the HCN being only slightly convex but still protruding into the lateral ventricle. Significant loss of neurons and increases in astrocyte number are evident in the dorsal half of the caudate nucleus and adjacent putamen. Astrocytes also show altered size and increased GFAP immunoreactivity. Initial signs of neurodegeneration are evident in the pallidum with the external segment displaying greater atrophy than the internal segment. Grade 4 is classified as displaying severe striatal atrophy with the medial contours of the internal capsule and head of the caudate nucleus now concave. There is also a 95% loss of striatal neurons^59,175^.

### C1q function blocking Antibody

The C1q function blocking antibody, termed ANX-M1 in some publications^30,34,116,176-179^, was generated by Annexon Biosciences (Annexon ATCC accession number PTA-120399) according to the protocol outlined in Hong et al., 2016^30^ and batch tested in rodent hemolytic assays (for details see Hong et al., 2016^30^). ANX-M1 is the murine parent antibody of the humanized anti-C1q antibody ANX005, currently being employed in a phase 2 clinical study in HD patients (clinical trials.gov NCT04514367). Conjugation to FITC or ammonium peroxidase was carried out using Abcam’s alkaline phosphatase and FITC conjugation kits (Abcam ab102850 and ab102884 respectively) according to the protocols outlined in the manufacturer’s instruction manuals. Measurements of serum levels of ANX-M1 and unbound C1q in mice treated with the C1q blocking antibody or control IgG were carried out according to the protocols outlined in Lansita et a., 2017 ^117^

### Motor Cortex Injections

#### P1/P2 mice

Mice were anesthetized on ice and a glass capillary attached to a stereotax used to deliver 0.4μl of pAAV2-hsyn-EGFP <u>(Addgene viral prep # 50465-AAV2;</u>http://n2t.netaddgene:50465;RRID:Addgene_50465) into the motor cortex using a Nanoject III programmable nanoliter injector (VWR 490019-810) and a micromanipulator MM33 (Pipette.com 3-000-024-R). Mice were subsequently allowed to recover on a warm plate before being transferred back to their home cage. At P120 mice were sacrificed and microglial engulfment assayed as described below. Using this AAV serotype and promoter no microglia could be seen expressing the construct in the cortex but both types of corticostriatal projection neurons, the pyramidal tract (PT)-type neurons, which are mainly found in layer V, and the intratelencephalically (IT)-type neurons which are mainly found in layer III and upper half of layer V were targeted (Extended data Fig. 6e,f). As expected VGLUT1 immunolabelled synaptic terminals, denoting corticostriatal synapses, were found to colocalize with GFP signal in the dorsal striatum of P120 injected mice but no overlap of GFP signal was observed with markers of the thalamostriatal synapse (Fig. 4a). This finding is comparable to what was observed with the use of anterograde tracers in Lei et al., 2013^67^.

#### P300 mice

Mice were anesthetized with isoflurane and a drill used to open a small hole in the skull. A glass pipette attached to a stereotax was subsequently used to deliver 0.4μl of pAAV2-hsyn-EGFP (Addgene viral prep # 50465-AAV2;http://n2t.netaddgene:50465;RRID:Addgene_50465). The skin was then closed with sutures and the mice allowed to recover in a warm chamber before being transferred back to the home cage. 14 days post injection mice were sacrificed and microglial engulfment assayed as described below.

### Intraperitoneal Injections

P120 mice were injected IP with either 40mg/kg of the C1q function blocking antibody or a control IgG (BioXCell BE0083) every 48hr for 4 weeks. A comparable injection paradigm has previously been shown to enable uptake of Iodine-125 labelled IgG antibodies into the brain^119^ and led to the C1q function blocking antibody becoming detectable in the CSF^117^. The weight of the mice was monitored throughout the injection period and did not show any significant changes with either treatment (Extended data Fig. 7e,f).

### Ex vivo slice electrophysiology

#### Acute brain slice preparation

Following 1 mo of treatment with either the C1q function blocking antibody or the control IgG, coronal sections (300 μm thickness) containing the striatum were prepared from the brains of 5 mo zQ175 and WT littermates (as well as those of untreated mice). As described previously^180^, mice were anesthetized with isoflurane and transcardially perfused with ice-cold artificial cerebrospinal fluid (ACSF) solution (in mM): 125 NaCl, 2.5 KCl, 25 NaHCO_3_, 2 CaCl_2_, 1 MgCl_2_, 1.25 NaH_2_PO_4_ and 11 glucose (300-305 mOsm/kg). After slicing the tissue in ice cold ACSF on a Leica VT1200, slices were transferred for 10 min at 33°C into a chamber filled with a choline-based recovery solution (in mM): 110 choline chloride, 25 NaHCO_3_, 2.5 KCl, 7 MgCl_2_, 0.5 CaCl_2_, 1.25 NaH_2_PO_4_, 25 glucose, 11.6 ascorbic acid, and 3.1 pyruvic acid. Slices were then transferred to a second 33°C chamber containing ACSF for 30 mins. Following this recovery period, the chamber was moved to room temperature for the duration of the experiment.

To isolate spontaneous excitatory postsynaptic currents (sEPSCs), recordings were performed at 28°C in ACSF containing gabazine to block inhibitory activity. Electrodes with a resistance of 2.5-4.5 MΩ were pulled from borosilicate glass (Sutter Instruments) and filled with a Cesium-based internal solution (in mM): 135 CsMeSO_3_, 10 HEPES, 1 EGTA, 3.3 QX-314 (Cl^−^ salt), 4 Mg-ATP, 0.3 Na-GTP, 8 Na_2_-Phosphocreatine (pH 7.3 adjusted with CsOH; 295 mOsm·kg^−1^). At a holding potential of -80 mV, spontaneous activity was recorded for 5 to 10 min per cell. All recordings were carried out blind to genotype and treatment.

#### Spontaneous excitatory postsynaptic current analysis

Data was collected with a Multiclamp 700B amplifier (Molecular Devices) and National Instruments acquisition board using custom ScanImage MATLAB software (Mathworks). Voltage clamp recordings were filtered at 2 kHz and digitized at 10 kHz. Only recording epochs with stable baseline potentials (<10% change) and access resistance (+/- 10% baseline) were utilized for analysis. To identify the frequency and amplitude of sEPSCs, custom MATLAB scripts and analysis pipeline were adapted from Merel et al., 2016^163^. This process was carried out blind to genotype and treatment. Statistical analyses were subsequently performed using GraphPad Prism software (GraphPad) and MATLAB.

### Immunohistochemistry

Immunohistochemistry was performed as described in Schafer et al., 2012^96^ and Lehrman et al., 2018^156^. Briefly, brains were harvested from mice following transcardial perfusion with 15ml PBS and 15ml 4% paraformaldehyde (PFA). Tissue was then postfixed in 4% PFA for 2 h, before being washed in PBS and transferred to a 30% sucrose solution for approximately 24-48hr. Subsequently tissue samples were embedded in a 2:1 mixture of 30% sucrose-PBS: Tissue-Tek O.C.T. Compound (Electron Microscope Sciences) and stored at -80°C. 14 μm or 25 μm cryosections were then cut using a Leica cryostat and affixed to Leica Surgipath X-tra slides before being processed for immunohistochemistry as follows. Slides were heated for 10 min at 35°C followed by 3 rinses in PBS. They were then blocked with either blocking solution (150 mM NaCl, 50mM Tris Base pH7.4, 5% BSA, 100mM L-Lysine, 0.04% sodium azide) and 0.3% Triton-X100 for staining with complement antibodies or 10% normal goat serum and 0.3% Triton-X 100 solution (Sigma) for 1 hr, before being incubated with primary antibodies overnight (O/N) at 4°C in 4:1 antibody buffer (150mM NaCl, 50mM Tris Base pH7.4, 1% BSA, 100mM L-lysine, 0.04% azide) : blocking solution or 5% normal goat serum (Sigma) and 0.3% Triton X-100 solution (Sigma). Slides were then washed for 3x 15 min in PBS followed by incubation with appropriate secondary antibodies in 4:1 antibody buffer : blocking solution or 5% normal goat serum for 2 hr at room temperature. Slides were then washed for 3x 15 min in PBS and mounted with Vectashield with DAPI (Vector Laboratories H-1000). For engulfment analysis, microglial density quantifications and Iba-1:P2RY12 or Iba-1:TMEM119 colocalization analysis 40 µm free-floating sections were cut with a sliding microtome (Leica) and stained with relevant antibodies in 10% normal goat serum (NGS) and 0.3% Triton-X 100 O/N at room temperature. All other steps are identical to those described above. After the final wash the tissue was spread on the slide and a coverslip mounted on top using Vectashield with DAPI (Vector Laboratories H-1000).

For the engulfment analysis images were acquired using an UltraVIEW VoX spinning disc confocal microscope and a 60x plan apochromat oil objective (1.4NA). Volocity image acquisition software was employed (Perkin Elmer). For other confocal images either an LSM 700 confocal microscope with 25x or 63x plan apochromat oil objectives (0.8NA and 1.4NA respectively) and Zen 2009 image acquisition software (Carl Zeiss), or an LSM 880 confocal microscope with 10x dry or 63x oil plan apochromat objectives (0.45NA and 1.4NA respectively) and Zen Black 2.3 image acquisition software (Carl Zeiss) or a LEICA SP8 white light confocal with a 100x plan apochromat oil objective (1.4NA) was used (N.B. for each different types of analysis performed the same microscope and imaging setup was used to collect images from every sample). For structured illumination microscopy a Zeiss ELYRA PS1 microscope with a 100x plan apochromat objective (1.57NA) and Zen Black 2012 image acquisition software was used. Finally for array tomography analysis an AxioImager Z1 microscope with a 60x plan-neofluar objective (1.4NA) and Axiovision software (Carl Zeiss) was used.

The antibody dilutions used were as follows: Homer1 (Synaptic systems (160003)) 1:200, VGLUT2 (Millipore Sigma (AB2251)), 1:1000 IHC, 1:10000 WB, VGLUT1 (Millipore Sigma (AB5905)) 1:1000 IHC, 1:10000 WB, Iba1 (Wako (019-19741)), 1:400, Iba1 (Wako (ncnp24)) 1:200, CD68 (Serotec clone FA-11 (MCA1957)), 1:200, CD68 (Dako (M087629-2)) 1/200, CD11b (Serotec clone 5C6 (MCA711G)), 1:200, clone 5C6, β -actin (Sigma (A2228)), 1:1000 WB, C1q (Abcam (ab182451)) (validated in C1qA KO mice https://www.abcam.com/c1q-antibody-48-ab182451.html#description_images_1), 1:500, C1q (Dako (A0136)) 1:500, C1q [JL-1] (Abcam (ab71940)) 1:500, C3d (Dako (A0063)), 1:500, C3c (Dako (F0201)) 1:1000, iC3b (Quidel (A209)) 1:500, PSD-95 (Millipore (MAB1596)) 1:500; S100□ (Dako (Z0311)) 1:500; TMEM119 (Abcam (ab209064)) 1:200, P2RY12 (Anaspec (AS-55043A)) 1:500 anti-fluorescein-POD (Roche (11 426 346 910)) 1/2000, anti digioxigenin (Roche (11207733910)) 1:2000, alexa-fluor conjugated secondary antibodies (Life Technologies) 1:250, Goat anti-Rabbit IgG H&L alkaline phosphatase (Abcam (ab97048)) 1/5000, Goat anti rabbit HRP (Promega (W4011)) 1/5000, Peroxidase-AffiniPure Donkey anti-guinea pig IgG (H+L) (Jackson ImmunoResearch 706-035-148). Further details can be found in the key resources table.

### Microglia density quantification

To quantify microglial cell density 40 μm sections from zQ175 heterozygote mice and WT littermates, collected at the specified ages, were stained with Iba1 and 3 fields of view in the dorsal striatum were imaged per animal (n = 4-5 per genotype). Microglia were then manually counted in each 25x magnification field (65571 μm^2^) using ImageJ software (NIH) and these values were averaged to generate a per field number for each animal. All analysis was performed blind to genotype.

### Assessing changes in microglia morphology and lysosomal protein levels

25 μm sections from zQ175 heterozygote mice and WT littermates, collected at the specified ages, were immunolabeled with CD68 and Iba1 and imaged with a 25x pan apochromat oil objective (0.8NA) using a Zeiss LSM700 confocal microscope with 0.4 μm z steps. 3 fields of view were captured in relevant brain regions and maximum intensity projections (MIPs) generated with Zeiss software. The phagocytic state of each microglia was categorized based on morphology and CD68 levels using an adapted version of the protocol in Schafer et al., 2012^96^ with 0 being the least phagocytic and 5 the most. Briefly process morphology was scored as 0 (thin long processes with multiple branches), 1 (thicker processes but with comparable branching), 2 (thick retracted processes with few branches), 3 (no clear processes). CD68 was analyzed and scored as 0 (no/scarce expression), 1 (punctate expression), 2 (aggregated expression or punctate expression all over the cell). For each cell analyzed, morphology and CD68 scores were summed and a final score for microglia phagocytic state (0-5) was assigned. Between 30 and 50 cells were assessed per animal and the number of microglia with a given score was represented as a percentage of the total population. All analysis was performed blind.

Sholl analysis was performed by surface rendering Iba1 staining using Imaris 9.3 software with the use of splitting algorithms to identify individual cells. The Filaments tool in Imaris was then used to trace individual cells, and subsequently identify and quantify both branch points and terminal projections at different distances from the soma. Over 100 cells were analyzed for each genotype and all analysis was performed blind.

### Assessing changes in CD11b levels

Sections from 3 mo zQ175 heterozygote mice and WT littermates were immunolabelled with CD11b and Iba1 antibodies and imaged with a 25x pan apochromat oil objective (0.8NA) with 0.4 μm z steps. 3 Fields of view were captured in relevant brain regions and maximum intensity projections (MIPs) generated with Zeiss software. Image J software was then used to assess the mean grey value of CD11b staining within individual fields of view which were subsequently averaged to generate a value for each animal. At least 30-50 cells were analyzed from each animal and all analysis was performed blind to genotype.

### Engulfment analysis

zQ175 heterozygous homer GFP mice and their WT homer GFP littermates or zQ175 heterozygous mice and WT littermates which had received an injection of pAAV2-hsyn-EGFP were sacrificed at P120 and P314 and transcardially perfused with 15 ml ice cold PBS followed by 15 ml of 4% PFA. The brains of these mice were then dissected out and post fixed for 2 h in 4% PFA before being washed and placed in a 30% sucrose solution in PBS for storage at 4°C for 24-48 h (as described above). Brains were subsequently sectioned on a sliding microtome (Leica) and 40 µm sections were stained with Iba1 or S100β (as described above). S100β is thought to be a marker of all known astrocyte populations in the striatum, and has identified comparable numbers of cells to those seen in ALDH1eGFP mice^104^. Images from the dorsal striatum ipsilateral to the injection site were then acquired on an UltraVIEW VoX spinning disc confocal microscope using a 0.2 μm z step. For each animal 10-15 cells were imaged. Images were subsequently processed and analyzed as described previously using Image J (NIH) and Imaris software (Bitplane)^96,101,156^. The data generated was used to calculate percent engulfment (volume engulfed GFP/volume of the cell) and input density (total volume GFP inputs/volume of field of view). All experiments were performed blind to genotype.

One potential caveat of this method of engulfment analysis is that it cannot exclude the contribution of autofluorescent granules, which have been reported to be present in the lysosomes of microglia, to the signal being analyzed^181-183^. However, when we quantified autofluorescence (the signal generated by exciting at a wavelength that will not activate the GFP fluorophore being expressed or any of the other fluorophores used for staining) we observed that less than 10% of the CD68 stained lysosome area was actually autofluorescent and more importantly that there was no difference in the autofluorescence area per field or the area of autofluorescence colocalized with CD68 when comparing WT and zQ175 heterozygote mice (Extended data Fig. 6c,d).

### Synapse quantification and complement co-localization with synaptic markers

#### For confocal images

A modified version of the protocol outlined in Hong et al., 2016^30^ was employed. Briefly,14 µm sections were collected from mice with different genotypes at the ages specified in the text and figure legends and from mice following treatment with different agents. They were stained with appropriate antibodies to synaptic or complement proteins and imaged with either a Zeiss LSM 700 or Leica SP8 white light confocal microscope. Three fields of view were captured at 63X magnification in relevant brain regions (101.6 μm^2^), and for each field a 3 µm z-stack comprised of 1µm z-steps was imaged. Z-planes were captured at a depth at which complement and synaptic staining was uniform across the field of view. ImageJ software was subsequently used to quantify the number of colocalized pre and postsynaptic puncta or the number of colocalized complement and presynaptic puncta in each z-plane (9 images total for each animal) using a method that incorporates thresholding the images into a binary state and identifying overlapping puncta which fulfil specific dimension criteria. These were then averaged to generate a per field number for each animal. All analysis was performed blind to genotype and treatment.

#### For structured illumination images

A modified version of the protocol outlined in Hong and Wilton et al., 2017^184^ was employed. Briefly, 14 □m sections from zQ175 heterozygote mice, zQ175 CR3KO mice and WT littermates were stained with appropriate synaptic markers or antibodies to complement proteins and imaged with a Zeiss Elyra PS1 microscope. Three fields of view were captured at 100x magnification in relevant brain regions, and for each field a 3 μm z stack comprised of 0.01 μm z steps was imaged. Zeiss software and proprietary algorithms were subsequently used to generate SIM-processed image files (images can appear slightly saturated because the demodulation algorithm used to reconstruct the image lacks a means of converting the raw detected signal into photon counts). To quantify different types of synapses or the number of synapses colocalized with different complement proteins, image files were opened in Imaris and spot channels generated for each set of immunoreactive puncta using dimensions determined empirically from averaged measurements. A MATLAB plugin (MathWorks) was then used to display and quantify only those spots within a threshold distance from each other (measured from the center of each spot). Depending on the analysis the number of colocalized spots was normalized to the total number of one of the spot populations. For the complement colocalization experiments an additional control was performed by rotating the channel containing the complement staining 90 degrees relative to the VGLUT1 images to enable assessment of the amount of colocalization that would be expected to occur via chance. Statistical tests were then used to compare the fold change relative to WT in the % of synaptic puncta colocalizing with complement proteins in the rotated vs the non-rotated images. All analysis was performed blind to genotype.

#### For array tomography

Array tomography was performed as previously described with minor modifications^1,92,96,185-188^. Briefly, 300 μm vibratome sections of dorsal striatum from 7 mo WT and zQ175 heterozygote mice were fixed in 4% paraformaldehyde for 1.5 h at room temperature and embedded in LR White resin. This tissue was then further sectioned to generate ribbons of between 20-30 serial 70nm thick sections, which were mounted on subbed glass coverslips and immunostained with VGLUT1 and Homer1 antibodies. The same field of view was subsequently imaged on serial sections using a Zeiss AxioImager Z1 microscope with a 63x objective. A 3D projection of this field was then obtained by aligning these images with ImageJ (NIH) and AutoAligner software (Bitplane AG). Finally, projections were analyzed using Imaris (Oxford instruments), to quantify colocalized puncta using the spots function, as outlined above for analysis of structured illumination images. Analysis was performed blind to genotype.

### Detecting C1q function blocking antibody in the neuropil

P120 zQ175 heterozygous mice were injected IP with 40mg/kg of FITC conjugated or unconjugated M1. 24 hr post injection mice were sacrificed and their brains harvested following transcardial perfusion with PBS and 4% paraformaldehyde (PFA). Tissue was then postfixed in 4% PFA for 2 h, washed and transferred to a 30% sucrose solution. 14 μm cryosections were subsequently prepared from tissue embedded in a 2:1 mixture of 20% sucrose: OCT, and blocked with a 5% bovine serum albumin (BSA) and 0.2% Triton-X 100 solution for 1 h, before being incubated with anti-fluorescein-POD (Roche (11 426 346 910 1/2000) O/N. Signal was detected and amplified using the TSA staining Kit (Perkin Elmer (NEL0701001KT)) according to the manufacturer’s instructions

### Immunohistochemistry using postmortem human tissue sections

Postmortem human tissue was fixed according to the procedures outlined in Waldvogel et al., 2008^189^. Briefly, the brain was extracted and the basal and internal carotid arteries identified. The tissue was subsequently perfused by attaching winged infusion needles to these arteries and flowing through a solution of PBS containing 1% sodium nitrite followed by 3 liters of a fixative containing 15% formalin. After perfusion, the brain was post fixed for 12-24hr in the same fixative before being dissected into blocks for sectioning. Tissue was stained by washing 25 μm free floating postmortem human tissue sections for 3x 5 min in PBS and then permeabilizing them in a 0.2% triton X-100 PBS solution for 1 h at room temperature. Sections were subsequently blocked in a 10% BSA, 0.2% triton X-100 PBS solution for 1 h at room temperature before applying appropriate primary antibodies in a 5% BSA, 0.2 % triton X-100 PBS solution O/N at 4°C. Sections were then washed for 3x 5 min in PBS before adding appropriate secondary antibody in 5% BSA PBS for 1 h at room temperature. After washing for a further 3x 5 min in PBS sections were incubated in a 0.5% Sudan Black solution dissolved in 70% ethanol to reduce autofluorescence from lipofuscin vesicles. Sections were then washed a further 7x in PBS to remove excess Sudan Black before being spread onto slides and left to dry. Coverslips were mounted in 90% glycerol PBS containing Hoechst (diluted 1/1000).

### Construction of a C3 expression construct

A mammalian expression plasmid containing the cDNA for mouse C3 was constructed according to the protocol outlined below. C3 cDNA obtained from OpenBiosystem/Horizon (Cat# MMM1013-202768722; clone Id# 5134713) was amplified using a two-step strategy. In the 1^st^ step the restriction site Sma I and a Kozac sequence were added to the 5’ end before a sequence encompassing everything up to the Xba I restriction site (2287bp) was amplified. In the 2^nd^ step a restriction site BamH I and a stop codon were added at the end of the gene (2705bp). Both sequences were subsequently inserted into the pUltra EGFP plasmid (Addgene 24129) by subcloning with Xba I and BamH I. The primer sequences employed were as follows: C3 beg Smal 1F (cccggggccaccatgggaccagcttcagggtcccagc); C3 beg Xbal 2R (tctagagataatatcttcttctgg); C3 end Xbal 1F (tctagaagccacttcccacagagc); C3 end BamH I 2R (ggatcctcagttgggacaaccataaacc).

### Testing the specificity of the anti-C3d antibody by carrying out immunocytochemistry (ICC) on HEK cells expressing a mouse C3 construct

HEK 293 cells (ATCC ref CRL-1573) grown on coverslips were transfected using Lipofectamine 2000 (Invitrogen 11668-019) and 1-2 μg of either pULTRA EGFP or pULTRA EGFP T2A C3 Ms (see above) (a culture in which no DNA was employed in the transfection was included as a control). 24 h following transfection cells were fixed using 4% paraformaldehyde for 10 min at room temperature. After washing for 3x 5 min with PBS cells were subsequently blocked for 30 min in a 10% goat serum PBS solution. Cells were then incubated O/N at 4°C with anti C3d diluted 1/500 in a 10% goat serum 0.2% triton PBS solution. After washing for 3x 5 min with PBS goat anti rabbit Alexa 594-conjugated secondary antibody diluted 1/1000 in PBS was added to the cells for 30 min at room temperature. Cells were then washed again and the coverslips mounted using Vectashield with DAPI (Vector Laboratories H-1000). Staining was subsequently visualized on a Zeiss axio imager Z.1 (Extended data fig. 1j). Note that while the complement C3d antibody employed here was raised against purified C3d from human plasma, it is a polyclonal entity and as such contains species that bind to multiple different epitopes within the structure of the protein. Given that the mouse and human protein sequences for complement component C3 share a 77% identity (NCBI BLAST), the cross reactivity of this antibody, which we demonstrate in Extended data figure 1j, should not be unexpected.

### Fluorescence in situ hybridization

*C1qa/C1QA*, and *C3* antisense RNA probes were made and labeled with Digoxigenin as described in Liddelow et al., 2017^190^, while the NSE antisense RNA probe was labeled with Fluorescein. Briefly plasmids for human *C3* (Open Biosystems reference MMM1013-202769931), human *C1QA* (Open Biosystems reference MMM1013-202702004) and mouse *C1qa* (Open Biosystems reference MMM1013-202704027) were digested with Sal I before RNA was transcribed from the T7 promoter. Alkaline hydrolysis was then performed at 60°C for 30 min to fragment the target RNA before labeling it with Digoxigenin and Fluorescein according to kit instructions (Roche).

Labelled RNA probes were subsequently applied to 14 μm cryosections of mouse and human tissue according to the procedure outlined below. Briefly, sections were initially prepared as described above for IHC analysis, before being dried at 65 °C for 30 min. Sections were then incubated in 100% methanol at -20 °C for 20 min before being washed for 3x 5 min in PBS. Sections were subsequently treated with a 1 μg/ml proteinase K solution (1 μg/ml in 50 mM Tris pH7.5 and 5 mM EDTA) for 10 min before being washed again for 3x 5 min in PBS. Tissue sections were then re-fixed in 4% PFA/PBS for 5 min before being washed for 3x 5 min in PBS. Sections were subsequently acetylated by being incubated in a solution containing 100 mM triethanolamine, 1.8 mM HCl and 0.5% acetic anhydride before being washed for 3x 5 min in PBS and permeabilized in a 1% triton/PBS solution for 30 min at RT. Endogenous peroxidase activity was then blocked by incubating in a 0.3% hydrogen peroxide solution for 30 min before washing for 2x 5min in PBS and then finally hybridizing with relevant probes overnight at 64 °C. Bound probes were detected with anti-Digoxigenin and anti-Fluorescein antibodies (Roche) and staining was amplified using a TSA staining Kit (Perkin Elmer (NEL0701001KT)) according to the manufacturer’s instructions. Immunostaining for Iba1 and S100□ was subsequently performed as described for IHC. The mouse *C1qa* probe was validated on tissue from *C1qa* KO mice (a kind gift from M. Botto, Imperial College London) (data not shown) and sense probes for human *C1Q*, and *C3* were found to generate no detectable signal when employed on sequential tissue sections (data not shown).

### Single molecule fluorescence in situ hybridization

14 μm cryosections of mouse brain tissue were generated according to the procedures outlined above for immunohistochemistry with the exception that once extracted mouse brains were post fixed for 24hr in 4% PFA before being washed and placed in a 30% sucrose solution in PBS for storage at 4°C for 24-48hr (as described above). Subsequent processing and sectioning steps were the same. Once generated sections were prepared for in situ hybridization using the procedures outlined by Advanced Cell System’s branched DNA technology (RNAscope) for employing fixed frozen tissue in their multiplexed fluorescent amplification assay (ACD user manual numbers 320535 and 320293). Briefly, sections were washed 3x in PBS before undergoing target retrieval by being placed into boiling 1X retrieval solution (ACD 322000) for 5 min. After washing in distilled water tissue sections were dehydrated by being placed into 100% ethanol. Following this tissue was subsequently permeabilized by treatment with protease III (ACD 322340) for 30 min at 40°C. After tissue preparation was complete hybridization using probes to *C3* (ACD 417841) and *Acta2* (ACD 319531) was carried out along with subsequent steps to amplify and detect signal. All procedures were performed according to the guidelines stipulated in the multiplexed fluorescent amplification assay manual (ACD 320293). Imaging was carried out using a Zeiss LSM 880 confocal microscope and Zen Black 2.3 image acquisition software (Carl Zeiss). Tissue quality was assessed as good by having robust and uniform signal following hybridization with the RNAscope Triplex positive control probes (ACD 320881) and minimal signal with the Triplex negative control (ACD 320871, a probe to DapB a bacterial transcript).

### Quantitative RTPCR

Microdissected mouse and human brain tissue was flash frozen and homogenized in Trizol using a TissueLyser II (Qiagen). Phenol chloroform extraction was then used together with the RNeasy mini kit (QIAGEN) to isolate and purify RNA. RNA quantity was subsequently measured using a Nanodrop (Thermo Scientific), and cDNA was synthesized from all samples by employing Superscript II reverse transcriptase (Thermo Fisher Scientific). After cDNA synthesis, qPCR reactions were assembled for the gene of interest and a housekeeping gene (*Gapdh*) (see key resources table for the sequences of all primers used) using SYBR Green (QIAGEN). Reactions were run on a Rotogene qPCR machine (QIAGEN) and expression levels were compared using the ddCt method with normalization to *Gapdh* levels.

### Immunoblotting

Microdissected mouse and human brain tissue was flash frozen on dry ice. Frozen tissue was then transferred into a triton based lysis buffer (25 mM Hepes, 0.1 M NaCl, 1% Triton X100) containing cOmplete protease inhibitor cocktail (Millipore Sigma). A 5 mm stainless steel bead was subsequently used to homogenize the tissue using the TissueLyser II system (Qiagen) for 2x 5 min at 20 hz. Lysates were then spun for 10 min at 15,000 g in a benchtop centrifuge to pellet insoluble material before the supernatant was removed to a new tube and protein concentration assayed using the BCA protein assay kit (Pierce) according to the manufacturer’s instructions. Protein samples (20 µg) were then mixed with 2x Laemmli buffer (Bio-Rad) before being loaded onto 10% tris-glycine gels and separated for 1 h at 150 V by SDS-PAGE. Separated proteins were transferred onto nitrocellulose membranes (GE Healthcare Amersham) which were subsequently blocked in 5% milk PBS for 1 h at room temperature before being probed with relevant antibodies O/N at 4°C in 5% milk PBS. Membranes were then washed for 3x 5 min in PBS before being incubated with relevant secondary antibodies diluted in 5% milk PBS for 1 h at room temperature. After washing again for 3x 5 min in PBS bands were visualized with luminol-based enhanced chemiluminescence HRP substrate (Super signal west dura, Thermo Fisher Scientific) and the ImageQuant LAS 4000 system (GE healthcare) avoiding saturation of any pixels. To strip membranes and re-probe with a different antibody ReBlot Plus strong antibody stripping solution (EMD Millipore) was employed according to the manufacturer’s instructions prior to re-blocking in 5% milk and staining. For quantification, the intensity of bands generated by staining with synaptic antibodies was normalized to those generated by a β-actin antibody (employed as a loading control) in the same lane. ImageJ software was used to carry out densitometry analysis of all bands, which was performed blind to genotype.

### ELISA assays

For tissue samples, microdissected human brain tissue was flash frozen and lysed to extract proteins using a TissueLyser II system (QIAGEN) and total protein concentration determined using the Pierce BCA protein assay kit (Thermo Fisher Scientific 23225), as described above for immunoblotting. In the case of human CSF and plasma samples, these were acquired, prepared and stored as described above. For subsequent interrogation of complement protein levels all samples were employed in either the C3 (Abcam ab108822 and 108823), iC3b, C1q or CR3 sandwich ELISA’s according to the procedures outlined below. Importantly for every ELISA the same 96 well plate configuration was employed with 2 columns of standards and two dilutions of each sample with technical replicates employed for each dilution. In addition, two common reference samples and four blank wells were included on each plate for calibration and a duplicate of every plate was tested. For both the CSF and plasma samples on each plate the gender balance and average age for each clinical group was kept equal and consistent with the average age of that clinical group as a whole. All assays were run on automated (Bravo liquid handler; Agilent) or semi-automated platforms (VIAFLO; INTEGRA) using novel pipelines and analysis was performed blind to sample type.

### C3 ELISA

5 μg of protein extracted from frozen human tissue, CSF (diluted 1/400 and 1/800 in sample diluent) or plasma (diluted 1/166.6 and 1/333.2 in sample diluent) samples were employed in C3 sandwich and competitive ELISA’s (Abcam 108823 (for CSF and tissue samples) and 108822 (for plasma) respectively). The same lot of both ELISA kits was employed for analysis of all samples and the assay was performed according to the manufacturer’s instructions. In addition, prior to employing this assay tests were carried out to confirm that minimal signal was generated with C3 depleted serum (data not shown).

### iC3b ELISA

16 μg of protein extracted from frozen human tissue, CSF (diluted 1/5 and 1/10 in PBS) or plasma (diluted 1/400 and 1/800 in PBS) samples were employed in an iC3b sandwich ELISA with a monoclonal iC3b antibody (Quidel A209) used for capture and a polyclonal C3 antibody (Dako F0201) used for detection.

Prior to employing this assay, tests were carried out to confirm that minimal signal was generated with C3 depleted serum, that it could detect increased levels of iC3b following *in vitro* complement activation (either through incubation of serum at 4°C for 7 days or zymosan treatment) (Extended data Fig. 1g,h) and that it was selective for iC3b over native C3 and other C3 cleavage fragments (Extended data Fig. 1i).

### C1Q ELISA

CSF (diluted 1/25 and 1/50 in PBS) or plasma (diluted 1/125 or 1/250 in PBS) samples were employed in a C1q sandwich ELISA with a polyclonal rabbit antibody (Abcam AB71940) used for capture and a monoclonal C1q antibody (Annexon M1) conjugated with alkaline phosphatase used for detection. Prior to employing this assay tests were carried out to confirm that minimal signal was generated with C1q depleted serum.

### CR3 ELISA

16 μg of protein extracted from frozen human tissue samples was employed in a human complement receptor 3 (CR3/ITGAM) ELISA (LSBio LS-F11850). The same lot of this kit was employed for the analysis of all samples and the assay was performed according to the manufacturer’s instructions.

### Hemoglobin ELISA

16 μg of protein extracted from frozen human tissue samples was employed in a hemoglobin sandwich ELISA (Abcam Ab157707). The same lot of this kit was employed for the analysis of all samples and the assay was performed according to the manufacturer’s instructions.

### Albumin ELISA

CSF (diluted 1/2000 and 1/4000 in sample diluent) samples were employed in an albumin sandwich ELISA (Abcam ab108788). The same lot of this kit was employed for the analysis of all samples and the assay was performed according to the manufacturer’s instructions.

All non-commercial ELISA assays were found to produce consistent results with an intra-assay coefficient of between 7 and 10% for both the iC3b and C1q ELISA assays based on internal quality control samples.

#### Age adjustment calculation

To control for the effects of normal aging on CSF concentrations of complement proteins in HD patients the CSF concentrations of C3 and iC3b in control (clinically normal) individuals were regressed on the basis of patient age. The regression coefficient generated from this, termed b, was then employed in the below calculation where Y_hdi is the observation of the protein level in Huntington’s disease gene expansion carriers (HDGEC), age_hdi is the corresponding age and mean(age) is the mean age in the control (clinically normal population).

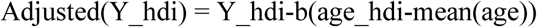

#### CAP score calculation

The CAG age product calculation was initially devised by Penny et al.,^191^ to estimate the progression of HD pathology as a function of both CAG repeat length and the time of exposure to the effects of the expansion and was driven by the observation that the age of clinical onset in HD is strongly influenced by the length of the CAG trinucleotide expansion within the HTT gene. The authors found that an index of this form was a good predictor of striatal pathology in the brains of HD patients at the time of autopsy. Subsequent studies have shown correlations between CAP score and levels of fluid biomarkers, motor and cognitive performance, differences in neuropsychiatric assessments, and structural and molecular imaging markers^132,137,139-141,143,192-197^.

When employed in this manuscript the CAP score is defined as follows,

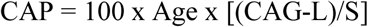

Where CAG is the patient’s CAG repeat length, Age is the patient’s current age at the time of observation, and L and S are constants of 30 and 627 respectively. S is a normalizing constant chosen so that the CAP score is approximately 100 at the patient’s expected age of onset as estimated by Langbehn et al., (Langbehn et al., 2004). L is a scaling constant that anchors CAG length approximately at the lower end of the distribution relevant to HD pathology. Intuitively, L might be thought of as the lower limit of the CAG lengths for which some pathological effect might be expected.

#### In vitro complement activation protocols

Serum samples from control (clinically normal) individuals or serum samples in which C3 and C4 were inactivated were either thawed and maintained at 4°C for 7 days or treated with 10mg/ml of zymosan for 30 min at 37°C. They were then immediately employed in the iC3b ELISA detailed above.

### Operant touchscreen visual discrimination and cognitive flexibility assays

#### Apparatus

Two-choice visual discrimination and cognitive flexibility assays were performed in Bussey-Saksida operant touchscreen chambers, sealed enclosures which have a trapezoidal wall shape structure with a touch sensitive computer screen at the front and a feeder tray at the rear through which a liquid reward can be dispensed (Lafayette Instrument Company model 80614). The chambers were set up with a 1×2 mask, to constrain access to two portions of the screen and operated using ABET II touch software housed on a WhiskerServer Controller. All chambers were configured using the standard touch screen environment file provided by Lafayette Instrument and the hardware was tested prior to running programs using the ‘12-TouchMouseTestLines’ protocol in ABET II and the display test pattern feature in WhiskerServer to ensure that all screens and input/output responses were functioning as proscribed. All chambers were housed in a sound attenuating cupboard.

#### Food Restriction Procedures

To motivate task engagement all animals underwent a diet restriction paradigm prior to testing in which the quantity of food pellets they were allowed access to was restricted. The mice were weighed on a daily basis and a sufficient quantity of pellets provided such that their mass was maintained at approximately 80-85% of their original free feeding weight. Food restriction was begun 1 week prior to training and continued throughout the testing period.

#### Training and shaping tasks

After the mice had reached target bodyweights they were placed in the operant touchscreen chambers and a habituation program was run to familiarize them with collecting liquid reward from the feeder tray. This program involved a tone being played and a light simultaneously being turned on in the feeder tray alongside the presentation of a liquid reward (a solution containing 14% sugar). Reward was presented on a variable schedule in which mice received a total of 60 reinforcers per 60 minute session. All mice underwent two sessions of this feeder habituation on consecutive days.

Following completion of feeder training mice were familiarized with the touchscreen interface by running a ‘must touch’ program in which physical interaction with a random object presented at different locations on the screen was required to initiate the dispensing of a reward. Random geometric shapes consisting of similar dimensions and pixel intensities were presented at variable locations across the base of the screen and physical interaction with the displayed image resulting in it being removed from the screen and a simultaneous dispensing of reward with the same mechanics of tone presentation and illumination of the feeder that were operated during feeder habituation. Images were presented on a variable schedule in which mice were offered the opportunity to complete 60 trials in a 60 minute period. All mice underwent three sessions of this training on consecutive days and by the end all had reached the criteria of completing at least 50 trials in a 60 minute period. The 1×2 mask was not employed during this training.

#### Acquisition phase of visual discrimination learning

In this phase of the task mice were presented with two different visual stimuli of similar size and pixel density which were randomly displayed in the two locations on the screen where access was not restricted by the 1×2 mask. Physical interaction with one of these two stimuli resulted in reward presentation through the same mechanisms outlined in the training section. If, however the mice interacted with the other image presented a different tone was played and a light positioned to illuminate the chamber was switched on for 3 seconds. In addition, no reward was dispensed. A trial session was complete when the mice had performed 60 of these trials or 60 minutes had elapsed. The inter-trial interval was 5 seconds and images were never presented in the same location more than 3 times consecutively. For each trial session the total number of trials performed, the % of those trials in which the rewarded visual presentation was chosen and the latency to collect the reward were recorded. Mice were not allowed to move on to the reversal phase until they had interacted with the rewarded stimuli in 70% of the trials carried out within a session on at least 3 occasions (with one trial session being carried out per day).

#### Reversal phase

In this phase of the task interaction with the previously rewarded stimulus resulted in presentation of the same aversive cues and non-dispense of the liquid reward that interacting with the previously unrewarded image yielded and vice versa. In all other aspects the procedures followed during reversal were identical to those of the acquisition phase. Mice underwent 15 days of reversal testing with a separate session of 60 trials carried out each day regardless of performance in the task.

#### Progressive ratio

Following completion of testing in the reversal phase, mice were transitioned to a progressive ratio task in which one image was displayed in a central location on the screen. At the start of this assessment a single physical interaction with this image resulted in reward being dispensed in the same manner as that described above however after completion of the 1^st^ trial, the image remained on the screen until a 2^nd^ physical interaction had taken place and only then was reward dispensed. This pattern was continued up to the 5^th^ trial where 5 physical interactions were required to elicit a reward. For all subsequent trials after this point 5 physical interactions were required to complete the trial and get the reward. The mice were assessed on how many trials of this nature they were able to complete in a 60 minute period. All mice underwent progressive ratio testing in this format for 3 days. After this point the whole procedure was repeated but this time with a food pellet present in the chamber. Both the total number of trials completed and the mass of food pellet consumed were recorded.

#### Image bias assessment

In order to discern whether mice show a pre-existing preference for either of the two images used in the visual discrimination learning and reversal tests a separate cohort of 4 mo C57BL/6J mice (Jax stock number 000664) were randomly assigned to two groups, one in which the standard stimuli reward associations previously used for testing were employed and one in which these were reversed. The acquisition and reversal phases of the task were then carried out as described above. This testing revealed that there is a small but significant preference for one of the two visual stimuli’s that exists prior to any reinforcement learning taking place and which is maintained throughout testing (Extended data Fig. 8k,l) This might explain the slightly lower than chance performance of all mice on day one of testing but as all of the genotypes assayed received equal numbers of trials in which this stimulus was associated with reward presentation it is likely not responsible for the genotype dependent differences observed.

#### Visual acuity assessment using the optomotor test

Visual acuity was assessed using the optomotor task outlined in Pursky et al., 2004^198^. In brief, each mouse was placed unrestrained on a platform surrounded by four computer monitors which were set to display a rotating cylinder covered with a vertical sine wave grating (set at 100% contrast) that was projected in three-dimensional space. Mice reflexively tracked the grating with head and neck movements that could be documented by an observer. The spatial frequency of the grating was subsequently clamped at the viewing position by repeatedly re-centering the cylinder on the head of the mouse and visual acuity assessed by gradually increasing the frequency until a reflexive optomotor response could no longer be detected. This was designated as the spatial frequency threshold of visual acuity and it was subsequently obtained for each direction of rotation. Previous studies have shown that visual acuity increases markedly between the 2^nd^ and 4^th^ postnatal weeks but remains stable at a spatial frequency of 0.4 cycles/degree after that point in C57BL/6J mice that have not undergone any procedures and which were maintained in similar conditions (Jax stock # 000664)^198^.

### Statistical Analysis

For all statistical analyses, GraphPad Prism 7 and Stata SE software were used. For data generated from mouse models consultation on appropriate statistical tests was carried out with Dr Kush Kapor an associate director of biostatistics at the Harvard Clinical and Translational Science Center. For data generated from CSF and plasma analysis consultations on power calculations and appropriate statistical tests were carried out with Dr Jen Ware, a director of experimental design, and Dr John Warner, a director of biostatistics, at the CHDI foundation.

To determine appropriate sample sizes for the CSF and plasma analysis a pilot study was performed using a small number of CSF and serum samples (sourced from the University of Washington). Based on the effect sizes observed a power analysis was performed using G*Power version 3.1.9.2 (Germany), which determined the number of individuals required to detect the same effect with 80% power at an alpha level of 2.5% (corrected for two primary comparisons using the Bonferroni method).

For all data analysis tests for normality (D’Agostino and Pearson normality test) were performed where appropriate. When comparing two groups either an unpaired two-tailed t-test, or the non-parametric Kolmogorov-Smirnov test were performed as appropriate. When comparing more than two groups one-way ANOVA (fixed effects, omnibus) was used followed by post hoc testing with Tukey’s multiple comparison test with a 95% confidence interval. For all comparisons involving more than two groups and data generated from human CSF and plasma the nonparametric Kruskal-Wallis test was employed followed by post hoc testing with Dunn’s multiple comparison test and a family-wise significance and confidence level of 0.05. The two-tailed nonparametric Spearman correlation test with a confidence interval of 95% was employed to test the significance of relationships between two variables within the CSF and plasma datasets. For the CSF, plasma and tissue extract analysis potentially confounding variables such as age, gender and blood contamination were examined and where appropriate were included as covariates or adjusted for. For the age adjustment calculation linear regression was used to compare the relationship between patient age and analyte concentration in control (clinically normal) individuals. All p values are indicated in both the figure panels and the figure legends.

## References

1. Ross, C.A., et al. Huntington disease: natural history, biomarkers and prospects for therapeutics. Nat Rev Neurol 10, 204–216 (2014).

2. Bates, G.P., et al. Huntington disease. Nat Rev Dis Primers 1, 15005 (2015).

3. A novel gene containing a trinucleotide repeat that is expanded and unstable on Huntington’s disease chromosomes. The Huntington’s Disease Collaborative Research Group. Cell 72, 971–983 (1993).

4. Crook, Z.R. & Housman, D. Huntington’s disease: can mice lead the way to treatment? Neuron 69, 423–435 (2011).

5. Ehrnhoefer, D.E., Butland, S.L., Pouladi, M.A. & Hayden, M.R. Mouse models of Huntington disease: variations on a theme. Dis Model Mech 2, 123–129 (2009).

6. Chan, A.W., et al. Progressive cognitive deficit, motor impairment and striatal pathology in a transgenic Huntington disease monkey model from infancy to adulthood. PLoS One 10, e0122335 (2015).

7. Chang, R., Liu, X., Li, S. & Li, X.J. Transgenic animal models for study of the pathogenesis of Huntington’s disease and therapy. Drug Des Devel Ther 9, 2179–2188 (2015).

8. Espinoza, F.A., et al. Resting-state fMRI dynamic functional network connectivity and associations with psychopathy traits. Neuroimage Clin 24, 101970 (2019).

9. Espinoza, F.A., et al. Whole-Brain Connectivity in a Large Study of Huntington’s Disease Gene Mutation Carriers and Healthy Controls. Brain Connect 8, 166–178 (2018).

10. Gu, X., et al. Pathological cell-cell interactions elicited by a neuropathogenic form of mutant Huntingtin contribute to cortical pathogenesis in HD mice. Neuron 46, 433–444 (2005).

11. Aylward, E.H., et al. Reduced basal ganglia volume associated with the gene for Huntington’s disease in asymptomatic at-risk persons. Neurology 44, 823–828 (1994).

12. Hobbs, N.Z., et al. Onset and progression of pathologic atrophy in Huntington disease: a longitudinal MR imaging study. AJNR Am J Neuroradiol 31, 1036–1041 (2010).

13. Cepeda, C., Cummings, D.M., Andre, V.M., Holley, S.M. & Levine, M.S. Genetic mouse models of Huntington’s disease: focus on electrophysiological mechanisms. ASN Neuro 2, e00033 (2010).

14. Cepeda, C., et al. Rescuing the Corticostriatal Synaptic Disconnection in the R6/2 Mouse Model of Huntington’s Disease: Exercise, Adenosine Receptors and Ampakines. PLoS Curr 2(2010).

15. Cepeda, C., Wu, N., Andre, V.M., Cummings, D.M. & Levine, M.S. The corticostriatal pathway in Huntington’s disease. Prog Neurobiol 81, 253–271 (2007).

16. Lahr, J., et al. Working Memory-Related Effective Connectivity in Huntington’s Disease Patients. Front Neurol 9, 370 (2018).

17. Langfelder, P., et al. Integrated genomics and proteomics define huntingtin CAG length-dependent networks in mice. Nat Neurosci 19, 623–633 (2016).

18. McColgan, P., et al. White matter predicts functional connectivity in premanifest Huntington’s disease. Ann Clin Transl Neurol 4, 106–118 (2017).

19. Wang, N., et al. Neuronal targets for reducing mutant huntingtin expression to ameliorate disease in a mouse model of Huntington’s disease. Nat Med 20, 536–541 (2014).

20. Yan, S., et al. A Huntingtin Knockin Pig Model Recapitulates Features of Selective Neurodegeneration in Huntington’s Disease. Cell 173, 989–1002 e1013 (2018).

21. Raymond, L.A., et al. Pathophysiology of Huntington’s disease: time-dependent alterations in synaptic and receptor function. Neuroscience 198, 252–273 (2011).

22. Veldman, M.B. & Yang, X.W. Molecular insights into cortico-striatal miscommunications in Huntington’s disease. Curr Opin Neurobiol 48, 79–89 (2018).

23. Plotkin, J.L. & Surmeier, D.J. Corticostriatal synaptic adaptations in Huntington’s disease. Curr Opin Neurobiol 33, 53–62 (2015).

24. Hintiryan, H., et al. The mouse cortico-striatal projectome. Nat Neurosci 19, 1100–1114 (2016).

25. Milnerwood, A.J. & Raymond, L.A. Corticostriatal synaptic function in mouse models of Huntington’s disease: early effects of huntingtin repeat length and protein load. J Physiol 585, 817–831 (2007).

26. Poudel, G.R., et al. Longitudinal change in white matter microstructure in Huntington’s disease: The IMAGE-HD study. Neurobiol Dis 74, 406–412 (2015).

27. Tabrizi, S.J., et al. Potential endpoints for clinical trials in premanifest and early Huntington’s disease in the TRACK-HD study: analysis of 24 month observational data. Lancet Neurol 11, 42–53 (2012).

28. Unschuld, P.G., et al. Impaired cortico-striatal functional connectivity in prodromal Huntington’s Disease. Neurosci Lett 514, 204–209 (2012).

29. Polosecki, P., et al. Resting-state connectivity stratifies premanifest Huntington’s disease by longitudinal cognitive decline rate. Sci Rep 10, 1252 (2020).

30. Hong, S., et al. Complement and microglia mediate early synapse loss in Alzheimer mouse models. Science 352, 712–716 (2016).

31. Vasek, M.J., et al. A complement-microglial axis drives synapse loss during virus-induced memory impairment. Nature 534, 538–543 (2016).

32. Norris, G.T., et al. Neuronal integrity and complement control synaptic material clearance by microglia after CNS injury. J Exp Med 215, 1789–1801 (2018).

33. Schafer, D.P., et al. Microglia contribute to circuit defects in Mecp2 null mice independent of microglia-specific loss of Mecp2 expression. Elife 5(2016).

34. Dejanovic, B., et al. Changes in the Synaptic Proteome in Tauopathy and Rescue of Tau-Induced Synapse Loss by C1q Antibodies. Neuron 100, 1322–1336 e1327 (2018).

35. Lui, H., et al. Progranulin Deficiency Promotes Circuit-Specific Synaptic Pruning by Microglia via Complement Activation. Cell 165, 921–935 (2016).

36. Werneburg, S., et al. Targeted Complement Inhibition at Synapses Prevents Microglial Synaptic Engulfment and Synapse Loss in Demyelinating Disease. Immunity 52, 167–182 e167 (2020).

37. Miller, J.R., et al. RNA-Seq of Huntington’s disease patient myeloid cells reveals innate transcriptional dysregulation associated with proinflammatory pathway activation. Hum Mol Genet 25, 2893–2904 (2016).

38. Trager, U., et al. HTT-lowering reverses Huntington’s disease immune dysfunction caused by NFkappaB pathway dysregulation. Brain 137, 819–833 (2014).

39. Trager, U., et al. Characterisation of immune cell function in fragment and full-length Huntington’s disease mouse models. Neurobiol Dis 73, 388–398 (2015).

40. Hensman Moss, D.J., et al. Huntington’s disease blood and brain show a common gene expression pattern and share an immune signature with Alzheimer’s disease. Sci Rep 7, 44849 (2017).

41. Tai, Y.F., et al. Microglial activation in presymptomatic Huntington’s disease gene carriers. Brain 130, 1759–1766 (2007).

42. Politis, M., et al. Increased central microglial activation associated with peripheral cytokine levels in premanifest Huntington’s disease gene carriers. Neurobiol Dis 83, 115–121 (2015).

43. Politis, M., et al. Microglial activation in regions related to cognitive function predicts disease onset in Huntington’s disease: a multimodal imaging study. Hum Brain Mapp 32, 258–270 (2011).

44. Oldham, M.C., Langfelder, P. & Horvath, S. Network methods for describing sample relationships in genomic datasets: application to Huntington’s disease. BMC Syst Biol 6, 63 (2012).

45. Franciosi, S., et al. Age-dependent neurovascular abnormalities and altered microglial morphology in the YAC128 mouse model of Huntington disease. Neurobiol Dis 45, 438–449 (2012).

46. Sapp, E., et al. Early and progressive accumulation of reactive microglia in the Huntington disease brain. J Neuropathol Exp Neurol 60, 161–172 (2001).

47. Simmons, D.A., et al. Ferritin accumulation in dystrophic microglia is an early event in the development of Huntington’s disease. Glia 55, 1074–1084 (2007).

48. Kwan, W., et al. Mutant huntingtin impairs immune cell migration in Huntington disease. J Clin Invest 122, 4737–4747 (2012).

49. Crotti, A., et al. Mutant Huntingtin promotes autonomous microglia activation via myeloid lineage-determining factors. Nat Neurosci 17, 513–521 (2014).

50. Crotti, A. & Glass, C.K. The choreography of neuroinflammation in Huntington’s disease. Trends Immunol 36, 364–373 (2015).

51. Savage, J.C., et al. Microglial physiological properties and interactions with synapses are altered at presymptomatic stages in a mouse model of Huntington’s disease pathology. J Neuroinflammation 17, 98 (2020).

52. Lee, H., et al. Cell Type-Specific Transcriptomics Reveals that Mutant Huntingtin Leads to Mitochondrial RNA Release and Neuronal Innate Immune Activation. Neuron 107, 891–908 e898 (2020).

53. Wilton, D.K. & Stevens, B. The contribution of glial cells to Huntington’s disease pathogenesis. Neurobiol Dis 143, 104963 (2020).

54. Singhrao, S.K., Neal, J.W., Morgan, B.P. & Gasque, P. Increased complement biosynthesis by microglia and complement activation on neurons in Huntington’s disease. Exp Neurol 159, 362–376 (1999).

55. Agus, F., Crespo, D., Myers, R.H. & Labadorf, A. The caudate nucleus undergoes dramatic and unique transcriptional changes in human prodromal Huntington’s disease brain. BMC Med Genomics 12, 137 (2019).

56. Hodges, A., et al. Regional and cellular gene expression changes in human Huntington’s disease brain. Hum Mol Genet 15, 965–977 (2006).

57. Labadorf, A., et al. RNA Sequence Analysis of Human Huntington Disease Brain Reveals an Extensive Increase in Inflammatory and Developmental Gene Expression. PLoS One 10, e0143563 (2015).

58. Pillai, J.A., et al. Clinical severity of Huntington’s disease does not always correlate with neuropathologic stage. Mov Disord 27, 1099–1103 (2012).

59. Vonsattel, J.P., et al. Neuropathological classification of Huntington’s disease. J Neuropathol Exp Neurol 44, 559–577 (1985).

60. Ferrante, R.J., Kowall, N.W. & Richardson, E.P., Jr. Proliferative and degenerative changes in striatal spiny neurons in Huntington’s disease: a combined study using the section-Golgi method and calbindin D28k immunocytochemistry. J Neurosci 11, 3877–3887 (1991).

61. Fourie, C., et al. Differential Changes in Postsynaptic Density Proteins in Postmortem Huntington’s Disease and Parkinson’s Disease Human Brains. J Neurodegener Dis 2014, 938530 (2014).

62. Reis, E.S., Mastellos, D.C., Hajishengallis, G. & Lambris, J.D. New insights into the immune functions of complement. Nat Rev Immunol 19, 503–516 (2019).

63. Noris, M. & Remuzzi, G. Overview of complement activation and regulation. Semin Nephrol 33, 479–492 (2013).

64. Halliday, G.M., et al. Regional specificity of brain atrophy in Huntington’s disease. Exp Neurol 154, 663–672 (1998).

65. Wakai, M., Takahashi, A. & Hashizume, Y. A histometrical study on the globus pallidus in Huntington’s disease. J Neurol Sci 119, 18–27 (1993).

66. Singh-Bains, M.K., Waldvogel, H.J. & Faull, R.L. The role of the human globus pallidus in Huntington’s disease. Brain Pathol 26, 741–751 (2016).

67. Lei, W., et al. Confocal laser scanning microscopy and ultrastructural study of VGLUT2 thalamic input to striatal projection neurons in rats. J Comp Neurol 521, 1354–1377 (2013).

68. Raju, D.V., Shah, D.J., Wright, T.M., Hall, R.A. & Smith, Y. Differential synaptology of vGluT2-containing thalamostriatal afferents between the patch and matrix compartments in rats. J Comp Neurol 499, 231–243 (2006).

69. Ding, J.B., Oh, W.J., Sabatini, B.L. & Gu, C. Semaphorin 3E-Plexin-D1 signaling controls pathway-specific synapse formation in the striatum. Nat Neurosci 15, 215–223 (2011).

70. Gray, M., et al. Full-length human mutant huntingtin with a stable polyglutamine repeat can elicit progressive and selective neuropathogenesis in BACHD mice. J Neurosci 28, 6182–6195 (2008).

71. Menalled, L.B., et al. Comprehensive behavioral and molecular characterization of a new knock-in mouse model of Huntington’s disease: zQ175. PLoS One 7, e49838 (2012).

72. Heikkinen, T., et al. Characterization of neurophysiological and behavioral changes, MRI brain volumetry and 1H MRS in zQ175 knock-in mouse model of Huntington’s disease. PLoS One 7, e50717 (2012).

73. Carty, N., et al. Characterization of HTT inclusion size, location, and timing in the zQ175 mouse model of Huntington’s disease: an in vivo high-content imaging study. PLoS One 10, e0123527 (2015).

74. Indersmitten, T., Tran, C.H., Cepeda, C. & Levine, M.S. Altered excitatory and inhibitory inputs to striatal medium-sized spiny neurons and cortical pyramidal neurons in the Q175 mouse model of Huntington’s disease. J Neurophysiol 113, 2953–2966 (2015).

75. Plotkin, J.L., et al. Impaired TrkB receptor signaling underlies corticostriatal dysfunction in Huntington’s disease. Neuron 83, 178–188 (2014).

76. Zhang, C., et al. Abnormal Brain Development in Huntington’ Disease Is Recapitulated in the zQ175 Knock-In Mouse Model. Cereb Cortex Commun 1, tgaa044 (2020).

77. Beaumont, V., et al. Phosphodiesterase 10A Inhibition Improves Cortico-Basal Ganglia Function in Huntington’s Disease Models. Neuron 92, 1220–1237 (2016).

78. Vezzoli, E., et al. Inhibiting pathologically active ADAM10 rescues synaptic and cognitive decline in Huntington’s disease. J Clin Invest 129, 2390–2403 (2019).

79. McKinstry, S.U., et al. Huntingtin is required for normal excitatory synapse development in cortical and striatal circuits. J Neurosci 34, 9455–9472 (2014).

80. Hisano, S., et al. Regional expression of a gene encoding a neuron-specific Na(+)-dependent inorganic phosphate cotransporter (DNPI) in the rat forebrain. Brain Res Mol Brain Res 83, 34–43 (2000).

81. Fremeau, R.T. Jr.,, et al. The expression of vesicular glutamate transporters defines two classes of excitatory synapse. Neuron 31, 247–260 (2001).

82. Fremeau, R.T. Jr.,, Voglmaier, S., Seal, R.P. & Edwards, R.H. VGLUTs define subsets of excitatory neurons and suggest novel roles for glutamate. Trends Neurosci 27, 98–103 (2004).

83. Hur, E.E. & Zaborszky, L. Vglut2 afferents to the medial prefrontal and primary somatosensory cortices: a combined retrograde tracing in situ hybridization study [corrected]. J Comp Neurol 483, 351–373 (2005).

84. Bacci, J.J., Kachidian, P., Kerkerian-Le Goff, L. & Salin, P. Intralaminar thalamic nuclei lesions: widespread impact on dopamine denervation-mediated cellular defects in the rat basal ganglia. J Neuropathol Exp Neurol 63, 20–31 (2004).

85. Parievsky, A., et al. Differential electrophysiological and morphological alterations of thalamostriatal and corticostriatal projections in the R6/2 mouse model of Huntington’s disease. Neurobiol Dis 108, 29–44 (2017).

86. Deng, Y.P., Wong, T., Wan, J.Y. & Reiner, A. Differential loss of thalamostriatal and corticostriatal input to striatal projection neuron types prior to overt motor symptoms in the Q140 knock-in mouse model of Huntington’s disease. Front Syst Neurosci 8, 198 (2014).

87. Octeau, J.C., et al. An Optical Neuron-Astrocyte Proximity Assay at Synaptic Distance Scales. Neuron 98, 49–66 e49 (2018).

88. Taft, C.E. & Turrigiano, G.G. PSD-95 promotes the stabilization of young synaptic contacts. Philos Trans R Soc Lond B Biol Sci 369, 20130134 (2014).

89. van den Bogaard, S.J., et al. Early atrophy of pallidum and accumbens nucleus in Huntington’s disease. J Neurol 258, 412–420 (2011).

90. Fonseca, M.I., et al. Cell-specific deletion of C1qa identifies microglia as the dominant source of C1q in mouse brain. J Neuroinflammation 14, 48 (2017).

91. Saunders, A., et al. Molecular Diversity and Specializations among the Cells of the Adult Mouse Brain. Cell 174, 1015–1030 e1016 (2018).

92. Stevens, B., et al. The classical complement cascade mediates CNS synapse elimination. Cell 131, 1164–1178 (2007).

93. Sapp, E., et al. Protein changes in synaptosomes of Huntington’s disease knock-in mice are dependent on age and brain region. Neurobiol Dis 141, 104950 (2020).

94. Zhao, X., et al. Noninflammatory Changes of Microglia Are Sufficient to Cause Epilepsy. Cell Rep 22, 2080–2093 (2018).

95. Reichert, F. & Rotshenker, S. Galectin-3 (MAC-2) Controls Microglia Phenotype Whether Amoeboid and Phagocytic or Branched and Non-phagocytic by Regulating the Cytoskeleton. Front Cell Neurosci 13, 90 (2019).

96. Schafer, D.P., et al. Microglia sculpt postnatal neural circuits in an activity and complement-dependent manner. Neuron 74, 691–705 (2012).

97. Burgold, J., et al. Cortical circuit alterations precede motor impairments in Huntington’s disease mice. Sci Rep 9, 6634 (2019).

98. Schippling, S., et al. Abnormal motor cortex excitability in preclinical and very early Huntington’s disease. Biol Psychiatry 65, 959–965 (2009).

99. Sasaki, Y., et al. Selective expression of Gi/o-coupled ATP receptor P2Y12 in microglia in rat brain. Glia 44, 242–250 (2003).

100. Bennett, M.L., et al. New tools for studying microglia in the mouse and human CNS. Proc Natl Acad Sci U S A 113, E1738–1746 (2016).

101. Schafer, D.P., Lehrman, E.K., Heller, C.T. & Stevens, B. An engulfment assay: a protocol to assess interactions between CNS phagocytes and neurons. J Vis Exp (2014).

102. Ebihara, T., Kawabata, I., Usui, S., Sobue, K. & Okabe, S. Synchronized formation and remodeling of postsynaptic densities: long-term visualization of hippocampal neurons expressing postsynaptic density proteins tagged with green fluorescent protein. J Neurosci 23, 2170–2181 (2003).

103. Chung, W.S., et al. Astrocytes mediate synapse elimination through MEGF10 and MERTK pathways. Nature 504, 394–400 (2013).

104. Tong, X., et al. Astrocyte Kir4.1 ion channel deficits contribute to neuronal dysfunction in Huntington’s disease model mice. Nat Neurosci 17, 694–703 (2014).

105. Jiang, R., Diaz-Castro, B., Looger, L.L. & Khakh, B.S. Dysfunctional Calcium and Glutamate Signaling in Striatal Astrocytes from Huntington’s Disease Model Mice. J Neurosci 36, 3453–3470 (2016).

106. Khakh, B.S., et al. Unravelling and Exploiting Astrocyte Dysfunction in Huntington’s Disease. Trends Neurosci 40, 422–437 (2017).

107. Bradford, J., et al. Expression of mutant huntingtin in mouse brain astrocytes causes age-dependent neurological symptoms. Proc Natl Acad Sci U S A 106, 22480–22485 (2009).

108. Huang, B., et al. Mutant huntingtin downregulates myelin regulatory factor-mediated myelin gene expression and affects mature oligodendrocytes. Neuron 85, 1212–1226 (2015).

109. Petkau, T.L., et al. Mutant huntingtin expression in microglia is neither required nor sufficient to cause the Huntington’s disease-like phenotype in BACHD mice. Hum Mol Genet 28, 1661–1670 (2019).

110. Wood, T.E., et al. Mutant huntingtin reduction in astrocytes slows disease progression in the BACHD conditional Huntington’s disease mouse model. Hum Mol Genet 28, 487–500 (2019).

111. Faideau, M., et al. In vivo expression of polyglutamine-expanded huntingtin by mouse striatal astrocytes impairs glutamate transport: a correlation with Huntington’s disease subjects. Hum Mol Genet 19, 3053–3067 (2010).

112. Ferrari Bardile, C., et al. Intrinsic mutant HTT-mediated defects in oligodendroglia cause myelination deficits and behavioral abnormalities in Huntington disease. Proc Natl Acad Sci U S A 116, 9622–9627 (2019).

113. Gaboriaud, C., et al. The crystal structure of the globular head of complement protein C1q provides a basis for its versatile recognition properties. J Biol Chem 278, 46974–46982 (2003).

114. Venkatraman Girija, U., et al. Structural basis of the C1q/C1s interaction and its central role in assembly of the C1 complex of complement activation. Proc Natl Acad Sci U S A 110, 13916–13920 (2013).

115. Almitairi, J.O.M., et al. Structure of the C1r-C1s interaction of the C1 complex of complement activation. Proc Natl Acad Sci U S A 115, 768–773 (2018).

116. Vukojicic, A., et al. The Classical Complement Pathway Mediates Microglia-Dependent Remodeling of Spinal Motor Circuits during Development and in SMA. Cell Rep 29, 3087–3100 e3087 (2019).

117. Lansita, J.A., et al. Nonclinical Development of ANX005: A Humanized Anti-C1q Antibody for Treatment of Autoimmune and Neurodegenerative Diseases. Int J Toxicol 36, 449–462 (2017).

118. Poduslo, J.F., Curran, G.L. & Berg, C.T. Macromolecular permeability across the blood-nerve and blood-brain barriers. Proc Natl Acad Sci U S A 91, 5705–5709 (1994).

119. Zuchero, Y.J., et al. Discovery of Novel Blood-Brain Barrier Targets to Enhance Brain Uptake of Therapeutic Antibodies. Neuron 89, 70–82 (2016).

120. Martinez-Horta, S., et al. Impaired face-like object recognition in premanifest Huntington’s disease. Cortex 123, 162–172 (2020).

121. Scahill, R.I., et al. Biological and clinical characteristics of gene carriers far from predicted onset in the Huntington’s disease Young Adult Study (HD-YAS): a cross-sectional analysis. Lancet Neurol 19, 502–512 (2020).

122. Langley, C., et al. Fronto-striatal circuits for cognitive flexibility in far from onset Huntington’s disease: evidence from the Young Adult Study. J Neurol Neurosurg Psychiatry 92, 143–149 (2021).

123. Curtin, P.C., et al. Cognitive Training at a Young Age Attenuates Deficits in the zQ175 Mouse Model of HD. Front Behav Neurosci 9, 361 (2015).

124. Piiponniemi, T.O., et al. Impaired Performance of the Q175 Mouse Model of Huntington’s Disease in the Touch Screen Paired Associates Learning Task. Front Behav Neurosci 12, 226 (2018).

125. Oakeshott, S., et al. Deficits in a Simple Visual Go/No-go Discrimination Task in Two Mouse Models of Huntington’s Disease. PLoS Curr 5(2013).

126. Deng, Y., Wang, H., Joni, M., Sekhri, R. & Reiner, A. Progression of basal ganglia pathology in heterozygous Q175 knock-in Huntington’s disease mice. J Comp Neurol 529, 1327–1371 (2021).

127. Luykx, J.J., et al. A common variant in ERBB4 regulates GABA concentrations in human cerebrospinal fluid. Neuropsychopharmacology 37, 2088–2092 (2012).

128. Reiber, H. Dynamics of brain-derived proteins in cerebrospinal fluid. Clin Chim Acta 310, 173–186 (2001).

129. Mouton-Barbosa, E., et al. In-depth exploration of cerebrospinal fluid by combining peptide ligand library treatment and label-free protein quantification. Mol Cell Proteomics 9, 1006–1021 (2010).

130. Higginbotham, L., et al. Integrated proteomics reveals brain-based cerebrospinal fluid biomarkers in asymptomatic and symptomatic Alzheimer’s disease. Sci Adv 6(2020).

131. Fang, Q., et al. Brain-specific proteins decline in the cerebrospinal fluid of humans with Huntington disease. Mol Cell Proteomics 8, 451–466 (2009).

132. Tabrizi, S.J., et al. Biological and clinical manifestations of Huntington’s disease in the longitudinal TRACK-HD study: cross-sectional analysis of baseline data. Lancet Neurol 8, 791–801 (2009).

133. Byrne, L.M., et al. Neurofilament light protein in blood as a potential biomarker of neurodegeneration in Huntington’s disease: a retrospective cohort analysis. Lancet Neurol 16, 601–609 (2017).

134. Wolf, R.C., et al. Brain activation and functional connectivity in premanifest Huntington’s disease during states of intrinsic and phasic alertness. Hum Brain Mapp 33, 2161–2173 (2012).

135. Paulsen, J.S., et al. fMRI biomarker of early neuronal dysfunction in presymptomatic Huntington’s Disease. AJNR Am J Neuroradiol 25, 1715–1721 (2004).

136. Langbehn, D.R., Hayden, M.R., Paulsen, J.S. & and the, P.-H.D.I.o.t.H.S.G. CAG-repeat length and the age of onset in Huntington disease (HD): a review and validation study of statistical approaches. Am J Med Genet B Neuropsychiatr Genet 153B, 397–408 (2010).

137. Harrington, D.L., et al. Cognitive domains that predict time to diagnosis in prodromal Huntington disease. J Neurol Neurosurg Psychiatry 83, 612–619 (2012).

138. Faria, A.V., et al. Linking white matter and deep gray matter alterations in premanifest Huntington disease. Neuroimage Clin 11, 450–460 (2016).

139. Phillips, O.R., et al. Major Superficial White Matter Abnormalities in Huntington’s Disease. Front Neurosci 10, 197 (2016).

140. Chen, L., et al. Altered brain iron content and deposition rate in Huntington’s disease as indicated by quantitative susceptibility MRI. J Neurosci Res 97, 467–479 (2019).

141. Warner, J.H. & Sampaio, C. Modeling Variability in the Progression of Huntington’s Disease A Novel Modeling Approach Applied to Structural Imaging Markers from TRACK-HD. CPT Pharmacometrics Syst Pharmacol 5, 437–445 (2016).

142. Zhang, Y., et al. Indexing disease progression at study entry with individuals at-risk for Huntington disease. Am J Med Genet B Neuropsychiatr Genet 156B, 751–763 (2011).

143. van Bergen, J.M., et al. Quantitative Susceptibility Mapping Suggests Altered Brain Iron in Premanifest Huntington Disease. AJNR Am J Neuroradiol 37, 789–796 (2016).

144. Kamitaki, N., et al. Complement genes contribute sex-biased vulnerability in diverse disorders. Nature 582, 577–581 (2020).

145. Daborg, J., et al. Cerebrospinal fluid levels of complement proteins C3, C4 and CR1 in Alzheimer’s disease. J Neural Transm (Vienna) 119, 789–797 (2012).

146. Gaya da Costa, M., et al. Age and Sex-Associated Changes of Complement Activity and Complement Levels in a Healthy Caucasian Population. Front Immunol 9, 2664 (2018).

147. Ritchie, R.F., Palomaki, G.E., Neveux, L.M. & Navolotskaia, O. Reference distributions for complement proteins C3 and C4: a comparison of a large cohort to the world’s literature. J Clin Lab Anal 18, 9–13 (2004).

148. Dalrymple, A., et al. Proteomic profiling of plasma in Huntington’s disease reveals neuroinflammatory activation and biomarker candidates. J Proteome Res 6, 2833–2840 (2007).

149. Ping, L., et al. Global quantitative analysis of the human brain proteome in Alzheimer’s and Parkinson’s Disease. Sci Data 5, 180036 (2018).

150. Gaboriaud, C., Frachet, P., Thielens, N.M. & Arlaud, G.J. The human c1q globular domain: structure and recognition of non-immune self ligands. Front Immunol 2, 92 (2011).

151. Brier, S., et al. Mapping surface accessibility of the C1r/C1s tetramer by chemical modification and mass spectrometry provides new insights into assembly of the human C1 complex. J Biol Chem 285, 32251–32263 (2010).

152. Ugurlar, D., et al. Structures of C1-IgG1 provide insights into how danger pattern recognition activates complement. Science 359, 794–797 (2018).

153. Litvinchuk, A., et al. Complement C3aR Inactivation Attenuates Tau Pathology and Reverses an Immune Network Deregulated in Tauopathy Models and Alzheimer’s Disease. Neuron 100, 1337–1353 e1335 (2018).

154. Wu, T., et al. Complement C3 Is Activated in Human AD Brain and Is Required for Neurodegeneration in Mouse Models of Amyloidosis and Tauopathy. Cell Rep 28, 2111–2123 e2116 (2019).

155. Ding, J., Peterson, J.D. & Surmeier, D.J. Corticostriatal and thalamostriatal synapses have distinctive properties. J Neurosci 28, 6483–6492 (2008).

156. Lehrman, E.K., et al. CD47 Protects Synapses from Excess Microglia-Mediated Pruning during Development. Neuron 100, 120–134 e126 (2018).

157. Estrada-Sanchez, A.M., et al. Cortical efferents lacking mutant huntingtin improve striatal neuronal activity and behavior in a conditional mouse model of Huntington’s disease. J Neurosci 35, 4440–4451 (2015).

158. Featherstone, R.E. & McDonald, R.J. Dorsal striatum and stimulus-response learning: lesions of the dorsolateral, but not dorsomedial, striatum impair acquisition of a simple discrimination task. Behav Brain Res 150, 15–23 (2004).

159. Crapser, J.D., et al. Microglial depletion prevents extracellular matrix changes and striatal volume reduction in a model of Huntington’s disease. Brain 143, 266–288 (2020).

160. Zeun, P., Scahill, R.I., Tabrizi, S.J. & Wild, E.J. Fluid and imaging biomarkers for Huntington’s disease. Mol Cell Neurosci 97, 67–80 (2019).

161. Silajdzic, E. & Bjorkqvist, M. A Critical Evaluation of Wet Biomarkers for Huntington’s Disease: Current Status and Ways Forward. J Huntingtons Dis 7, 109–135 (2018).

162. Byrne, L.M., et al. Evaluation of mutant huntingtin and neurofilament proteins as potential markers in Huntington’s disease. Sci Transl Med 10(2018).

163. Merel, J., Shababo, B., Naka, A., Adesnik, H. & Paninski, L. Bayesian methods for event analysis of intracellular currents. J Neurosci Methods 269, 21–32 (2016).

164. Coxon, A., et al. A novel role for the beta 2 integrin CD11b/CD18 in neutrophil apoptosis: a homeostatic mechanism in inflammation. Immunity 5, 653–666 (1996).

165. Truett, G.E., et al. Preparation of PCR-quality mouse genomic DNA with hot sodium hydroxide and tris (HotSHOT). Biotechniques 29, 52, 54 (2000).

166. Mason, S.L., et al. Predicting clinical diagnosis in Huntington’s disease: An imaging polymarker. Ann Neurol 83, 532–543 (2018).

167. Langbehn, D.R., et al. A new model for prediction of the age of onset and penetrance for Huntington’s disease based on CAG length. Clin Genet 65, 267–277 (2004).

168. Fanali, G., et al. Human serum albumin: from bench to bedside. Mol Aspects Med 33, 209–290 (2012).

169. Frick, E. & Scheid-Seydel, L. [Exchange processes between plasma and cerebrospinal fluid examined with radio-iodine labeled albumin]. Klin Wochenschr 36, 66–69 (1958).

170. Hofmann, G. & Leupold-Lowenthal, H. [Studies on the blood-brain and the blood-cerebrospinal fluid barrier. I. Penetration of radioiodine labeled albumin into the choroid plexus]. Wien Z Nervenheilkd Grenzgeb 12, 165–170 (1955).

171. Fishman, R.A. Exchange of albumin between plasma and cerebrospinal fluid. Am J Physiol 175, 96–98 (1953).

172. Poduslo, J.F., et al. Altered blood-nerve barrier in experimental lead neuropathy assessed by changes in endoneurial albumin concentration. J Neurosci 2, 1507–1514 (1982).

173. Seyfert, S., Faulstich, A. & Marx, P. What determines the CSF concentrations of albumin and plasma-derived IgG? J Neurol Sci 219, 31–33 (2004).

174. Huang, Y.C., et al. Increased prothrombin, apolipoprotein A-IV, and haptoglobin in the cerebrospinal fluid of patients with Huntington’s disease. PLoS One 6, e15809 (2011).

175. Rub, U., et al. Huntington’s disease (HD): the neuropathology of a multisystem neurodegenerative disorder of the human brain. Brain Pathol 26, 726–740 (2016).

176. McGonigal, R., et al. C1q-targeted inhibition of the classical complement pathway prevents injury in a novel mouse model of acute motor axonal neuropathy. Acta Neuropathol Commun 4, 23 (2016).

177. Jiao, H., et al. Subretinal macrophages produce classical complement activator C1q leading to the progression of focal retinal degeneration. Mol Neurodegener 13, 45 (2018).

178. Holden, S.S., et al. Complement factor C1q mediates sleep spindle loss and epileptic spikes after mild brain injury. Science 373, eabj2685 (2021).

179. Absinta, M., et al. A lymphocyte-microglia-astrocyte axis in chronic active multiple sclerosis. Nature 597, 709–714 (2021).

180. Pluta, S., et al. A direct translaminar inhibitory circuit tunes cortical output. Nat Neurosci 18, 1631–1640 (2015).

181. Porta, E.A. Pigments in aging: an overview. Ann N Y Acad Sci 959, 57–65 (2002).

182. Eichhoff, G., Busche, M.A. & Garaschuk, O. In vivo calcium imaging of the aging and diseased brain. Eur J Nucl Med Mol Imaging 35 Suppl 1, S99–106 (2008).

183. Sierra, A., Gottfried-Blackmore, A.C., McEwen, B.S. & Bulloch, K. Microglia derived from aging mice exhibit an altered inflammatory profile. Glia 55, 412–424 (2007).

184. Hong, S., Wilton, D.K., Stevens, B. & Richardson, D.S. Structured Illumination Microscopy for the Investigation of Synaptic Structure and Function. Methods Mol Biol 1538, 155–167 (2017).

185. Greer, P.L., et al. The Angelman Syndrome protein Ube3A regulates synapse development by ubiquitinating arc. Cell 140, 704–716 (2010).

186. Margolis, S.S., et al. EphB-mediated degradation of the RhoA GEF Ephexin5 relieves a developmental brake on excitatory synapse formation. Cell 143, 442–455 (2010).

187. Micheva, K.D. & Smith, S.J. Array tomography: a new tool for imaging the molecular architecture and ultrastructure of neural circuits. Neuron 55, 25–36 (2007).

188. Ross, S.E., et al. Loss of inhibitory interneurons in the dorsal spinal cord and elevated itch in Bhlhb5 mutant mice. Neuron 65, 886–898 (2010).

189. Waldvogel, H.J., et al. The collection and processing of human brain tissue for research. Cell Tissue Bank 9, 169–179 (2008).

190. Liddelow, S.A., et al. Neurotoxic reactive astrocytes are induced by activated microglia. Nature 541, 481–487 (2017).

191. Penney, J.B. Jr.,, Vonsattel, J.P., MacDonald, M.E., Gusella, J.F. & Myers, R.H. CAG repeat number governs the development rate of pathology in Huntington’s disease. Ann Neurol 41, 689–692 (1997).

192. Vez, S., et al. Auditory time perception in Huntington’s disease. Neuropsychologia 119, 247–252 (2018).

193. Vinther-Jensen, T., et al. Selected CSF biomarkers indicate no evidence of early neuroinflammation in Huntington disease. Neurol Neuroimmunol Neuroinflamm 3, e287 (2016).

194. Novak, M.J., et al. White matter integrity in premanifest and early Huntington’s disease is related to caudate loss and disease progression. Cortex 52, 98–112 (2014).

195. Phillips, O., et al. Tractography of the corpus callosum in Huntington’s disease. PLoS One 8, e73280 (2013).

196. Di Paola, M., et al. Multimodal MRI analysis of the corpus callosum reveals white matter differences in presymptomatic and early Huntington’s disease. Cereb Cortex 22, 2858–2866 (2012).

197. Matusch, A., et al. Cross sectional PET study of cerebral adenosine A(1) receptors in premanifest and manifest Huntington’s disease. Eur J Nucl Med Mol Imaging 41, 1210–1220 (2014).

198. Prusky, G.T., Alam, N.M., Beekman, S. & Douglas, R.M. Rapid quantification of adult and developing mouse spatial vision using a virtual optomotor system. Invest Ophthalmol Vis Sci 45, 4611–4616 (2004).

