## Supplemental tables and extended data figures for "Microglia Mediate Early Corticostriatal Synapse Loss and Cognitive Dysfunction in Huntington’s Disease Through Complement-Dependent Mechanisms"

#### Supplemental Table 1

| Study cohort | Sample ID | Sex | Age | HD Category | CAG high | CAG low |
|---|---|---|---|---|---|---|
| HDClarity | EMHD1 | male | 54.4 | early HD | 43 | 17 |
| HDClarity | EMHD2 | female | 50.5 | early HD | 43 | 17 |
| HDClarity | EMHD3 | female | 60.3 | early HD | 39 | 15 |
| HDClarity | EMHD4 | male | 69 | early HD | 41 | 15 |
| HDClarity | EMHD5 | male | 39 | early HD | 42 | 18 |
| HDClarity | EMHD6 | female | 52.3 | early HD | 43 | 10 |
| HDClarity | EMHD7 | male | 55.8 | early HD | 43 | 17 |
| HDClarity | EMHD8 | male | 63.9 | early HD | 43 | 23 |
| HDClarity | EMHD9 | male | 42.3 | early HD | 44 | 25 |
| HDClarity | EMHD10 | female | 58.5 | early HD | 44 | 23 |
| HDClarity | EMHD11 | female | 58.6 | early HD | 42 | 19 |
| HDClarity | EMHD12 | male | 50.9 | early HD | 44 | 17 |
| HDClarity | EMHD13 | male | 67.2 | early HD | 41 | 21 |
| HDClarity | EMHD14 | male | 52.8 | early HD | 42 | 18 |
| HDClarity | EMHD15 | female | 45.8 | early HD | 44 | 10 |
| HDClarity | EMHD16 | male | 42.3 | early HD | 43 | 17 |
| HDClarity | EMHD17 | male | 45.9 | early HD | 43 | 17 |
| HDClarity | EMHD18 | female | 55.4 | early HD | 45 | 22 |
| HDClarity | EMHD19 | female | 63.2 | early HD | 41 | 17 |
| HDClarity | EMHD20 | male | 64.1 | early HD | 45 | 18 |
| HDClarity | EMHD21 | male | 33.5 | early HD | 50 | 18 |
| HDClarity | EMHD22 | female | 47.5 | early HD | 44 | 18 |
| HDClarity | EMHD23 | female | 55.2 | early HD | 40 | 17 |
| HDClarity | EMHD24 | male | 57.8 | early HD | 42 | 18 |
| HDClarity | EMHD25 | male | 63.5 | early HD | 41 | 17 |
| HDClarity | EMHD26 | female | 70 | early HD | 41 | 17 |
| HDClarity | EMHD27 | female | 40.9 | early HD | 45 | 17 |
| HDClarity | EMHD28 | male | 47.3 | early HD | 43 | 18 |
| HDClarity | EMHD29 | male | 52.6 | early HD | 37 | 17 |
| HDClarity | EMHD30 | female | 58.9 | early HD | 41 | 18 |
| HDClarity | EMHD31 | female | 60 | early HD | 41 | 20 |
| HDClarity | EMHD32 | male | 69.3 | early HD | 42 | 15 |
| HDClarity | EPMHD1 | female | 26.4 | early pre-manifest HD | 41 | 15 |
| HDClarity | EPMHD2 | male | 51.5 | early pre-manifest HD | 40 | 17 |
| HDClarity | EPMHD3 | male | 33.4 | early pre-manifest HD | 42 | 20 |
| HDClarity | EPMHD4 | female | 47 | early pre-manifest HD | 40 | 17 |
| HDClarity | EPMHD5 | female | 52.6 | early pre-manifest HD | 40 | 18 |
| HDClarity | EPMHD6 | female | 35 | early pre-manifest HD | 41 | 18 |
| HDClarity | EPMHD7 | male | 25.3 | early pre-manifest HD | 41 | 17 |
| HDClarity | EPMHD8 | female | 29.1 | early pre-manifest HD | 42 | 17 |

| | | | | | | |
|---|---|---|---|---|---|---|
| HDClarity | EPMHD9 | male | 25.9 | early pre-manifest HD | 40 | 21 |
| HDClarity | EPMHD10 | female | 31 | early pre-manifest HD | 43 | 17 |
| HDClarity | EPMHD11 | female | 33.8 | early pre-manifest HD | 42 | 22 |
| HDClarity | EPMHD12 | female | 34.8 | early pre-manifest HD | 41 | 16 |
| HDClarity | EPMHD13 | male | 37 | early pre-manifest HD | 42 | 19 |
| HDClarity | LPMHD1 | male | 41 | late pre-manifest HD | 42 | 17 |
| HDClarity | LPMHD2 | female | 33.7 | late pre-manifest HD | 45 | 15 |
| HDClarity | LPMHD3 | male | 22.7 | late pre-manifest HD | 47 | 17 |
| HDClarity | LPMHD4 | female | 64.2 | late pre-manifest HD | 41 | 17 |
| HDClarity | LPMHD5 | female | 49.1 | late pre-manifest HD | 43 | 20 |
| HDClarity | LPMHD6 | male | 57 | late pre-manifest HD | 41 | 17 |
| HDClarity | LPMHD7 | male | 31.4 | late pre-manifest HD | 44 | 21 |
| HDClarity | LPMHD8 | female | 24.5 | late pre-manifest HD | 47 | 22 |
| HDClarity | LPMHD9 | male | 54.4 | late pre-manifest HD | 45 | 18 |
| HDClarity | LPMHD10 | female | 41.8 | late pre-manifest HD | 46 | 20 |
| HDClarity | LPMHD11 | female | 47.1 | late pre-manifest HD | 42 | 21 |
| HDClarity | LPMHD12 | male | 53.6 | late pre-manifest HD | 41 | 17 |
| HDClarity | LPMHD13 | male | 27.4 | late pre-manifest HD | 45 | 18 |
| HDClarity | LPMHD14 | female | 43.6 | late pre-manifest HD | 44 | 17 |
| HDClarity | LPMHD15 | male | 45.7 | late pre-manifest HD | 44 | 19 |
| HDClarity | LPMHD16 | female | 50.6 | late pre-manifest HD | 41 | 21 |
| HDClarity | LPMHD17 | female | 29.9 | late pre-manifest HD | 44 | 15 |
| HDClarity | LPMHD18 | male | 46.8 | late pre-manifest HD | 43 | 15 |
| HDClarity | LPMHD19 | male | 47.4 | late pre-manifest HD | 41 | 18 |
| HDClarity | HC1 | female | 48.2 | healthy control | 21 | 18 |
| HDClarity | HC2 | female | 52.8 | healthy control | 24 | 22 |
| HDClarity | HC3 | male | 62 | healthy control | 17 | 17 |
| HDClarity | HC4 | female | 29.2 | healthy control | 22 | 14 |
| HDClarity | HC5 | male | 54.6 | healthy control | 24 | 17 |
| HDClarity | HC6 | male | 33.5 | healthy control | 24 | 17 |
| HDClarity | HC7 | male | 71.9 | healthy control | 17 | 17 |
| HDClarity | HC8 | male | 52 | healthy control | 17 | 9 |
| HDClarity | HC9 | female | 43.2 | healthy control | 20 | 18 |
| HDClarity | HC10 | female | 53.2 | healthy control | 22 | 19 |
| HDClarity | HC11 | male | 53.7 | healthy control | 16 | 16 |
| HDClarity | HC12 | male | 74.6 | healthy control | 17 | 15 |
| HDClarity | HC13 | female | 58.4 | healthy control | 17 | 15 |
| HDClarity | HC14 | female | 39.6 | healthy control | 18 | 9 |
| HDClarity | HC15 | male | 26.5 | healthy control | 20 | 16 |
| HDClarity | HC16 | female | 49 | healthy control | 18 | 17 |
| HDClarity | HC17 | female | 28.9 | healthy control | 18 | 17 |
| HDClarity | HC18 | male | 48.3 | healthy control | 15 | 15 |
| HDClarity | HC19 | male | 50.6 | healthy control | 17 | 15 |

| | | | | | | |
|---|---|---|---|---|---|---|
| HDClarity | HC20 | female | 53.4 | healthy control | 12 | 10 |
| HDClarity | HC21 | male | 54 | healthy control | 21 | 17 |
| HDClarity | HC22 | female | 61 | healthy control | 24 | 20 |
| HDClarity | HC23 | male | 35.4 | healthy control | 19 | 15 |
| HDClarity | HC24 | female | 46.8 | healthy control | 17 | 17 |
| HDClarity | HC25 | male | 52.2 | healthy control | 19 | 17 |
| HDClarity | HC26 | female | 55.1 | healthy control | 22 | 19 |
| HDClarity | HC27 | male | 57 | healthy control | 17 | 15 |
| HDClarity | HC28 | male | 37.4 | healthy control | 24 | 19 |
| HDClarity | HC29 | female | 44.7 | healthy control | 17 | 17 |
| HDClarity | HC30 | male | 52.5 | healthy control | 22 | 15 |
| HDClarity | HC31 | female | 53.1 | healthy control | 17 | 17 |
| HDClarity | HC32 | male | 56.5 | healthy control | 18 | 17 |
| U of W | EPMHD14 | female | 31 | early pre-manifest HD | 40 | unknown |
| U of W | LPMHD20 | female | 34 | late pre-manifest HD | 50 | unknown |
| U of W | LPMHD21 | male | 46 | late pre-manifest HD | 42 | unknown |
| U of W | LPMHD22 | male | 55 | late pre-manifest HD | 42 | unknown |
| U of W | LPMHD23 | female | 53 | late pre-manifest HD | 46 | unknown |
| U of W | LPMHD24 | female | 38 | late pre-manifest HD | 48 | unknown |
| U of W | LPMHD25 | male | 31 | late pre-manifest HD | 44 | unknown |
| U of W | LPMHD26 | male | 35 | late pre-manifest HD | 35 | unknown |
| U of W | LPMHD27 | female | 52 | late pre-manifest HD | 42 | unknown |
| U of W | LPMHD28 | male | 42 | late pre-manifest HD | 42 | unknown |
| U of W | HC33 | female | 59 | Healthy control | unknown | unknown |
| U of W | HC34 | female | 21 | Healthy control | unknown | unknown |
| U of W | HC35 | female | 20 | Healthy control | unknown | unknown |
| U of W | HC36 | female | 51 | Healthy control | unknown | unknown |
| U of W | HC37 | female | 57 | Healthy control | unknown | unknown |
| U of W | HC38 | female | 60 | Healthy control | unknown | unknown |

U of W = University of Washington
EMHD = Early manifest HD
EPMHD = Early pre-manifest HD
LPMHD = Late pre-manifest HD
HC = Healthy control

#### Supplemental Table 2

| Sample ID | Source | Sample prepar-ation | Sex | PMI | Age at death | Sample Type | CAG low/ high | Onset | COD | Brain Region | Vonsattel Grade |
|---|---|---|---|---|---|---|---|---|---|---|---|
| HC113 | NZ Human Brain Bank | Fixed tissue | Male | 14 | 58 | HD | 28/4 4 | . | Broncho-pneumonia | Caudate Nucleus | 2 |
| HC114 | NZ Human Brain Bank | Fixed tissue | Fema le | 12 | 53 | HD | 21/4 7 | . | Pneumonia | Caudate Nucleus | 2 |
| HC120 | NZ Human Brain Bank | Fixed tissue | Male | 15 | 51 | HD | 10/4 6 | . | Pneumonia | Caudate Nucleus | 2 |
| HC126 | NZ Human Brain Bank | Fixed tissue | Male | 8 | 61 | HD | 17/4 3 | . | Pneumonia | Caudate Nucleus | 2 |
| HC109 | NZ Human Brain Bank | Fixed tissue | Fema le | 7 | 59 | HD | 23/4 7 | . | Bronchopne umonia | Caudate Nucleus | 4 |
| HC116 | NZ Human Brain Bank | Fixed tissue | Male | 8 | 54 | HD | 17/4 6 | . | Pneumonia | Caudate Nucleus | 4 |
| HC122 | NZ Human Brain Bank | Fixed tissue | Male | . | 52 | HD | 10/5 0 | . | Bowel obstruction | Caudate Nucleus | 4 |
| HC143 | NZ Human Brain Bank | Fixed tissue | Fema le | 16 | 45 | HD | . | . | Respiratory arrest | Caudate Nucleus | 4 |
| H170 | NZ Human Brain Bank | Fixed tissue | Male | 17 | 60 | Control | 10/1 7 | N/A | Ischemic heart disease | Caudate Nucleus | N/A |
| H189 | NZ Human Brain Bank | Fixed tissue | Male | 16 | 41 | Control | 18/2 2 | N/A | Asphyxia | Caudate Nucleus | N/A |
| H165 | NZ Human Brain Bank | Fixed tissue | Fema le | 26 | 43 | Control | 17/1 7 | N/A | Nitrogen poisoning | Caudate Nucleus | N/A |
| H238 | NZ Human Brain Bank | Fixed tissue | Fema le | 16 | 63 | Control | 14/1 6 | N/A | Dissecting aortic aneurysm | Caudate Nucleus | N/A |
| H145 | NZ Human Brain Bank | Fixed tissue | Male | 6.5 | 54 | Control | . | N/A | Ischemic heart disease | Caudate Nucleus | N/A |
| H140 | NZ Human Brain Bank | Fixed tissue | Male | 16 | 51 | Control | . | N/A | Cardiomyop athy | Caudate Nucleus | N/A |
| B3470 | Boston University | Fresh frozen tissue | Male | 57.30 | 89 | HD | 40 | 70 | . | Globus Pallidus | 3 |
| B3701 | Boston University | Fresh frozen tissue | Fema le | 14.30 | 67 | HD | 44 | 40 | . | Globus Pallidus | 3 |
| B3703 | Boston University | Fresh frozen tissue | Fema le | 25.00 | 61 | HD | 45 | 35 | . | Globus Pallidus | 3 |
| B4183 | Boston University | Fresh frozen tissue | Fema le | 20.30 | 76 | HD | 43 | . | . | Globus Pallidus | 4 |

| | | | | | | | | | | | |
|---|---|---|---|---|---|---|---|---|---|---|---|
| B4230 | Boston University | Fresh frozen tissue | Male | 7.08 | 76 | HD | 41 | 58 | . | Globus Pallidus | 3 |
| B4255 | Boston University | Fresh frozen tissue | Fema le | 17.50 | 52 | HD | 47 | . | . | Globus Pallidus | 4 |
| 5222 | Human brain and spinal fluid resource center | Fresh frozen tissue | Male | . | 61 | Control | N/A | N/A | Micro-infarct (cerebrum) | Globus Pallidus | N/A |
| 5214 | Human brain and spinal fluid resource center | Fresh frozen tissue | Male | . | 61 | Control | N/A | N/A | . | Globus Pallidus | N/A |
| 5293 | Human brain and spinal fluid resource center | Fresh frozen tissue | Fema le | . | 41 | Control | N/A | N/A | Liver failure | Globus Pallidus | N/A |
| 4308 | Human brain and spinal fluid resource center | Fresh frozen tissue | Male | . | 70 | Control | N/A | N/A | Myocardial infarction | Globus Pallidus | N/A |
| 5190 | Human brain and spinal fluid resource center | Fresh frozen tissue | Male | . | 68 | Control | N/A | N/A | Myocardial infarction | Globus Pallidus | N/A |
| AN05954 | Boston University | Fresh frozen tissue | Male | . | 49 | Pre-HD | . | N/A | . | Caudate and Putamen | N/A |
| AN16102 | Boston University | Fresh frozen tissue | Fema le | . | 86 | Pre-HD | . | N/A | . | Caudate and Putamen | N/A |
| AN18592 | Boston University | Fresh frozen tissue | Fema le | . | 82 | Control | N/A | N/A | . | Caudate and Putamen | N/A |
| AN15392 | Boston University | Fresh frozen tissue | Male | . | 51 | Control | N/A | N/A | . | Caudate and Putamen | N/A |

PMI = post mortem interval
COD = cause of death
HD = Huntington’s disease
N/A = not applicable
. = unknown

### Extended data figure 1

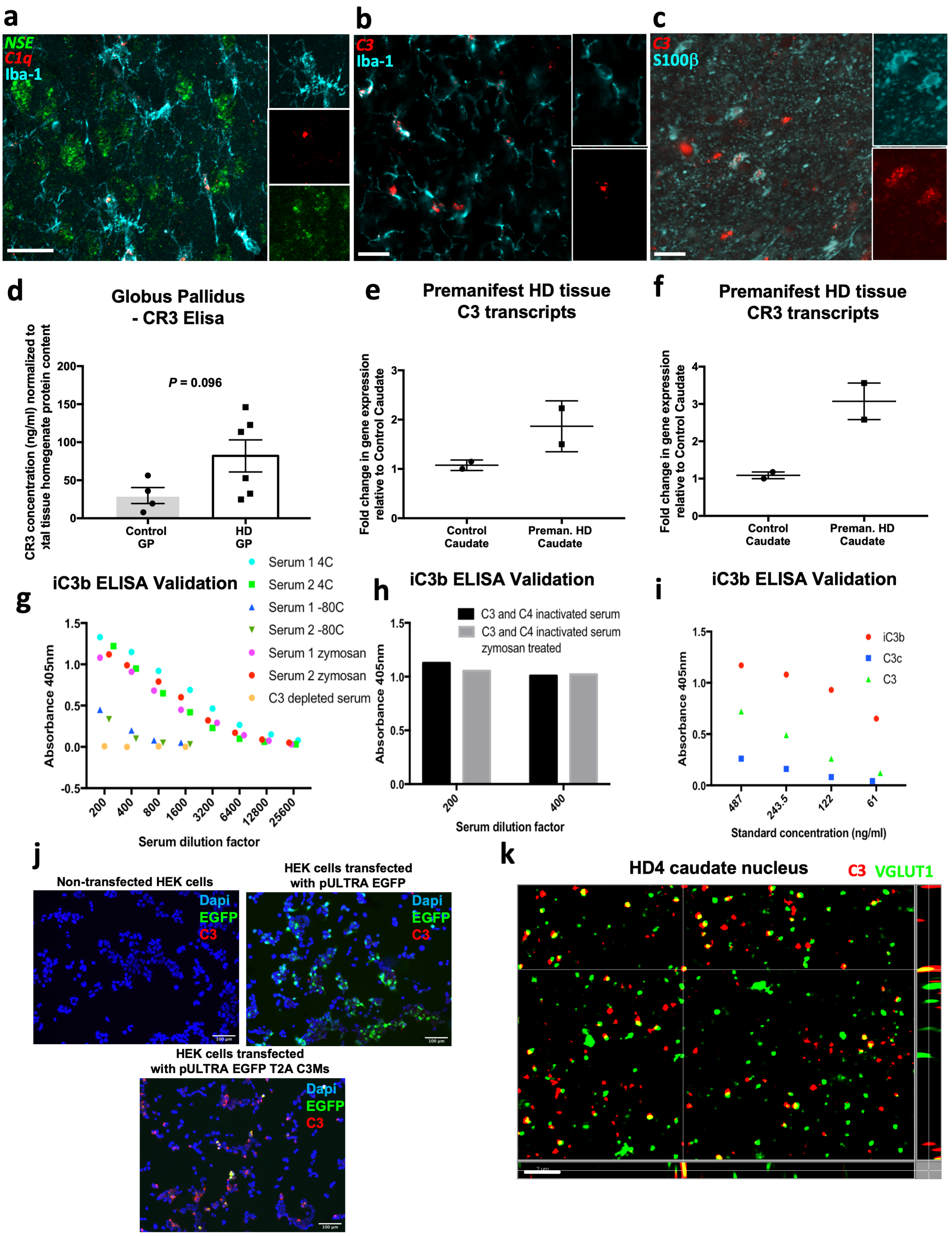

**Extended data figure 1: (a)** Representative confocal image of in situ staining for *C1Q* and *NSE* alongside IHC for microglial marker IBA1 in the caudate nucleus of postmortem tissue from an HD patient (Vonsattel grade 2). Scale bar = 20 μm. **(b)** Representative confocal image of in situ staining for *C3* alongside IHC for microglial marker IBA1 in the caudate nucleus of postmortem tissue from an HD patient (Vonsattel grade 2). Scale bar = 20 μm. **(c)** Representative confocal image of an in situ staining for *C3* alongside IHC for astrocytic marker S100β in the caudate nucleus of postmortem tissue from an HD patient (Vonsattel grade 2). Scale bar = 20 μm **(d)** Bar chart showing ELISA measurements of the concentration of complement receptor CR3 in proteins extracted from the globus pallidus (GP) of postmortem tissue from manifest HD patients and control (no documented evidence of neurodegenerative disease; see methods and supplemental table 2) individuals after normalization for total tissue homogenate protein content, n=4 control GP and 6 HD GP (there was not enough protein available from one of the control sample that had previously been employed in the C3, iC3b and hemoglobin ELISA's (the results of which are depicted in Figure 1d,e and f) and as such it was not tested here). Unpaired t-test p=0.096 **(e)** Dot plot showing the fold change in mRNA levels of complement component C3 in samples from the caudate nucleus of two premanifest HD patients relative to those seen in two clinically normal (see methods and Supplemental table 2) individuals. **(f)** Dot plot showing the fold change in mRNA levels of complement receptor CR3 in samples from the caudate nucleus of two premanifest HD patients relative to those seen in two clinically normal (see methods and Supplemental table 2) individuals. **(g)** Superimposed scatter blot showing the relative absorbance values of different serum samples employed in the iC3b ELISA. Note that in two independent serum samples, in which the complement pathway has been activated either by incubating the serum at 4 °C for 7 days or treating with 10 mg/ml of zymosan for 30 min at 37 °C, the absorbance values for the highest concentration of serum tested are approximately double that of the same samples left untreated and maintained at -80 °C. Thus demonstrating that this assay reflects complement cascade activation as would be predicted for an ELISA measuring levels of iC3b, a cleavage fragment of complement component C3 formed following cascade activation. **(h)** Bar graph showing the relative absorbance values of C3/C4 inactivated serum samples employed in the iC3b ELISA. Note that, unlike in **(g)** treatment of this serum with zymosan fails to increase levels of iC3b. Thus confirming the specificity of the ELISA by demonstrating that it reflects changes in a species that increases in response to complement cascade activation but is prevented from forming in the absence of functioning C3 and C4. **(i)** Superimposed scatter blot showing the relative absorbance values of different concentrations of complement component C3 standards purified from human serum using PEG precipitation and DEAE ion chromatography. Note that at all concentrations tested iC3b (generated by the cleavage of C3b with factor I in the presence of factor H) gave a higher absorbance value than uncleaved full length C3 or a subsequent cleavage component C3c (generated by treating iC3b with "trypsin like" proteases). Thus further demonstrating the specificity of this ELISA for iC3b versus the full-length protein or other cleavage components. **(j)** Representative images of non-transfected HEK 293 cells or those transfected with pULTRA EGFP or pULTRA EGFP T2A C3Ms stained with the same C3 antibody employed in the immunohistochemical analysis depicted in Figure 1 and in Figure 3, Extended data figure 3, and Extended data figure 7  Scale bar = 100 μm. **(k)** Orthogonal view of a representative structured illumination image showing C3 and VGLUT1 staining in the caudate nucleus of tissue from an HD patient (Vonsattel grade 4). Scale bar = 2 μm. For bar charts, bars depict the mean. All error bars represent SEM. Stars depict level of significance with *=p<0.05, **p=<0.01 and ***p<0.0001

### Extended data figure 2

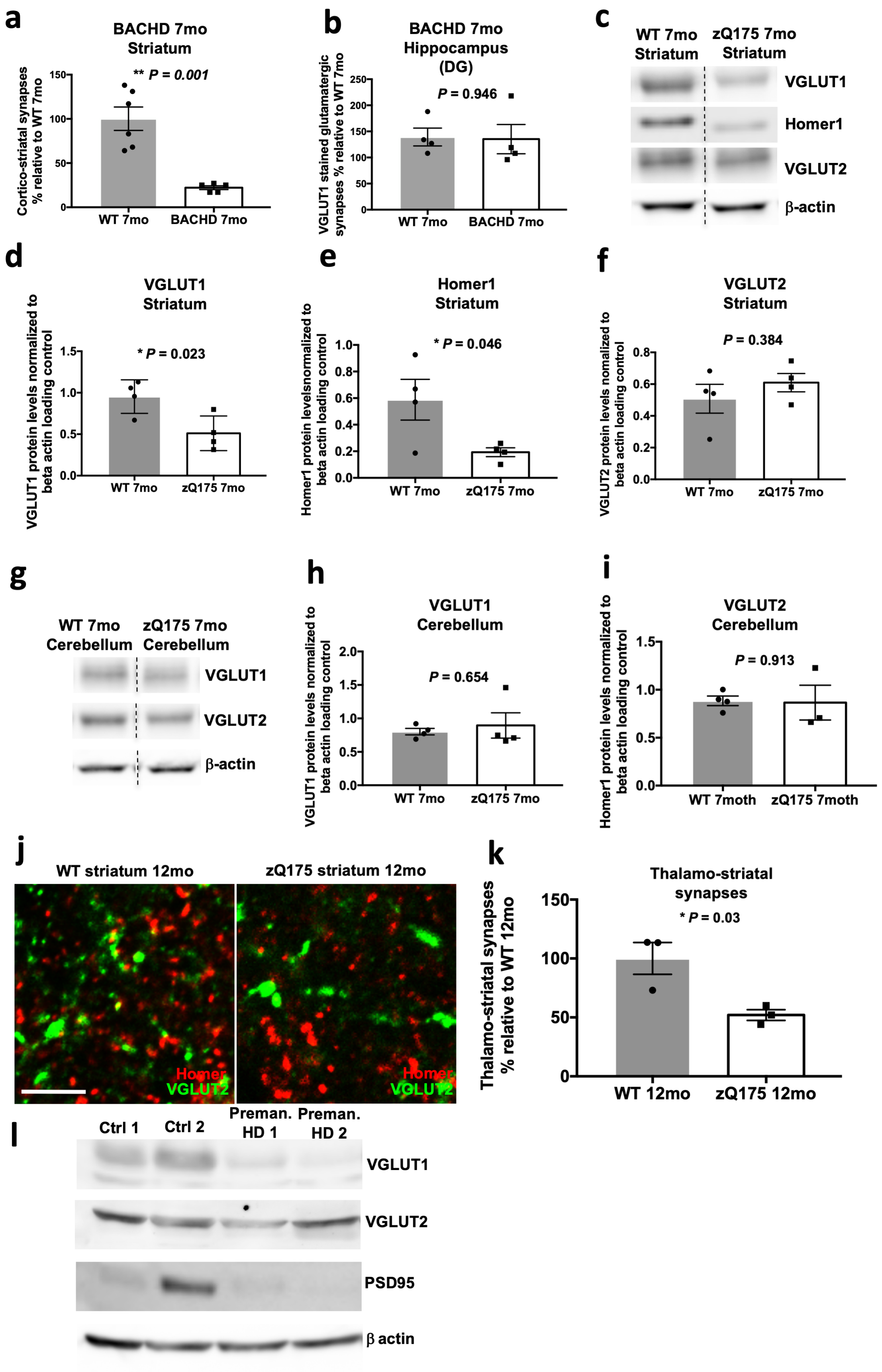

**Extended data figure 2: (a)** Bar chart shows quantification of corticostriatal synapses in the dorsolateral striatum of 7 mo BACHD mice and WT littermates. As seen in 7 mo zQ175 mice (Fig. 2a,b,c) there is a significant reduction in the number of corticostriatal synapses in the dorsolateral striatum of the BACHD mice, n=6 WT 5 BACHD mice. Unpaired t-test p=0.001. **(b)** Bar chart shows quantification of VGLUT1 stained glutamatergic synapses in the hippocampus (dentate gyrus) of 7 mo BACHD mice and WT littermates. Again, as seen in zQ175 mice at this age, (Fig. 2d) there is no difference in the number of glutamatergic synapses in this brain region in BACHD and WT mice, n=4 WT and 4 BACHD mice. Unpaired t-test p=0.946. **(c)** Representative immunoblot showing that levels of pre and postsynaptic markers of the corticostriatal synapse, VGLUT1 and Homer1 respectively, are reduced in the striatum of 7 mo zQ175 mice relative to that seen in WT littermate controls but levels of the thalamostriatal synapse marker, VGLUT2, are not. Hashed lines are used to denote the fact that non adjacent lanes from the same immunoblot are being depicted. Images showing the full lane chemiluminescent signal can be found in Source data figure 1. **(d,e,f)** Bar charts showing quantification of the relative protein levels of these synaptic markers in 7 mo zQ175 mice and WT littermates. Levels of VGLUT1 and Homer1 are significantly reduced in the zQ175 striatum but VGLUT2 levels are not changed, n=4 WT and 4 zQ175 mice. Unpaired t-tests, VGLUT1 p=0.0227; Homer1 p=0.0455; VGLUT2 p=0.3836 **(g)** Representative immunoblot showing that protein levels of VGLUT1 and VGLUT2 are not changed in the cerebellum of 7 mo zQ175 mice relative to that seen in WT littermate controls. Hashed lines are used to denote the fact that non adjacent lanes from the same immunoblot are being depicted. Images showing the full lane chemiluminescent signal can be found in Source data 1. **(h,i)** Bar charts showing quantification of relative protein levels in the cerebellum of 7 mo zQ175 mice and WT littermates, for VGLUT1 n=4 WT and 4 zQ175 mice for VGLUT2 n=4 WT and 3 zQ175 mice. Unpaired t-tests, VGLUT1 p=0.654; VGLUT2 p=0.913. **(j)** Representative confocal images of VGLUT2 and Homer1 staining in the dorsolateral striatum of 12 mo zQ175 mice and WT littermates. Scale bar = 5 $\mu$m. **(k)** Bar chart shows quantification of colocalized VGLUT2 and Homer1 puncta denoting thalamostriatal synapses in these mice. In older symptomatic zQ175 mice there is a significant loss of thalamostriatal synapses, n=3WT and 3zQ175 mice. Unpaired t-test, p=0.03 **(l)** Immunoblot of protein samples from the caudate nucleus of two premanifest HD patients and two clinically normal individuals stained with antibodies to VGLUT1, VGLUT2, PSD95 and $\beta$ actin. Note the reduction in VGLUT1 levels (a marker of the corticostriatal synapse) and decrease in PSD-95 levels (a marker of the postsynaptic density) but no change in VGLUT2 levels (a marker of the thalamostriatal synapse), n=2 caudate samples from control (clinically normal; see methods and supplemental table 2) individuals and 2 caudate samples from individuals with premanifest HD. Images showing the full lane chemiluminescent signal can be found in source data figure 2. For bar charts, bars depict the mean. All error bars represent SEM. Stars depict level of significance with *=p<0.05, **p=<0.01 and ***p<0.0001

### Extended data figure 3

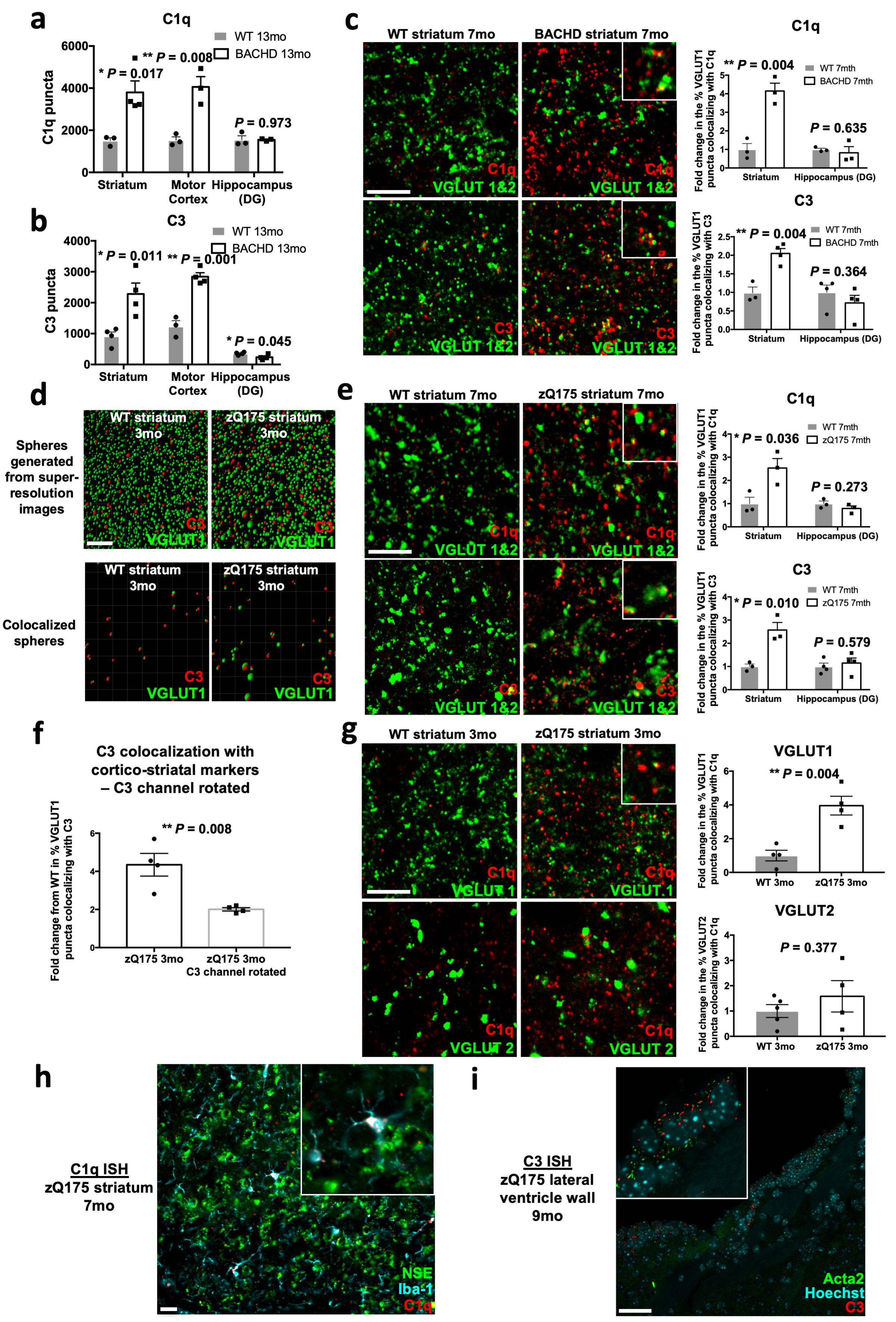

**Extended data figure 3: (a)** Bar charts showing quantification of C1q puncta in different brain regions of 13 mo BACHD mice and WT littermates. In disease affected regions (dorsolateral striatum and motor cortex) of BACHD mice but not the less affected dentate gyrus there is a significant increase in the levels of C1q relative to that seen in WT littermates, striatum n=3 WT and 4 BACHD mice, motor cortex n=3 WT and 3 BACHD mice and hippocampus (DG) n=3 WT and 3 BACHD mice. Unpaired t-test for comparisons of WT and BACHD in each brain region, striatum p=0.017, motor cortex p=0.008, hippocampus (DG) p=0.973. **(b)** Bar charts showing quantification of C3 puncta in different brain regions of 13 mo BACHD mice and WT littermates. In disease affected regions (dorsolateral striatum and motor cortex) of BACHD mice but not the less affected dentate gyrus there is a significant increase in the levels of C3 relative to that seen in WT littermates, striatum n=4 WT and 4 BACHD mice, motor cortex n=3 WT and 4 BACHD mice and hippocampus (DG) n=4 WT and 4 BACHD mice. Unpaired t-test for comparisons of WT and BACHD in each brain region, striatum p=0.011, motor cortex p=0.001, hippocampus (DG) p=0.045. **(c)** Representative confocal images of the dorsolateral striatum of 7 mo WT and BACHD mice co-stained with C1q and VGLUT1 and 2 or C3 and VGLUT1 and 2. Note the increased association of both complement proteins with these synaptic markers in the BACHD tissue. Insets show examples of complement proteins co-localized with the presynaptic markers VGLUT1 and 2. Scale bar = 5 μm. Bar charts show quantification of the percentage of VGLUT 1 and 2 puncta colocalizing with C1q or C3 in the disease affected striatum and less disease affected hippocampus (DG). There is a significant increase in the % of VGLUT 1 and 2 co-localizing with both complement proteins in the striatum but not in the dentate gyrus, for C1q n=3 WT and 3 BACHD mice for C3 n= 3 WT and 4 BACHD mice. Unpaired t-test for comparisons of WT and BACHD, Striatum C1q p=0.004, Hippocampus (DG) C1q p=0.635, Striatum C3 p=0.004, Hippocampus (DG) C3 p=0.364 **(d)** Representative pictographs of Imaris processed super-resolution images from the dorsolateral striatum of 7 mo WT and zQ175 mice co-stained with C3 and VGLUT1 in which the spheres function has been used to reflect immunoreactive puncta. The top two panels show all spheres generated from the super-resolution images and the bottom panels only show C3 and VGLUT1 spheres which are colocalized (defined as a distance of 0.3 μm or less between the center of each sphere). Scale bar = 5 μm. Note that there are more colocalized spheres in the 3 mo zQ175 striatum than in the 3 mo WT striatum. Quantification of these images is shown in the bar charts in Fig. 3g. **(e)** Confocal images of the dorsolateral striatum of 7 mo WT and zQ175 mice co-stained with C1q and VGLUT1 and 2 or C3 and VGLUT1 and 2. Bar charts show quantification of the percentage of VGLUT1 and 2 puncta colocalizing with C1q or C3 in disease affected regions (striatum) or less affected regions (dentate gyrus of the hippocampus). As seen in the BACHD model there is a significant increase in the % of VGLUT1 and 2 puncta co-localizing with both complement proteins in the striatum but not in the dentate gyrus of zQ175 mice relative to that seen in their WT littermates, for C1q n=3 WT and 3 zQ175 mice, for C3 striatum n=3 WT and 3 zQ175 mice and for C3 hippocampus (DG) n=4 WT and 4 zQ175 mice. Unpaired t-test for comparisons of WT and zQ175, Striatum C1q p=0.036, Hippocampus (DG) C1q p=0.273, Striatum C3 p=0.010, Hippocampus (DG) C3 p=0.579. **(f)** Bar chart comparing the fold enrichment of C3 at VGLUT1 puncta in 3 mo zQ175 mice (taken from Fig. 3g) with that same analysis carried out after rotating the C3 channel 90 degrees, n=4 WT and 4 zQ175 mice. Unpaired t-test p=0.008 **(g)** Representative confocal images of the dorsolateral striatum of 3 mo zQ175 and WT mice co-stained with antibodies to C1q and VGLUT1 or C1q and VGLUT2. Scale bar = 5 μm. Bar charts show quantification of the % of VGLUT1 or VGLUT2 puncta colocalizing with C1q in both genotypes. There is a significant increase in the % of VGLUT1 puncta co-localized with C1q but not VGLUT2 puncta co-localized with C1q in the zQ175 mice relative to that seen in their WT littermates, for VGLUT1 n=4 WT and 4 zQ175 mice, for VGLUT2 n=5 WT and 4 zQ175 mice. Unpaired t-test for comparisons of WT and zQ175, VGLUT1 p=0.004, VGLUT2 p=0.377 **(h)** Representative in situ hybridization (ISH) staining of *C1q* and *NSE* together with IHC for Iba1 in the dorsal striatum of 7 mo zQ175 mice. Inset shows a magnification of a selected area in the field. Scale bar = 20 μm **(i)** Representative ISH staining of *C3* and *Acta2* in the wall of the lateral ventricle. Inset shows a magnification of a selected area in the field Scale bar = 50 μm. For bar charts, bars depict the mean. All error bars represent SEM. Stars depict level of significance with *=p<0.05, **p=<0.01 and ***p<0.0001

### Extended data figure 4

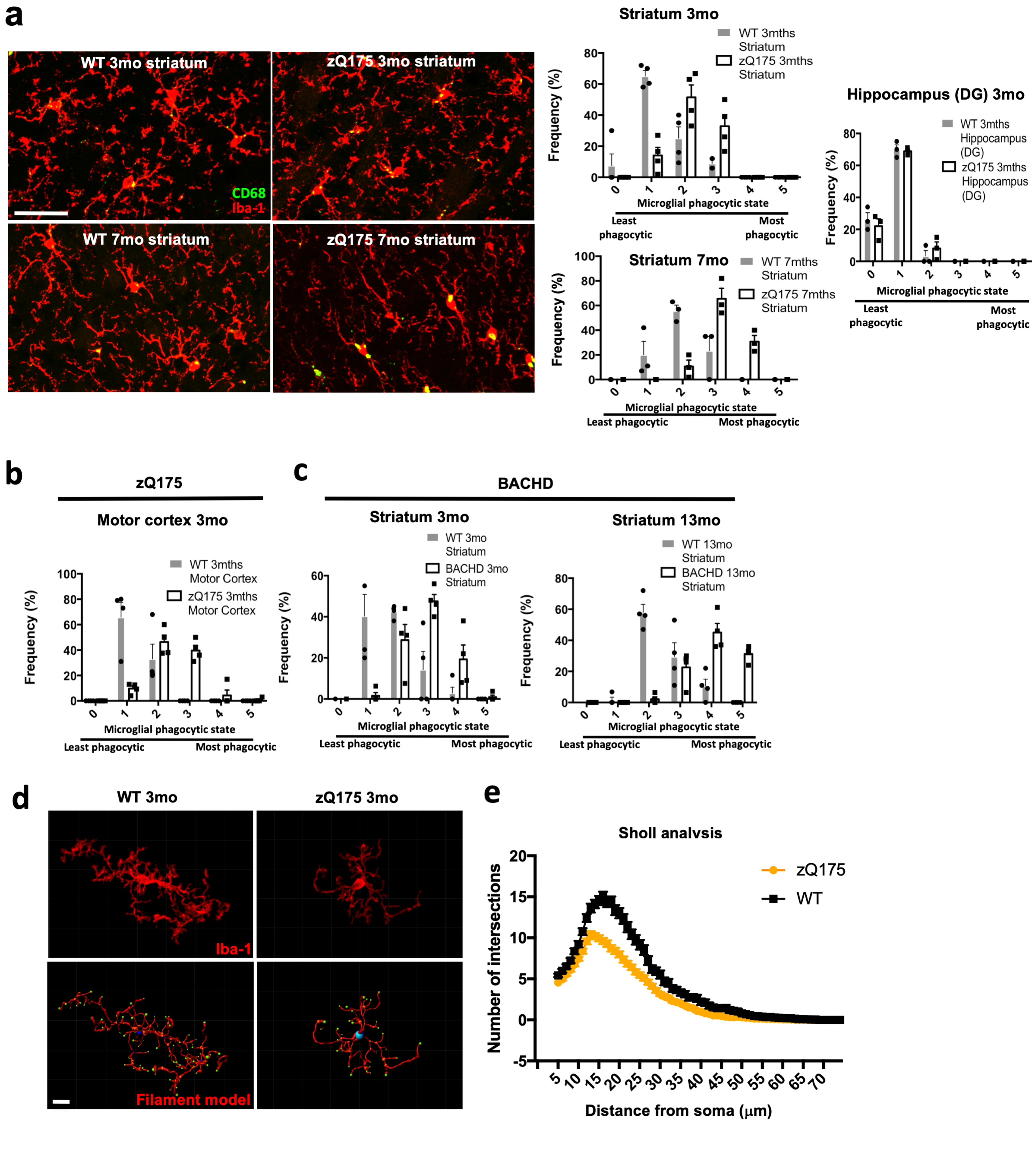

**Extended data figure 4: (a)** Representative maximum intensity projections generated from confocal images of Iba1 and CD68 staining in the dorsal striatum of 3 and 7 mo zQ175 mice and their WT littermates. Scale bar = 20 μm. Note the increased level of the lysosomal marker CD68 and the reduced branching and thicker processes of the microglia in the zQ175 mice at both ages. Bar charts show quantification of the phagocytic state of microglia in these images with 5 being the most phagocytic and 0 the least. At both 3 and 7 mo microglia in the dorsal striatum of zQ175 mice show a shift towards a more phagocytic state, for 3 mo n=4 WT and 4 zQ175 mice; for 7 mo n=3 WT and 3 zQ175mice. Two way anova, at 3 mo interaction of score and genotype p=<0.0001; at 7 mo interaction of score and genotype p=<0.0001. This is not the case in less disease affected regions such as the dentate gyrus of the hippocampus, n=3 WT and 3 zQ175 mice. Two-way anova, interaction of score and genotype p=0.641. **(b)** Bar chart showing quantification of microglial phagocytic state in the motor cortex of 3 mo zQ175 mice and WT littermates. There is a shift towards a more phagocytic state in the zQ175 mice, n=4 WT and 4 zQ175 mice. Two way anova, at 3 mo interaction of score and genotype p=<0.0001. **(c)** Bar charts, showing that there is also a shift towards a more phagocytic microglial state in the striatum of 3 mo and 13 mo BACHD mice, for both 3 mo and 13 mo n=4 WT and 4 BACHD mice. Two way anova, at 3 mo interaction of score and genotype p=<0.0001; at 13 mo interaction of score and genotype p=<0.0001 **(d)** Confocal images and filament renderings of individual microglia stained with Iba1 in the dorsal striatum of 3 mo zQ175 mice and WT littermates. Scale bar = 10 μm. In the filament renderings the blue sphere indicates the soma, orange spheres denote branch points and green spheres indicate terminal points of microglial processes. **(e)** Sholl analysis of confocal images of microglia from the dorsal striatum of 3 mo zQ175 mice and WT littermates. Analysis was performed on filament rendered images using Imaris software, n=3 WT and 4 zQ175 mice with over 100 cells analyzed per genotype. Two way anova, p=<0.0001 with Sidak's multiple comparisons test shows a significant difference between WT and zQ175 at distances from the soma ranging from 11 to 30 μm p<0.0001. For bar charts, bars depict the mean. All error bars represent SEM. Stars depict level of significance with *=p<0.05, **p=<0.01 and ***p<0.0001.

### Extended data figure 5

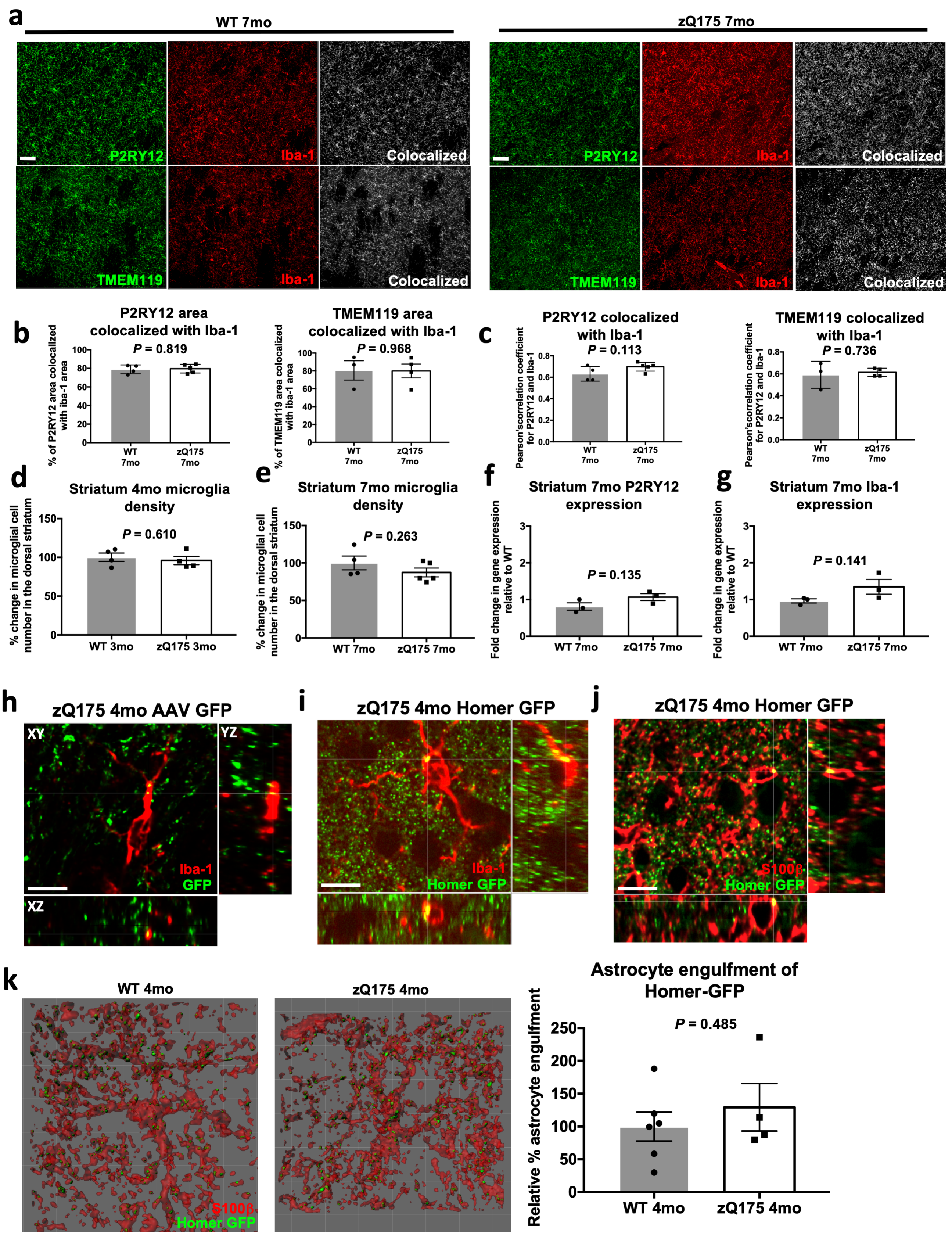

**Extended data figure 5: (a)** Confocal images of microglia in the dorsal striatum of 7 mo zQ175 mice and WT littermates co-stained with antibodies to Iba1 and putative microglia identity markers P2RY12 and TMEM119. Scale bar = 50 μm **(b)** Bar charts show the % of P2RY12 and TMEM119 immunoreactive area above a set threshold (determined using an algorithm developed by Costes and Lockett[199]) that colocalizes with the area of Iba1 staining for the images in **(a)**, for P2RY12 and Iba1 n=4 WT and 5 zQ175 mice; for TMEM119 and Iba1 n=3 WT and 4 zQ175 mice. Unpaired t-test for comparison of WT and zQ175 mice, for P2RY12 and Iba1 p=0.819 and for TMEM119 and Iba1 p=0.968. **(c)** Bar charts show the Pearson's correlation coefficient for Iba1 and P2RY12 and Iba1 and TMEM119 for the images shown in **(a)**, for P2RY12 and Iba1 n=4 WT and 5 zQ175 mice; for TMEM119 and Iba1 n=3 WT and 4 zQ175 mice. Unpaired t test for comparison of WT and zQ175 mice P2RY12 and Iba1 p=0.113 and for TMEM119 and Iba1 p=0.736. **(d)** Bar chart shows quantification of microglial cell density in the dorsal striatum of 4 mo zQ175 mice and WT littermates, n=4 WT and 4 zQ175 mice. Unpaired t-test p=0.610. **(e)** Bar chart shows quantification of microglial cell density in the striatum of 7 mo Q175mice and WT littermates, n=4 WT and 5 zQ175 mice. Unpaired t-test p=0.263 **(f)** Bar chart shows the level of P2RY12 transcripts in striatal extracts from 7 mo zQ175 mice and WT littermates, n=3 WT and 3 zQ175 mice. Unpaired t-test p=0.135. **(g)** Bar chart shows the level of Iba1 transcripts in striatal extracts from 7 mo zQ175 and mice and WT littermates, n=3 WT and 3 zQ175 mice. Unpaired t-test p=0.141. **(h)** Representative orthogonal view of an Iba1 stained microglia in the dorsal striatum of a 4 mo zQ175 mice that had previously received a motor cortex injection of pAAV2-hsyn-EGFP at P1/2. Scale bar = 10 μm **(i)** Representative orthogonal view of an Iba1 stained microglia in the dorsal striatum of a 4 mo zQ175 Homer-GFP mouse. Scale bar = 10 μm **(j)** Representative orthogonal view of an S100β stained astrocyte in the dorsal striatum of a 4 mo zQ175 Homer-GFP mouse. Scale bar = 10 μm **(k)** Representative surface rendered images of S100β stained astrocytes (red) and engulfed Homer-GFP inputs (green) in the dorsal striatum of 4 mo zQ175 Homer-GFP mice and WT Homer-GFP littermates. Scale bar = 10 μm. Bar chart shows quantification of the relative % astrocyte engulfment of Homer-GFP (the volume of engulfed Homer-GFP expressed as a percentage of the total volume of the astrocyte) in 4 mo zQ175 Homer-GFP mice relative to that seen in WT Homer-GFP littermate controls, n=6 WT Homer-GFP and 4 zQ175 Homer-GFP mice. Unpaired t-test, p=0.485. For bar charts, bars depict the mean. All error bars represent SEM. Stars depict level of significance with *=p<0.05, **p=<0.01 and ***p<0.0001.

### Extended data figure 6

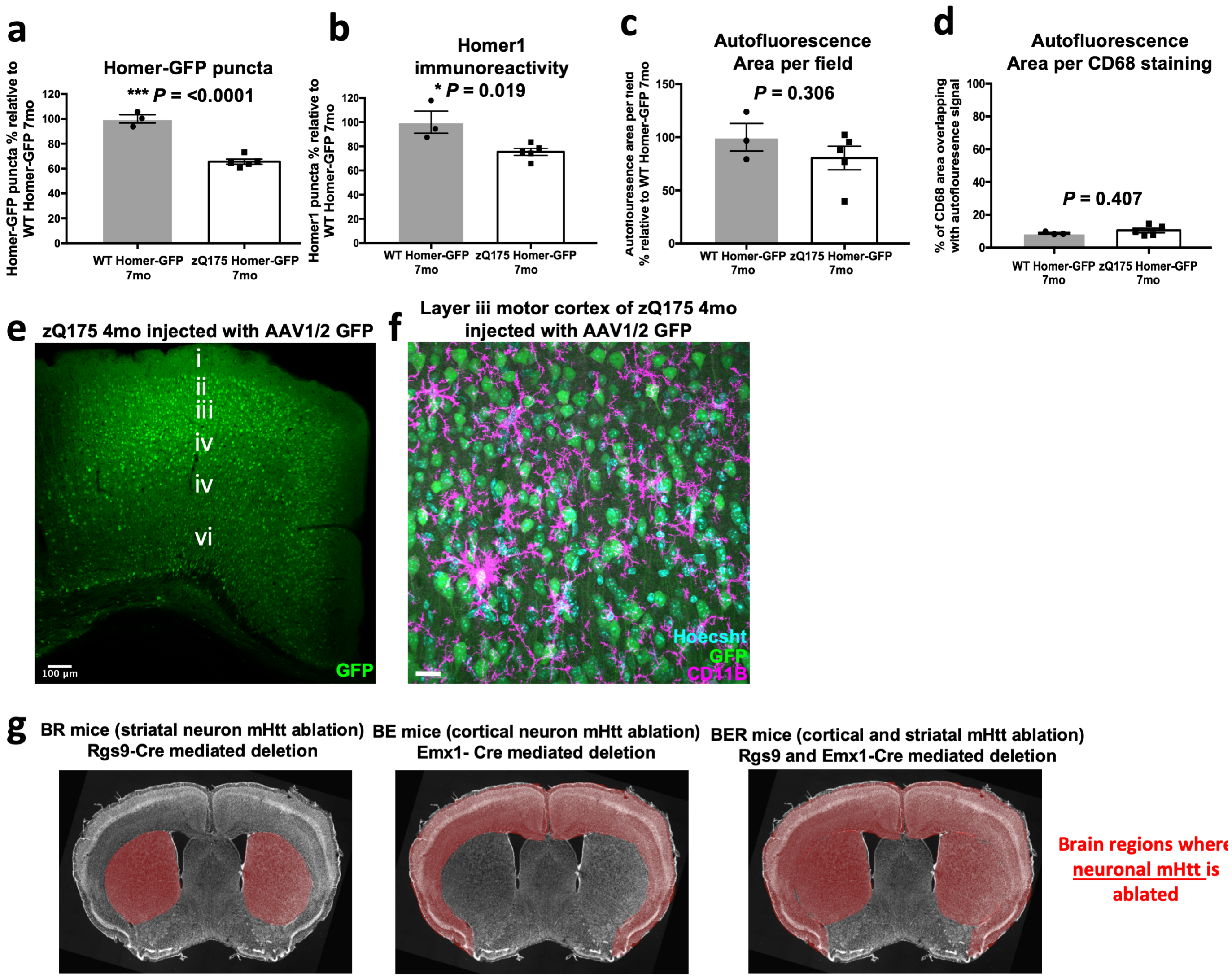


**Extended data figure 6: (a,b)** Bar charts showing the percentage of Homer-GFP puncta **(a)** and the percentage of Homer1 immunoreactive puncta **(b)** in 7 mo zQ175 Homer-GFP mice relative to that seen in 7 mo WT Homer-GFP littermates, n=3 WT Homer-GFP and 5 zQ175 Homer-GFP mice. Unpaired t-test, for Homer-GFP puncta p=<0.0001; for Homer1 immunoreactive puncta p=0.019. **(c)** Bar chart showing the relative % area per field occupied by auto-fluorescent signal in 7 mo zQ175 Homer-GFP mice relative to that seen in 7 mo WT Homer-GFP littermates, n=3 WT Homer-GFP and 5 zQ175 Homer-GFP mice. Unpaired t-test p=0.306. **(d)** Bar chart showing the % of CD68 area overlapping with autofluorescent signal in 7 mo zQ175 Homer-GFP mice and WT Homer-GFP littermates, n=3WT Homer-GFP and 5 zQ175 Homer-GFP mice. Unpaired t-test p=0.407. **(e)** Representative 10x confocal image of the motor cortex of a 4 mo zQ175 mouse injected with pAAV2-hsyn-EGFP showing the distribution of transduced cells in different cortical layers. Scale bar =100 µm. **(f)** Representative 60x confocal image of layer iii of the motor cortex of a 4 mo zQ175 mice injected with pAAV2-hsyn-EGFP showing transduced cells and microglia denoted by CD11B staining. Scale bar = 10 µm. **(e)** Coronal section of a mouse brain showing the brain regions (highlighted in red) in which *mHtt* has been genetically removed from selected neuronal populations in BR (RGS-9 cre to excise *mHtt* in striatal neurons) BE (Emx-1 cre to excise *mHtt* in cortical neurons) and BER (Rgs-9 and Emx-1 cre to excise *mHtt* in both striatal and cortical neurons) mice. For bar charts, bars depict the mean. All error bars represent SEM. Stars depict level of significance with *=p<0.05, **p=<0.01 and ***p<0.0001.

### Extended data figure 7

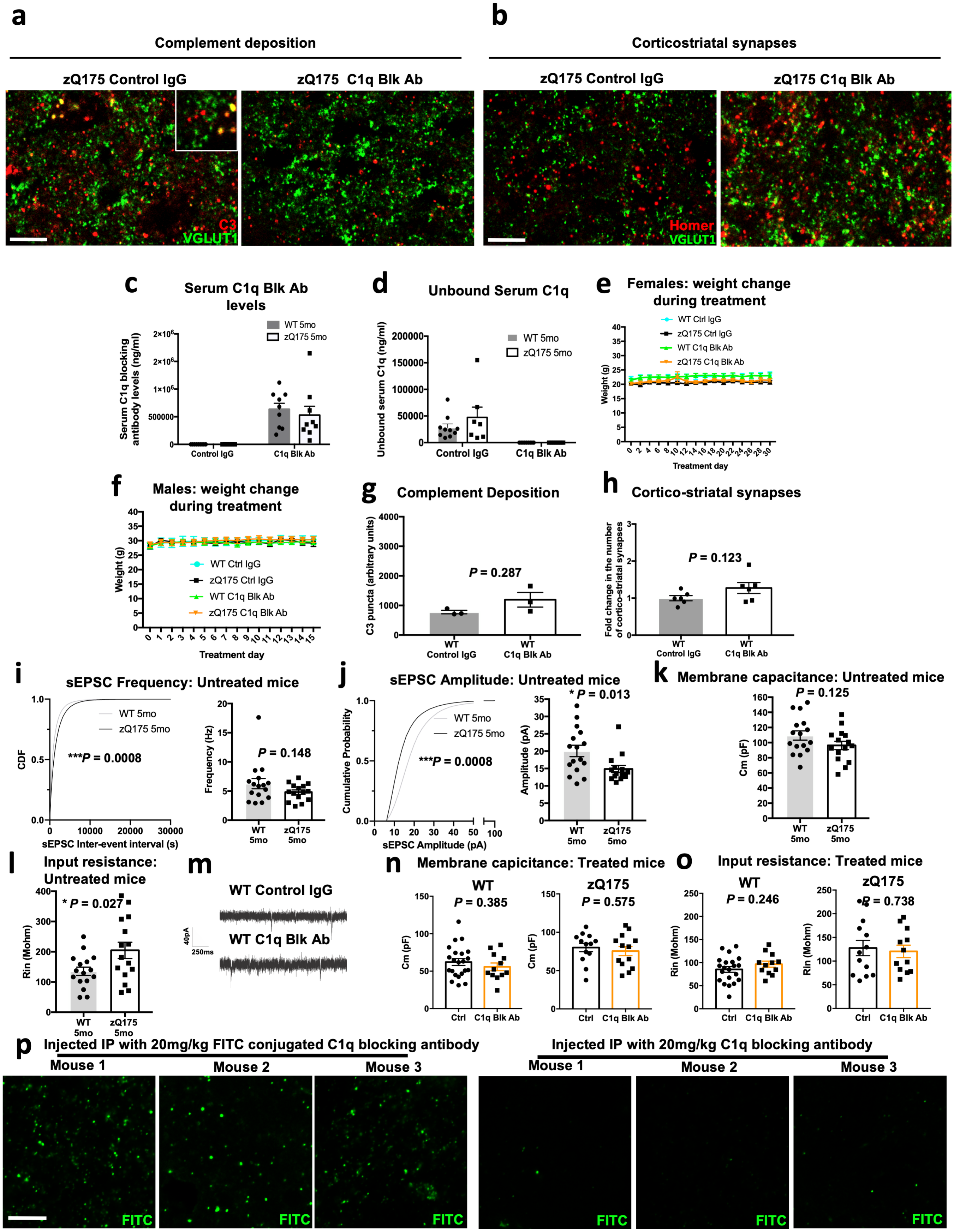

**Extended data figure 7: (a)** Representative confocal images of VGLUT1 and C3 staining in the dorsolateral striatum of 4 mo zQ175 mice that received intraperitoneal injections of a C1q function blocking antibody (M1) or a control IgG. Scale bar = 5 μm. **(b)** Representative confocal images of Homer1 and VGLUT1 staining in the dorsolateral striatum of zQ175 mice treated with the C1q function blocking antibody or a control IgG. Scale bar = 5 μm. **(c)** Bar chart showing quantification of the serum levels of the C1q function blocking antibody in 5 mo zQ175 mice and WT littermates following 1 mo treatment with either the blocking antibody or a control IgG. As expected mice treated with the blocking antibody have significantly higher levels of the antibody in their serum and the levels are not significantly different between treated WT and zQ175 mice, n=10 WT mice with control IgG; 9 WT mice with C1q Blk Ab; 7 zQ175 mice with control IgG; 9 zQ175 mice with C1q Blk Ab. Two way anova p= <0.0001 for treatment type and p=0.6 for genotype with WT control IgG vs zQ175 control IgG p=>0.999; WT control IgG vs WT C1q Blk Ab p=0.0003; WT control IgG vs zQ175 C1q Blk Ab p=0.003; zQ175 control IgG vs WT C1q Blk Ab p=0.001; zQ175 control IgG vs zQ175 C1q Blk Ab p=0.007; and WT C1q Blk Ab vs zQ175 C1q Blk Ab p=0.973 via Sidak's multiple comparisons test. **(d)** Bar chart showing levels of unbound C1q in the same serum tested in **(c).** There is significantly less unbound C1q in the serum of mice treated with the C1q function blocking antibody than in those treated with the control IgG but again no difference was seen in the levels present in blocking antibody treated WT and zQ175 mice, n=10 WT mice with control IgG; 9 WT mice with C1q Blk Ab; 7 zQ175 mice with control IgG; 9 zQ175 mice with C1qBlkAb. Two way anova p=0.0001 for treatment type and p=0.286 for genotype with WT control IgG vs zQ175 control IgG p=0.603; WT control IgG vs WT C1q Blk Ab p=0.123; WT control IgG vs zQ175 C1q Blk Ab p=0.123; zQ175 control IgG vs WT C1q Blk Ab p=0.005; zQ175 control IgG vs zQ175 C1q Blk Ab p=0.005; WT C1q Blk Ab vs zQ175 C1q Blk Ab p=>0.999 via Sidak's multiple comparisons test. **(e)** Weight changes in female mice treated with the C1q function blocking antibody or a control IgG, n=7 WT with Ctrl IgG, n=5 zQ175 with Ctrl IgG, n=7 WT with C1q Blk Ab, n=4 zQ175 with C1q Blk Ab. Two way anova with genotype/treatment as a source of variation p=0.223; WT Ctrl IgG vs zQ175 Ctrl IgG p=0.324; WT Ctrl IgG vs WT C1q Blk Ab p=0.999; zQ175 Ctrl IgG vs zQ175 C1q Blk Ab p=0.970; WT C1q Blk Ab vs zQ175 C1q Blk Ab p=0.619; WT Ctrl IgG vs zQ175 C1q Blk Ab p = 0.659; zQ175 Ctrl IgG vs WT C1q Blk Ab p = 0.294 via Sidak's multiple comparisons test. **(f)** Weight changes in male mice treated with the C1q function blocking antibody or a control IgG, n=4 WT Ctrl IgG, n=5 zQ175 Ctrl IgG, n=5 WT C1q Blk Ab, n= 7zQ175 C1q Blk Ab. Two way anova with genotype/treatment as a source of variation p=0.959, with WT Ctrl IgG vs zQ175 Ctrl IgG p=0.995; WT Ctrl IgG vs WT C1q Blk Ab p=0.989; WT Ctrl IgG vs zQ175 C1q Blk Ab p= 0.999; zQ175 Ctrl IgG vs zQ175 C1q Blk Ab p=0.979; WT C1q Blk Ab vs zQ175 C1q Blk Ab p=0.961; zQ175 Ctrl IgG vs WT C1q Blk Ab p=0.999 via Sidak's multiple comparisons test **(g)** Bar chart showing quantification of C3 puncta in the neuropil of 5 mo WT mice treated for 1 mo with the C1q function blocking antibody or a control IgG, n=3 mice. Unpaired t-test p=0.177. **(h)** Bar chart showing quantification of corticostriatal synapses in the dorsolateral striatum of 5 mo WT mice treated for 1 mo with the C1q function blocking antibody or a control IgG, n=3 mice. Unpaired t-test p=0.287. **(i)** Cumulative distribution plot of interspike intervals (ISI) obtained from whole-cell voltage clamp recordings of medium spiny neuron spontaneous excitatory postsynaptic currents. Recordings were carried out in slices from 5 mo WT and zQ175 mice (Grey = WT, Black = zQ175). Bar chart shows average frequency (Hz) per cell recorded across conditions, n=16 WT cells and 15 zQ175 cells from 5 WT and 4 zQ175 mice. Kolmogorov-Smirnov test for the cumulative distribution plot p=0.0008; Unpaired t test for the bar chart, p=0.148. **(j)** Cumulative distribution plot of amplitude obtained from whole-cell voltage clamp recordings of medium spiny neuron spontaneous excitatory postsynaptic currents. Recordings were carried out in slices from 5 mo WT and zQ175 mice (Grey = WT, Black = zQ175). Bar chart shows average amplitude (pA) per cell recorded across conditions, n=16 WT cells and 15 zQ175 cells from 5 WT and 4 zQ175 mice. Kolmogorov-Smirnov test for the cumulative distribution plot p=0.0008; Unpaired t test for the bar chart, p=0.013. **(k)** Bar chart showing the mean capacitance per cell from sEPSC recordings of MSN's in slices taken from 5 mo zQ175 mice and WT littermates, n=16 WT cells and 15 zQ175 cells from 5 WT and 4 zQ175 mice . Unpaired t-test, p=0.125. **(l)** Bar chart showing the mean input resistance per cell from sEPSC recordings of MSN's in slices taken from 5 mo zQ175 mice and WT littermates, n=16 WT cells and 15 zQ175 cells from 5 WT and 4 zQ175 mice. Unpaired t-test, p=0.027. **(m)** Representative traces of sEPCS recorded from MSN's in striatal slices from 5 mo WT mice which had been treated for 1 mo with control IgG or the C1q function blocking antibody. **(n)** Bar charts showing the mean capacitance per cell from sEPSC recordings of MSN's in slices taken from 5 mo zQ175 mice and WT littermates treated with the C1q Blk antibody (M1) or a control IgG, n=13-23 cells per condition from 4-7 animals. Unpaired t-test for WT p=0.385; for zQ175 p=0.575. **(o)** Bar charts showing the mean input resistance per cell from sEPSC recordings of MSN's in slices taken from 5 mo zQ175 mice and WT littermates treated with the C1q Blk antibody (M1) or a control IgG, n=13-23 cells per condition from 4-7 animals. Unpaired t-test for WT p=0.246;

for zQ175 $p=0.738$. **(p)** Representative confocal images of the dorsal striatum of 7 mo zQ175 mice injected IP with 20mg/kg of FITC conjugated C1q function blocking antibody, or unconjugated blocking antibody 24 h prior to sacrifice. Scale bar = 10 μm. Note that only in mice treated with FITC conjugated C1q function blocking antibody is there evidence of punctate staining in the neuropil, n=3 7 mo zQ175 mice treated with 20mg/kg of FITC conjugated C1q function blocking antibody and n=3 7 mo zQ175 mice treated with 20mg/kg of unconjugated C1q function blocking antibody. For bar charts, bars depict the mean and all error bars represent SEM. Stars depict level of significance with *=$p<0.05$, **p=<0.01 and ***$p<0.0001$.

### Extended data figure 8

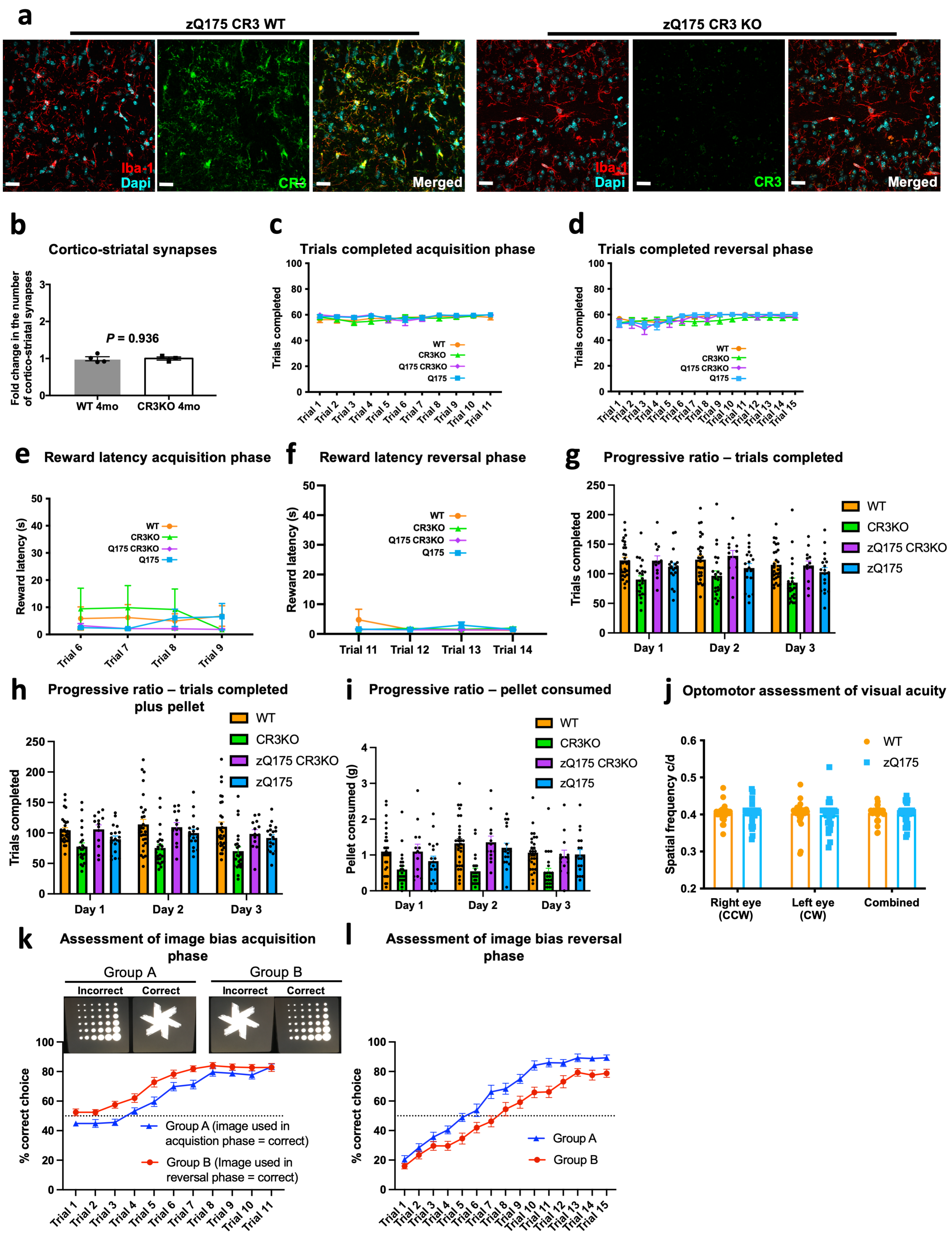

**Extended data figure 8: (a)** Representative confocal images of the dorsal striatum of 4 mo zQ175 CR3 WT and 4 mo zQ175 CR3 KO mice stained with antibodies to Iba-1 and CR3. Scale bar = 20 μm. **(b)** Bar chart showing quantification of corticostriatal synapses in the dorsolateral striatum of 4 mo CR3KO mice and WT littermates, n=4 WT mice and 3 CR3KO mice. Unpaired t-test p=0.9356. **(c)** Line graph showing the mean number of trials completed on each trial day for each genotype during the acquisition phase (maximum = 60), n= 29 WT mice, 18 zQ175 mice, 24 CR3KO mice and 13 zQ175 CR3KO mice. Two way anova: for WT vs zQ175 p=0.535 for the combination of genotype x trial session as a significant source of variation and p=0.461 for genotype as a significant source of variation; for zQ175 vs zQ175 CR3KO p=0.501 for the combination of genotype x trial session as a significant source of variation and p=0.007 for genotype as a significant source of variation; for WT vs zQ175 CR3KO p=0.689 for the combination of genotype x trial session as a significant source of variation and p=0.569 for genotype as a significant source of variation; for WT vs CR3 KO p=0.908 for the combination of genotype x trial session as a significant source of variation and p=0.888 for genotype as a significant source of variation. **(d)** Line graph showing the mean number of trials completed on each trial day for each genotype during the reversal phase (maximum = 60), n= 29 WT mice, 18 zQ175 mice, 24 CR3KO mice and 13 zQ175 CR3KO mice. Two way anova: for WT vs zQ175 p=0.787 for the combination of genotype x trial session as a significant source of variation and p=0.708 for genotype as a significant source of variation; for zQ175 vs zQ175 CR3KO p=0.899 for the combination of genotype x trial session as a significant source of variation and p=0.448 for genotype as a significant source of variation; for WT vs zQ175 CR3KO p=0.960 for the combination of genotype x trial as a significant source of variation and p=0.331 for genotype as a significant source of variation; for WT vs CR3 KO p=0.132 for the combination of genotype x trial session as a significant source of variation and p=0.361 for genotype as a significant source of variation. **(e)** Line graph showing reward latency for each genotype for trial days 6,7,8 and 9 in the acquisition phase, n= 29 WT mice, 18 zQ175 mice, 24 CR3KO mice and 13 zQ175 CR3KO mice. Two way anova: for WT vs zQ175 p=0.499 for the combination of genotype x trial session as a significant source of variation and p=0.750 for genotype as a significant source of variation; for zQ175 vs zQ175 CR3KO p=0.526 for the combination of genotype x trial session as a significant source of variation and p=0.362 for genotype as a significant source of variation; for WT vs zQ175 CR3KO p=0.868 for the combination of genotype x trial as a significant source of variation and p=0.525 for genotype as a significant source of variation; for WT vs CR3 KO p=0.240 for the combination of genotype x trial session as a significant source of variation and p=0.812 for genotype as a significant source of variation. **(f)** Line graph showing reward latency for each genotype for trial days 11,12,13 and 14 in the reversal phase, n= 29 WT mice, 18 zQ175 mice, 24 CR3KO mice and 13 zQ175 CR3KO mice. Two way anova: for WT vs zQ175 p=0.487 for the combination of genotype x trial session as a significant source of variation and p=0.804 for genotype as a significant source of variation; for zQ175 vs zQ175 CR3KO p=0.271 for the combination of genotype x trial session as a significant source of variation and p=0.148 for genotype as a significant source of variation; for WT vs zQ175 CR3KO p=0.751 for the combination of genotype x trial as a significant source of variation and p=0.590 for genotype as a significant source of variation; for WT vs CR3 KO p=0.431 for the combination of genotype x trial session as a significant source of variation and p=0.634 for genotype as a significant source of variation. **(g)** Bar chart showing the average total number of trials completed by each genotype in the progressive ratio task, n= 29 WT mice, 18 zQ175 mice, 24 CR3KO mice and 13 zQ175 CR3KO mice. Two way anova: for WT vs zQ175 p=0.919 for the combination of genotype x testing day as a significant source of variation and p=0.162 for genotype as a significant source of variation; for zQ175 vs zQ175 CR3KO p=0.391 for the combination of genotype x testing day as a significant source of variation and p=0.175 for genotype as a significant source of variation; for WT vs zQ175 CR3KO p=0.583 for the combination of genotype x testing day as a significant source of variation and p=0.839 for genotype as a significant source of variation; for WT vs CR3 KO p=0.754 for the combination of genotype x testing day as a significant source of variation and p=0.0012 for genotype as a significant source of variation. **(h)** Bar chart showing the average total number of trials completed by each genotype in the progressive ratio task when the assay is conducted in the presence of a food pellet, n= 29 WT mice, 18 zQ175 mice, 24 CR3KO mice and 13 zQ175 CR3KO mice. Two way anova: for WT vs zQ175 p=0.813 for the combination of genotype x testing day as a significant source of variation and p=0.085 for genotype as a significant source of variation; for zQ175 vs zQ175 CR3KO p=0.592 for the combination of genotype x testing day as a significant source of variation and p=0.175 for genotype as a significant source of variation; for WT vs zQ175 CR3KO p=0.247 for the combination of genotype x testing day as a significant source of variation and p=0.645 for genotype as a significant source of variation; for WT vs CR3 KO p=0.051 for the combination of genotype x testing day as a significant source of variation and p=0.0003 for genotype as a significant source of variation. **(i)** Bar chart showing the amount in grams of a food pellet consumed by each genotype while carrying out the progressive ratio task, n= 29 WT mice, 18 zQ175 mice, 24 CR3KO mice and 13 zQ175 CR3KO mice. Two way anova: for WT vs zQ175 p=0.470 for the combination of genotype x testing day as a significant source of variation and p=0.321 for genotype as a significant source of variation; for zQ175 vs zQ175 CR3KO p=0.399 for the combination of genotype x testing day as a significant source of variation and p=0.482 for genotype as a significant source of variation; for WT vs zQ175 CR3KO p=0.826 for the combination of genotype x testing day as a significant source of variation and p=0.898 for genotype as a significant source of variation; for WT vs CR3 KO p=0.153 for the combination of genotype x testing day as a significant source of variation and p=<0.0001 for genotype as a significant source of variation. **(j)** Bar chart showing the average performance of WT and zQ175 mice in the optomotor assay of visual acuity. Note the equivalent performance of both genotypes in this test, n = 33 WT mice and 28 zQ175 mice. Two anova for WT vs zQ175 p=0.598 for genotype as a significant source of variation with p=0.982, 0.933 and 0.962 for the right eye, left eye or the combined performance of both respectively via Sidak's multiple comparisons test. **(k)** Line graph showing image bias during the acquisition phase of the visual discrimination task, n = 15 WT mice in group A (where the image presentation is the same as that used for the testing in Figure 4 h) and 15 WT mice in group B (where the image presentation is the reverse of the used for the testing in Figure 4 h). Two way anova for Group A vs Group B p=0.088 for genotype x trial as a significant source of variation and p=0.0007 for genotype as a significant source of variation. **(l)** Line graph showing image bias during the reversal phase of the task, n = 15 WT mice in group A and 15 WT mice in group B. Two way anova for Group A vs Group B p=0.014 for genotype x trial as a significant source of variation and p=0.0003 for genotype as a significant source of variation. For bar charts, bars depict the mean and all error bars represent SEM. Stars depict level of significance with *=p<0.05, **p=<0.01 and ***p<0.0001.

### Extended data figure 9

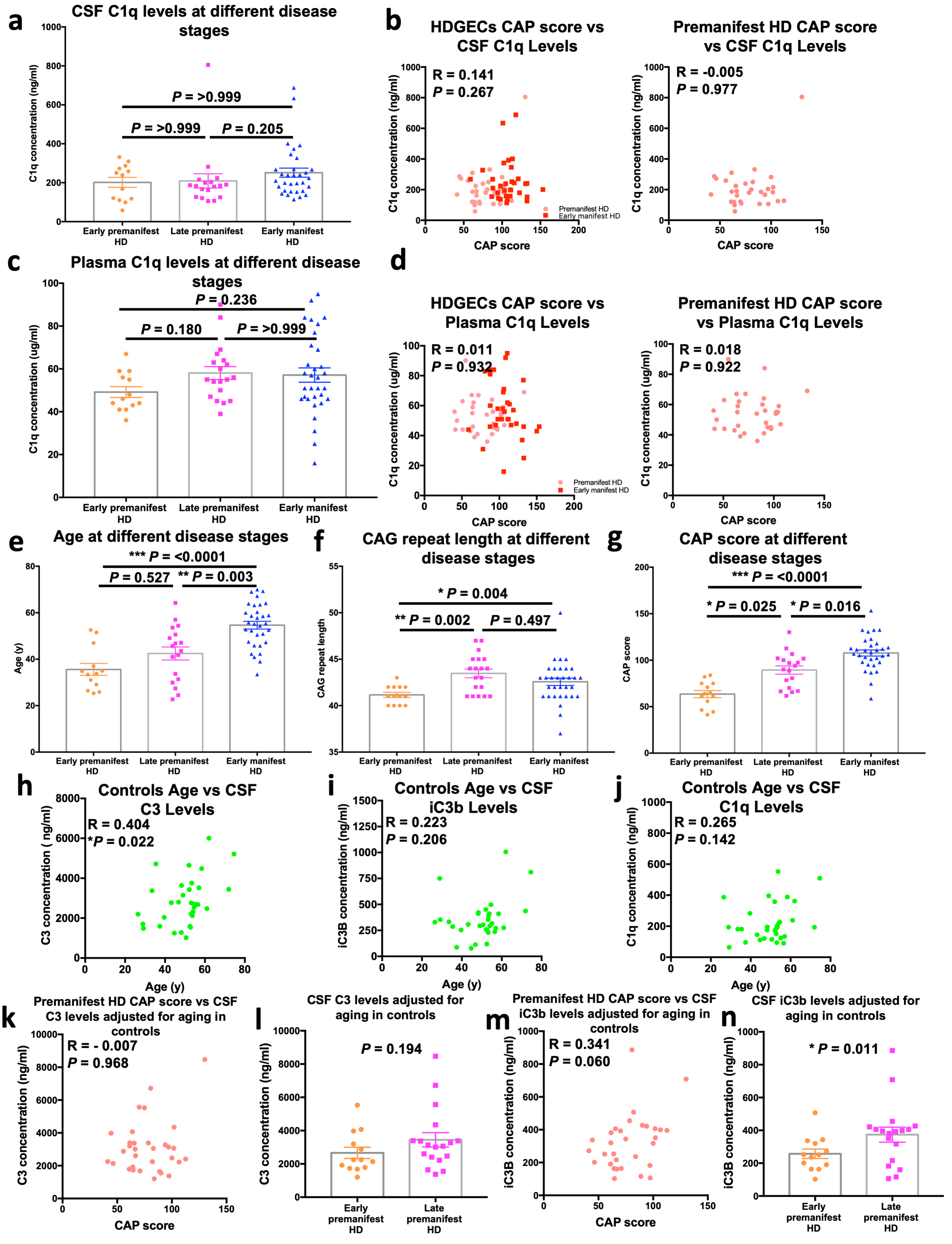

**Extended data figure 9:** **(a)** Bar chart showing the concentration of complement component C1q in CSF samples from early premanifest HD patients (see methods for inclusion criteria), late premanifest HD patients (see methods for inclusion criteria) and early manifest HD patients (see methods for inclusion criteria) recruited into the HDClarity study. Each dot represents a sample from a separate individual and the bar denotes the mean for each group n=13 early premanifest HD, n=18 late premanifest HD and n=32 early manifest HD. Kurkasl-Wallis test (non-parametric ANOVA) p=0.175 with early premanifest versus late premanifest HD p=>0.999, late premanifest versus early manifest HD p=0.205 and early premanifest versus early manifest HD p=>0.999 via Dunn's multiple comparison test. **(b)** Graphs showing the association between CAP score and CSF C1q concentration for all samples from Huntington's Disease gene expansion carriers (HDGECs) as well as those just from premanifest HD patients recruited into the HDClarity study. Each dot represents a sample from a separate individual n=63 HDGEC's and n=31 premanifest HD. Spearman r for HDGEC's p=0.267; for premanifest HD p=0.977. **(c)** Bar chart showing the concentration of complement component C1q in plasma samples from early premanifest HD patients (see methods for inclusion criteria), late premanifest HD patients (see methods for inclusion criteria) and early manifest HD patients (see methods for inclusion criteria) recruited into the HDClarity study. Each dot represents a sample from a separate individual and the bar denotes the mean for each group n=13 early premanifest HD, n=19 late premanifest HD and n=32 early manifest HD. Kurkasl-Wallis test (non-parametric ANOVA) p=0.133 with early premanifest versus late premanifest HD p = 0.180, late premanifest versus early manifest HD p=>0.999 and early premanifest versus early manifest HD p=0.236 via Dunn's multiple comparison test. **(d)** Graphs showing the association between CAP score and plasma C1q concentration for all samples from Huntington's Disease gene expansion carriers (HDGECs) as well as those just from premanifest HD patients recruited into the HDClarity study. Each dot represents a sample from a separate individual n=64 HDGEC's and n=32 premanifest HD. Spearman r for HDGEC's p=0.932; for premanifest HD p=0.922. **(e)** Bar chart showing the ages of the early premanifest HD patients (see methods for inclusion criteria), late premanifest HD patients (see methods for inclusion criteria) and early manifest HD patients (see methods for inclusion criteria) from the HDClarity study whose samples were assessed in this study. Each dot represents a sample from a separate individual and the bar denotes the mean for each group n=13 early premanifest HD, n=19 late premanifest HD and n=32 early manifest HD. Kurkasl-Wallis test (non-parametric ANOVA) p=<0.0001 with early premanifest versus late premanifest HD p=0.527, late premanifest versus early manifest HD p=0.003 and early premanifest versus early manifest HD p=<0.0001 via Dunn's multiple comparison test. **(f)** Bar chart showing the 'high' CAG repeat number of the early premanifest HD patients (see methods for inclusion criteria), late premanifest HD patients (see methods for inclusion criteria) and early manifest HD patients (see methods for inclusion criteria) from the HDClarity study whose samples were assessed in this study. Each dot represents a sample from a separate individual and the bar denotes the mean for each group n=13 early premanifest HD, n=19 late premanifest HD and n=32 early manifest HD. Kurkasl-Wallis test (non-parametric ANOVA) p=0.003 with early premanifest versus late premanifest HD p=0.002, late-premanifest versus early manifest HD p=0.497 and early premanifest versus early manifest HD p=0.004 via Dunn's multiple comparison test. **(g)** Bar chart showing the CAP score of the early premanifest HD patients (see methods for inclusion criteria), late premanifest HD patients (see methods for inclusion criteria) and early manifest HD patients (see methods for inclusion criteria) from the HDClarity study whose samples were assessed in this study. Each dot represents a sample from a separate individual and the bar denotes the mean for each group n=13 early premanifest HD, n=18 late premanifest HD and n=32 early manifest HD. Kurkasl-Wallis test (non-parametric ANOVA) p=<0.0001 with early premanifest versus late premanifest HD p=0.025, late premanifest versus early manifest HD p=0.016 and early premanifest versus early manifest HD p=<0.0001 via Dunn's multiple comparison test. **(h)** Graph showing the association between age and CSF C3 concentration for control (clinically normal) individuals recruited into the HDClarity study. Each dot represents a sample from a separate individual n=32. Spearman r p=0.022. **(i)** Graph showing the association between age and CSF iC3b concentration for control (clinically normal) individuals recruited into the HDClarity study. Each dot represents a sample from a separate individual n=32. Spearman r p=0.206. (J) Graph showing the association between age and CSF C1q concentration for control (clinically normal) individuals recruited into the HDClarity study. Each dot represents a sample from a separate individual n=32. Spearman r p=0.142. **(k)** Graph showing the association between CAP score and CSF C3 concentration (after adjustment for the effects of aging in controls) in samples from premanifest HD patients recruited into the HDClarity study. Each dot represents a sample from a separate individual n=31 premanifest HD. Spearman r p=0.968. **(l)** Bar chart showing the CSF C3 concentration (after adjustment for the effects of aging in controls) in samples from early premanifest HD and late premanifest HD patients. Each dot represents a sample from a separate individual and bars equal the mean for each group n=13 early premanifest HD and n=18 late premanifest HD. Kolmogorov-Smirnov test p=0.194. **(m)** Graph showing the association between CAP score and CSF iC3b concentration (after adjustment for the effects of aging in controls) in samples from premanifest HD patients recruited into the HDClarity study. Each dot represents a sample from a separate individual n=31 premanifest HD. Spearman r p=0.060. **(n)** Bar chart showing the CSF iC3b concentration (after adjustment for aging in controls) in samples from early premanifest HD and late premanifest HD patients. Each dot represents a sample from a separate individual and bars equal the mean for each group n=13 early premanifest HD and n=18 late premanifest HD. Kolmogorov-Smirnov test p=0.011. For bar charts, bars depict the mean. All error bars represent SEM. Stars depict level of significance with *=p<0.05, **p=<0.01 and ***p<0.0001

### Extended data figure 10

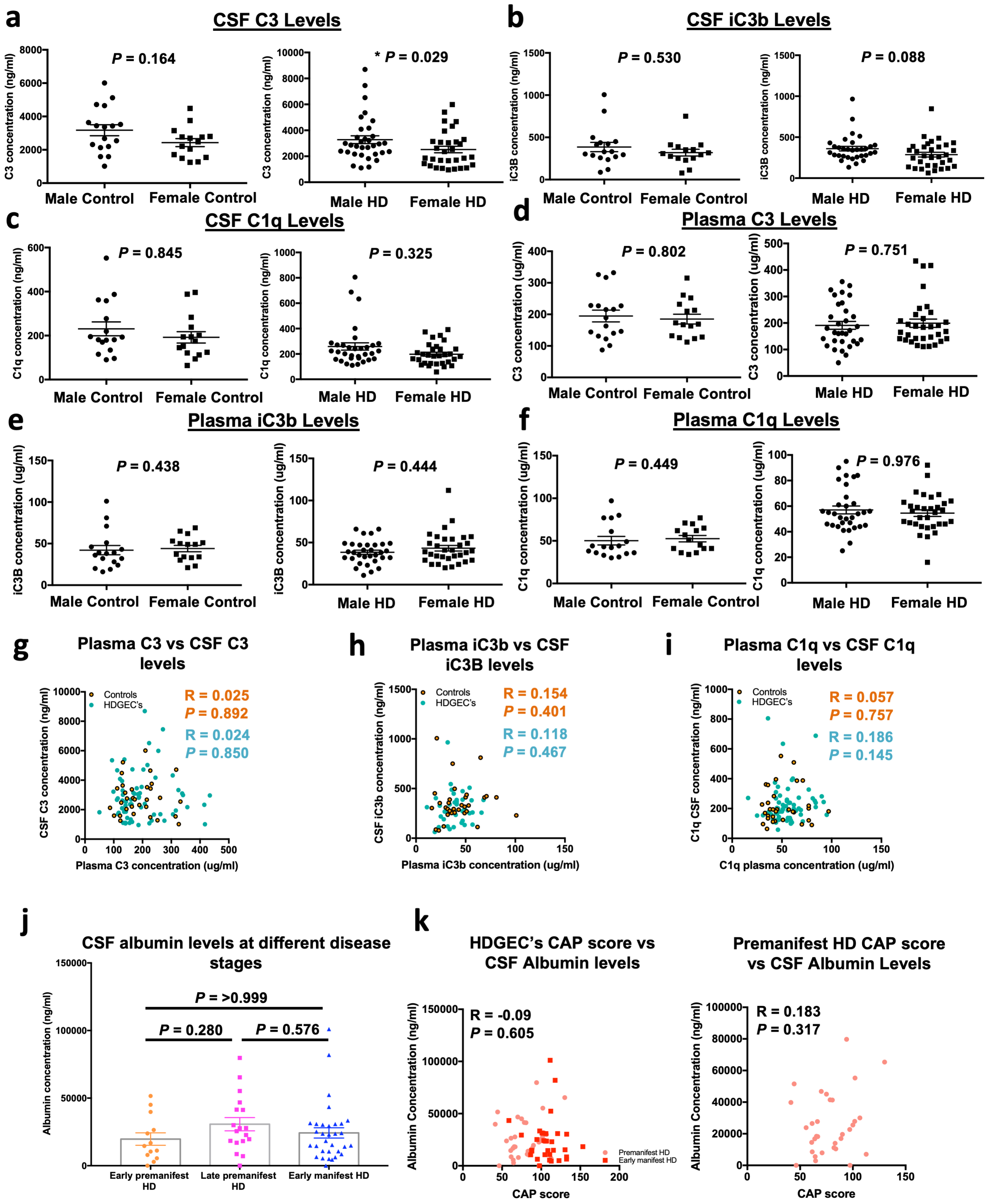

**Extended data figure 10: (a)** Graphs showing the C3 concentration in CSF samples from males and females in the control (clinically normal) group and in CSF samples from males and females with HD (both premanifest and manifest (HDGEC's)). Each dot represents a sample from a separate individual and the line denotes the mean; for the controls n=17 males and 15 females, in the HD samples n=32 males and 31 females. Kolmogorov-Smirnov test p=0.164 for the controls and p=0.029 for the HDGEC's. **(b)** Graphs showing the iC3b concentration in CSF samples from males and females in the control (clinically normal) group and in CSF samples from males and females with HD (both premanifest and manifest (HDGEC's)). Each dot represents a sample from a separate individual and the line denotes the mean; for the controls n=17 males and 15 females, in the HD samples n=32 males and 31 females. Kolmogorov-Smirnov test p=0.530 for the controls and p=0.088 for the HDGEC's. **(c)** Graphs showing the C1q concentration in CSF samples from males and females in the control (clinically normal) group and in CSF samples from males and females with HD (both premanifest and manifest (HDGEC's)). Each dot represents a sample from a separate individual and the line denotes the mean; for the controls n=17 males and 15 females, in the HD samples n=32 males and 31 females. Kolmogorov-Smirnov test p=0.845 for the controls and p=0.325 for the HDGEC's. **(d)** Graphs showing the C3 concentration in plasma samples from males and females in the control (clinically normal) group and in CSF samples from males and females with HD (both premanifest and manifest (HDGEC's)). Each dot represents a sample from a separate individual and the line denotes the mean; for the controls n=17 males and 15 females, in the HD samples n=33 males and 31 females. Kolmogorov-Smirnov test p=0.802 for the controls and p=0.751 for the HDGEC's. **(e)** Graphs showing the iC3b concentration in plasma samples from males and females in the control (clinically normal) group and in CSF samples from males and females with HD (both premanifest and manifest (HDGEC's)). Each dot represents a sample from a separate individual and the line denotes the mean; for the controls n=17 males and 15 females, in the HD samples n=33 males and 31 females. Kolmogorov-Smirnov test p=0.438 for the controls and p=0.444 for the HDGEC's. **(f)** Graphs showing the C1q concentration in plasma samples from males and females in the control (clinically normal) group and in CSF samples from males and females with HD (both premanifest and manifest (HDGEC's)). Each dot represents a sample from a separate individual and the line denotes the mean; for the controls n=17 males and 15 females, in the HD samples n=33 males and 31 females. Kolmogorov-Smirnov test p=0.449 for the controls and p=0.976 for the HDGEC's. **(g)** Graph showing the association between plasma C3 concentration and CSF C3 concentration in samples from Huntington's Disease gene expansion carriers (HDGECs) as well as those from control (clinically normal) individuals recruited into the HDClarity study. Each dot represents a sample from a separate individual, n=63 HDGEC's and n=32 controls. Spearman r for HDGEC's p=0.850; for controls p=0.892. **(h)** Graph showing the association between plasma iC3b concentration and CSF iC3b concentration in samples from Huntington's Disease gene expansion carriers (HDGECs) as well as those from control (clinically normal) individuals recruited into the HDClarity study. Each dot represents a sample from a separate individual, n=63 HDGEC's and n=32 controls. Spearman r for HDGEC's p=0.467; for controls p=0.401. **(i)** Graph showing the association between Plasma C1q concentration and CSF C1q concentration in samples from Huntington's Disease gene expansion carriers (HDGECs) as well as those from control (clinically normal) individuals recruited into the HDClarity study. Each dot represents a sample from a separate individual, n=63 HDGEC's and n=32 controls. Spearman r for HDGEC's p=0.145; for controls p=0.757. **(j)** Bar chart showing the concentration of albumin in CSF samples from early premanifest HD patients (see methods for inclusion criteria), late premanifest HD patients (see methods for inclusion criteria) and early manifest HD patients (see methods for inclusion criteria) recruited into the HDClarity study. Each dot represents a sample from a separate individual and the bar denotes the mean for each group n=13 early premanifest HD, n=18 late premanifest HD and n=32 early manifest HD. Kurkasl-Wallis test (non-parametric ANOVA) p=0.215 with early premanifest versus late premanifest HD p=0.280, late premanifest versus early manifest HD p=0.576 and early premanifest versus early manifest HD p=>0.999 via Dunn's multiple comparison test. **(k)** Graphs showing the association between CAP score and CSF albumin concentration for all samples from Huntington's Disease gene expansion carriers (HDGECs) as well as those just from premanifest HD patients recruited into the HDClarity study. Each dot represents a sample from a separate individual n=63 HDGEC's and n=31 premanifest HD. Spearman r for HDGEC's p=0.605; for premanifest HD p=0.317. For bar charts and dot plots, bars and lines depict the mean. All error bars represent SEM. Stars depict level of significance with *=p<0.05, **p=<0.01 and ***p<0.0001.

### Extended data figure 11

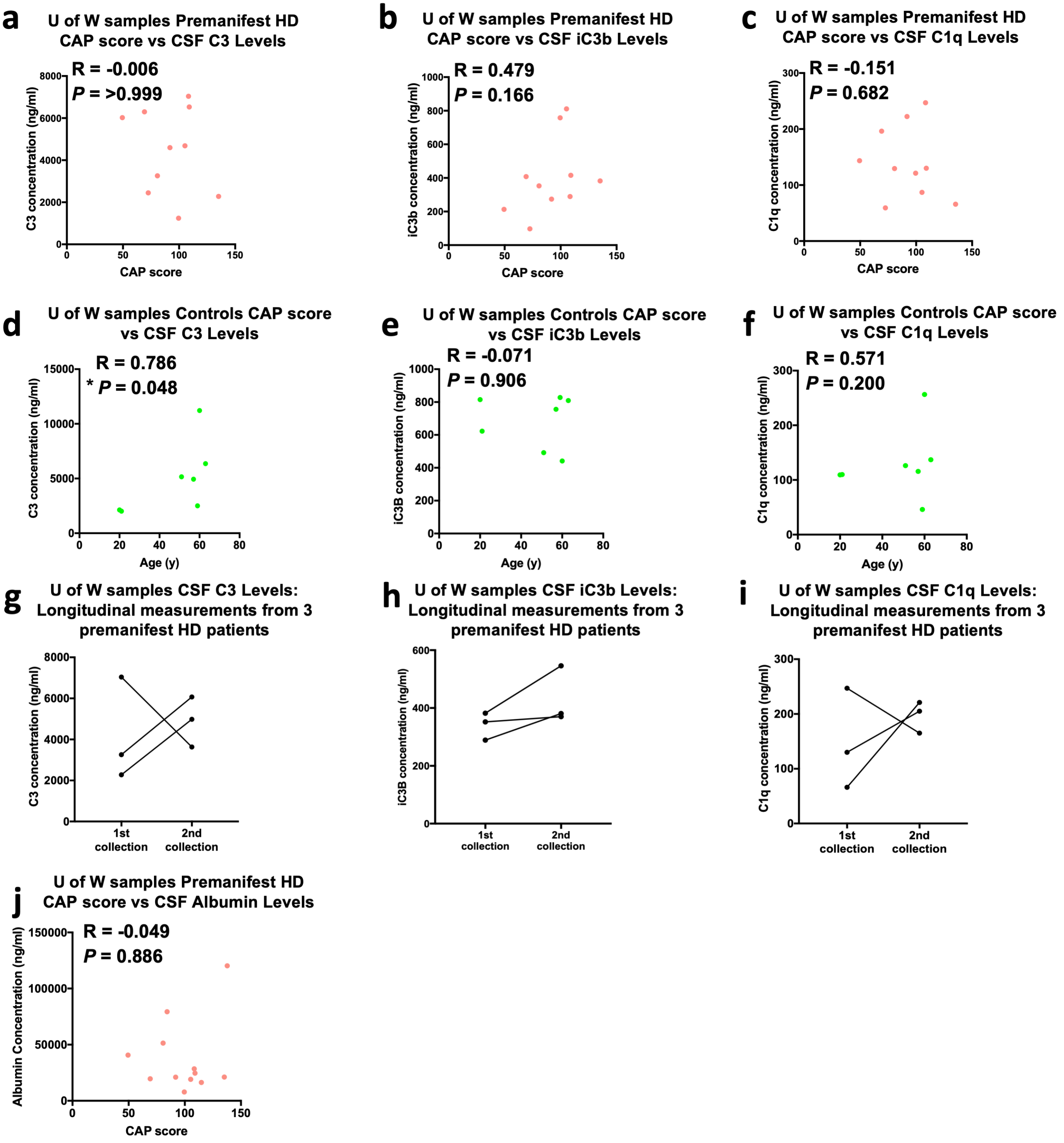

**Extended data figure 11: (a)** Graph showing the association between CAP score and CSF C3 concentration in samples from premanifest HD patients recruited into the University of Washington study. Each dot represents a sample from a separate individual n=10 Spearman r p=>0.999. **(b)** Graph showing the association between CAP score and CSF iC3b concentration in samples from premanifest HD patients recruited into the University of Washington study. Each dot represents a sample from a separate individual n=10 Spearman r p=0.166. **(c)** Graph showing the association between CAP score and CSF C1q concentration in samples from premanifest HD patients recruited into the University of Washington study. Each dot represents a sample from a separate individual n=10 Spearman r p=0.682. **(d)** Graph showing the association between patient age and CSF C3 concentration in samples from control (clinically normal) individuals recruited into the University of Washington study. Each dot represents a sample from a separate individual n=7 Spearman r p=0.048. **(e)** Graph showing the association between patient age and CSF iC3b concentration in samples from control (clinically normal) individuals recruited into the University of Washington study. Each dot represents a sample from a separate individual n=7 Spearman r p=0.906. **(f)** Graph showing the association between patient age and CSF C1q concentration in samples from control (clinically normal) individuals recruited into the University of Washington study. Each dot represents a sample from a separate individual n=7 Spearman r p=0.200. **(g)** Graph showing CSF C3 concentrations in samples from 3 premanifest HD patients from the University of Washington cohort that underwent a 2$^{nd}$ lumbar puncture approximately 1.5yr after the 1$^{st}$ sampling (see methods) n=3. **(h)** Graph showing CSF iC3b concentrations in samples from 3 premanifest HD patients from the University of Washington cohort that underwent a 2$^{nd}$ lumbar puncture approximately 1.5yr after the 1$^{st}$ sampling (see methods) n=3. **(i)** Graph showing CSF C1q concentrations in samples from 3 premanifest HD patients from the University of Washington cohort that underwent a 2$^{nd}$ lumbar puncture approximately 1.5yr after the 1$^{st}$ sampling (see methods) n=3. **(j)** Graph showing the association between CAP score and CSF albumin concentration for all premanifest samples from the University of Washington. Each dot represents a sample from a separate individual n=12 (this includes the 2$^{nd}$ collection samples referenced in panels g,h and i and excludes one sample employed in a, b and c which had a CAP score of 73, and was unfortunately no longer available). Spearman r p=0.886. Stars depict level of significance with *=p<0.05, **p=<0.01 and ***p<0.0001

### Source data figure 1

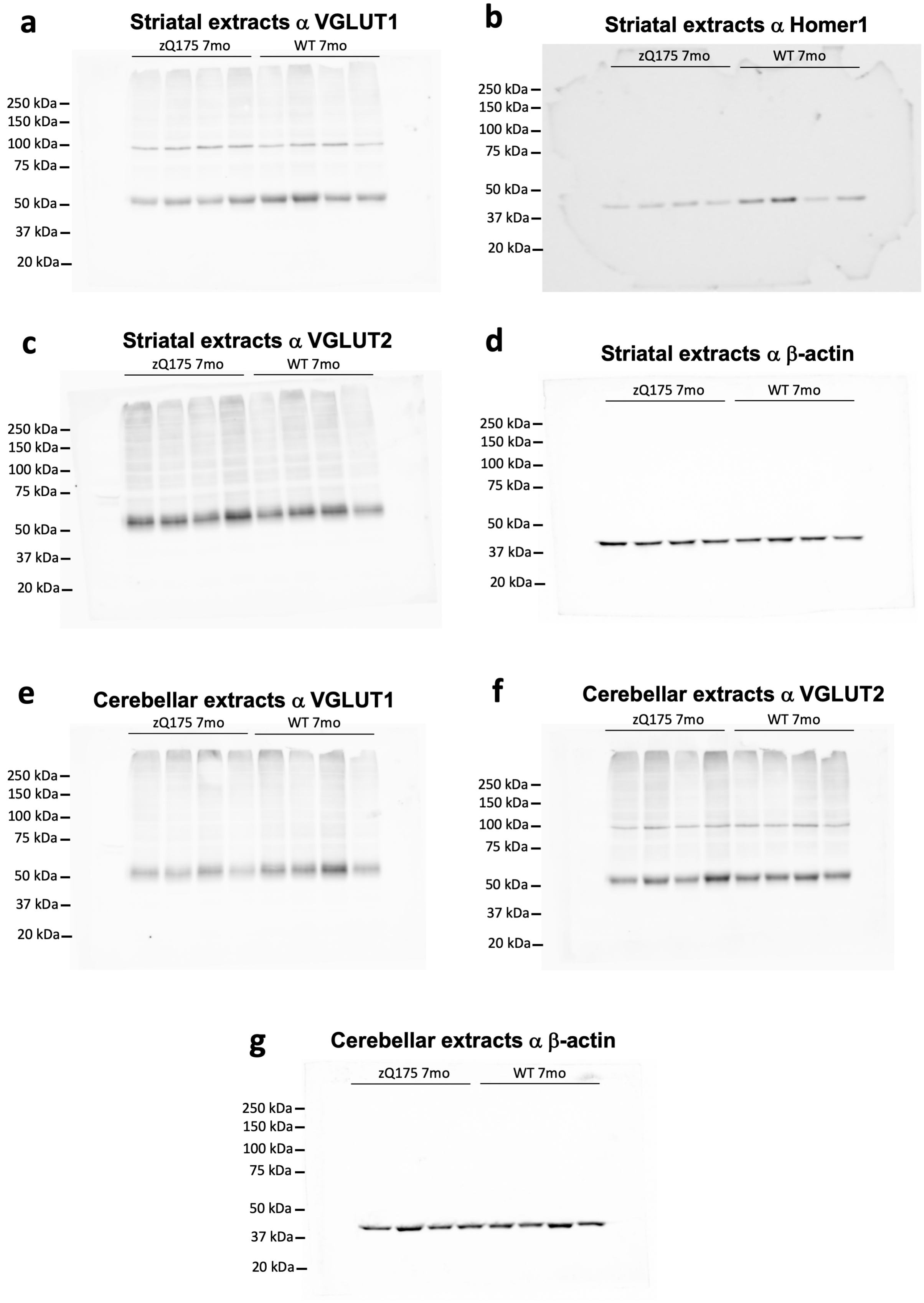

**Source data figure 1 (a)** Full length lanes of the chemiluminescent anti VGLUT1 (~55kDa) signal displayed in Extended data figure 1c; approximate molecular weights, estimated with Precision Plus Protein Kaleidoscope prestained protein standards (BIO-RAD), are shown to the left. **(b)** Full length lanes of the chemiluminescent anti Homer1 (~39kDa) signal displayed in Extended data figure 1c; approximate molecular weights, estimated with Precision Plus Protein Kaleidoscope prestained protein standards (BIO-RAD), are shown to the left. **(c)** Full length lanes of the chemiluminescent anti VGLUT2 (~56kDa) signal displayed in Extended data figure 1c; approximate molecular weights, estimated with Precision Plus Protein Kaleidoscope prestained protein standards (BIO-RAD), are shown to the left. **(d)** Full length lanes of the chemiluminescent anti β-actin (~42kDa) signal displayed in Extended data figure 1c; approximate molecular weights, estimated with Precision Plus Protein Kaleidoscope prestained protein standards (BIO-RAD), are shown to the left. **(e)** Full length lanes of the chemiluminescent anti VGLUT1 (~55kDa) signal displayed in Extended data figure 1g; approximate molecular weights, estimated with Precision Plus Protein Kaleidoscope prestained protein standards (BIO-RAD), are shown to the left. **(f)** Full length lanes of the chemiluminescent anti VGLUT2 (~56kDa) signal displayed in Extended data figure 1g; approximate molecular weights, estimated with Precision Plus Protein Kaleidoscope prestained protein standards (BIO-RAD), are shown to the left. **(g)** Full length lanes of the chemiluminescent anti β-actin (~42kDa) signal displayed in Extended data figure 1g; approximate molecular weights, estimated with Precision Plus Protein Kaleidoscope prestained protein standards (BIO-RAD), are shown to the left.

### Source data figure 2

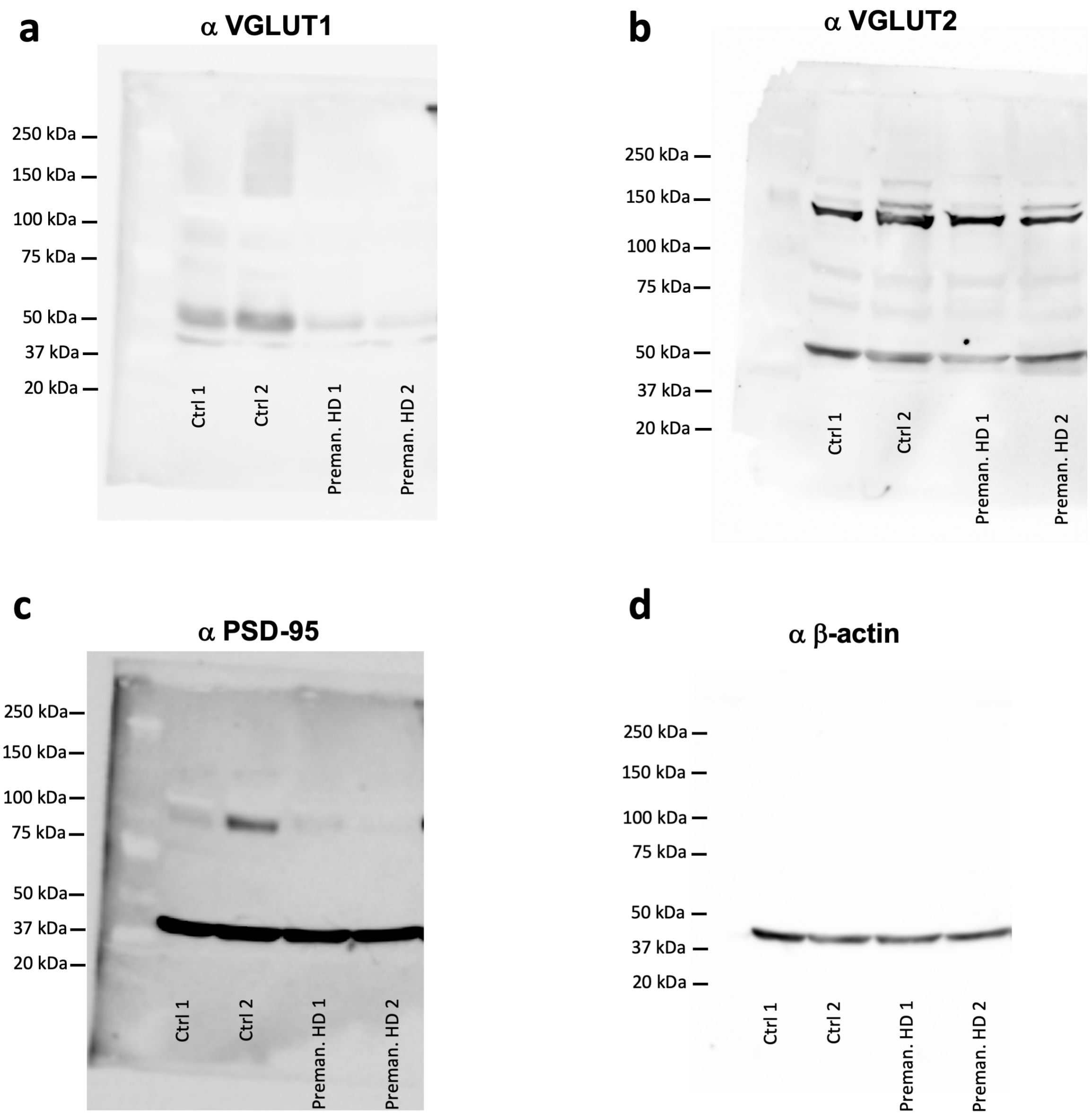

**Source data figure 2 (a)** Full length lanes of the chemiluminescent anti VGLUT1 (~55kDa) signal displayed in Extended data figure 1l; approximate molecular weights, estimated with Precision Plus Protein Kaleidoscope prestained protein standards (BIO-RAD), are shown to the left. **(b)** Full length lanes of the chemiluminescent anti VGLUT2 (~56kDa) signal displayed in Extended data figure 1l; approximate molecular weights, estimated with Precision Plus Protein Kaleidoscope prestained protein standards (BIO-RAD), are shown to the left. **(c)** Full length lanes of the chemiluminescent anti PSD-95 (~95kDa) signal displayed in Extended data figure 1l; approximate molecular weights, estimated with Precision Plus Protein Kaleidoscope prestained protein standards (BIO-RAD), are shown to the left. **(d)** Full length lanes of the chemiluminescent anti β-actin (~42kDa) signal displayed in Extended data figure 1l; approximate molecular weights, estimated with Precision Plus Protein Kaleidoscope prestained protein standards (BIO-RAD), are shown to the left. Please note that for all blots additional lanes to the right of Preman.HD 2, which contained extracts from a different source and which were run to address questions for an alternate project, have been cropped for clarity. Abbreviations: Preman. = Premanifest. Ctrl = Control
