## Supplementary material for "Microglia Mediate Early Corticostriatal Synapse Loss and Cognitive Dysfunction in Huntington’s Disease Through Complement-Dependent Mechanisms": Key resources table

| REAGENT or RESOURCE | SOURCE | IDENTIFIER |
|---|---|---|
| Antibodies | | |
| Homer1 | Synaptic systems | Cat#160-003; RRID:AB _887730 |
| VGLUT1 | Millipore Sigma | Cat#AB5905; RRID:AB_2301751 |
| VGLUT2 | Millipore Sigma | Cat#AB2251, RRID:AB_2665454 |
| PSD-95 | Millipore Sigma | Cat#MAB1596, RRID:AB_2092365 |
| Iba-1 | Wako | Cat# 019-19741, RRID:AB_839504 |
| Iba-1 | Wako | Cat# ncnp24, RRID:AB_2811160 |
| CD11b | Serotec | Cat# MCA711G; RRID:AB_321292 |
| CD68 | Serotec | Cat# MCA1957; RRID:AB_322219 |
| CD68 | Agilent Dako | Cat# M087629-2; RRID:AB_2074844 |
| P2RY12 | Anaspec | Cat# AS-55043A; RRID:AB_2298886 |
| TMEM119 | Abcam | Cat# ab209064; RRID:AB_2800343 |
| C1q | Abcam | Cat# ab182451; RRID: AB_2732849 |
| C1q | Agilent Dako | Cat# A0136; RRID: AB_2335698 |
| C1q [JL-1] | Abcam | Cat# ab71940; RRID: AB_10711046 |
| C3d | Agilent Dako | Cat# A006302; RRID: AB_578478 |
| C3c | Agilent Dako | Cat# F0201; RRID: AB_2335709 |
| iC3B | Quidel | Cat# A209; RRID: AB_452480 |
| β-actin | Millipore Sigma | Cat#A2228; RRID:AB_476697 |
| S100β | Agilent Dako | Cat#Z0311; RRID: AB_10013383 |
| Secondary antibodies, Alexa Fluor conjugates various species | Thermo Fisher Scientific - Life Technologies | Cat#'s A-11073, A-11006, A-11012, A-21245; RRID's AB_2534117, AB_2534074, AB_141359, AB_141775 |
| Digoxigenin | Roche | Cat#11207733910; RRIDAB_514500 |
| Fluorescein | Roche | Cat#11426346910; RRIDAB_840257 |
| Goat anti rabbit alkaline phosphatase | Abcam | Cat#ab97048; RRID:AB_10680574 |

| | | |
|---|---|---|
| Goat anti rabbit HRP | Promega | Cat#W4011; RRID:AB_430833 |
| Peroxidase-AffiniPure Donkey anti-guinea pig IgG (H+L) | Jackson ImmunoResearch | Cat# 706-035-148 RRID:AB_2340447 |
| Chemicals, Peptides, and Recombinant Proteins | | |
| iC3B protein | Complement technologies | Cat# A115 |
| C3c protein | Complement technologies | Cat# A116 |
| C3 protein | Complement technologies | Cat# A113 |
| C1q protein | Quidel | Cat# A400 |
| Zymosan (preactivated) | Complement technologies | Cat# B400 |
| C3/C4 inactivated serum | Complement technologies | Cat# A340 |
| C3 depleted sera | Quidel | Cat# A508 |
| C1q depleted sera | Quidel | Cat# A509 |
| NaCl | Millipore Sigma | Cat# S9888 |
| KCl | Millipore Sigma | Cat# P3911 |
| $NaHCO_3$ | Millipore Sigma | Cat# S6014 |
| $CaCl_2$ | Millipore Sigma | Cat# C1016 |
| $MgCl_2$ | Millipore Sigma | Cat# 208337 |
| $NaH_2PO_4$ | Millipore Sigma | Cat# S3139 |
| Glucose | Millipore Sigma | Cat# G8270 |
| Choline chloride | Millipore Sigma | Cat# C7017 |
| Ascorbic acid | Millipore Sigma | Cat# 1043003 |
| Pyruvic acid | Millipore Sigma | Cat# 107360 |
| InVivoMab mouse IgG1 isotype control | Bio X Cell | Cat# BE0083; RRIDAB_1107784 |
| Gabazine | Millipore Sigma | Cat# S106 |
| $CsMeSO_3$ | Millipore Sigma | Cat# C1426 |
| Hepes | Millipore Sigma | Cat# 54457 |
| EDTA | Millipore Sigma | Cat# 324626 |
| MgATP | Millipore Sigma | Cat# A9187 |
| QX-314 | Millipore Sigma | Cat# 552233 |
| Na-GTP | Millipore Sigma | Cat# 51120 |
| Phosphocreatine | Millipore Sigma | Cat# P7936 |
| CsOH | Millipore Sigma | Cat# 232068 |
| 16% paraformaldehyde | Electron microscope sciences | Cat# 15700 |
| $KH_2PO_4$ | Millipore Sigma | Cat# NIST200B |
| Sucrose | Millipore Sigma | Cat# S0389 |
| Tissue-Tek O.C.T. Compound | Electron microscope sciences | Cat# 4583 |
| Bovine serum albumin | Millipore Sigma | Cat# A2153 |
| Triton-X 100 | Millipore Sigma | Cat# T8787 |

| | | |
|---|---|---|
| Normal goat serum | Millipore Sigma | Cat# G9023 |
| Vectashield with DAPI | Vector Laboratories | Cat# H-1000 |
| 2,2'-Thiodiethanol | Millipore Sigma | Cat# 166782 |
| Dabco 33-LV | Millipore Sigma | Cat# 290734 |
| Apex Red Taq DNA Master Mix, 2.0X | Genesee Scientific | Cat# 42-138 |
| Power SYBR Green PCR Master Mix | Thermo Fisher Scientific | Cat# 4368708 |
| 1-Step PNPP Substrate solution | Thermo Fisher Scientific | Cat# 37621 |
| ReBlot Plus Strong Antibody Stripping Solution 10x | EMD Millipore | Cat# 2504 |
| Precision Plus Protein Kaleidoscope Standards | Bio-Rad | Cat# 161-0375 |
| Super signal West Dura Substrate Trial Kit | Thermo Fisher Scientific | Cat# 37076 |
| Glycine | Thermo Fisher Scientific - Fisher Bioreagents | Cat# BP381-1 |
| ROCHE cOmplete protease inhibitor cocktail | Millipore Sigma | Cat# 4693116001 |
| Sudan Black B | Millipore Sigma | Cat# 199664 |
| Glycerol | Millipore Sigma | Cat# G5516 |
| Hoechst | Thermo Fisher Scientific | Cat# H3570 |
| LR white resin | Millipore Sigma | Cat# 62661 |
| L-lysine | Millipore Sigma | Cat# L5501 |
| Sodium azide | Millipore Sigma | Cat# 71289 |
| Laemmli buffer | Bio-Rad | Cat# 161-0737 |
| Sodium hydroxide | Millipore Sigma | Cat# 221465 |
| Tris | Thermo Fisher Scientific | Cat# 15504020 |
| SDS (sodium dodecyl sulphate) | Bio-Rad | Cat# 1610301 |
| Proteinase K | Worthington Biochemical | Cat# LS004222 |
| Phenol: Chloroform: Isoamyl Alcohol | Thermo Fisher Scientific | Cat# 15593031 |
| Ethyl, alcohol, Pure | Millipore Sigma | Cat# E7023 |
| Trizma base | Millipore Sigma | Cat# T1503 |
| Methanol | Millipore Sigma | Cat# 179957 |
| Triethanolamine | Millipore Sigma | Cat # T58300 |
| Acetic anhydride | Millipore Sigma | Cat# AX0080-6 |
| Hydrogen peroxide | Millipore Sigma | Cat# 95321 |
| Experimental Models: Organisms/Strains | | |
| Mouse: zQ175/ B6J.zQ175 KI | The Jackson Laboratory | Cat#027410; RRID:IMSR JAX:027410 |
| Mouse: BACHD (and its variants BE/BR/BER[1]) | The Jackson Laboratory | Cat#008197; RRID:IMSR JAX:008197 |
| Mouse: CR3KO/ B6.129S4-Itgam$^{tm1Myd}$ [2] | The Jackson Laboratory | Cat#003991; RRID:IMSR_JAX:003991 |
| Mouse: Homer GFP[3] | From the laboratory of Professor Shigeo Okabe (Ebihara et al., 2003) | N/A |

| | | |
|---|---|---|
| Oligonucleotides | | |
| CR3 (Genotyping 1)<br>TAGGCTATCCAGAGGTAGAC | This paper | N/A |
| CR3 (Genotyping 2)<br>CATACCTGTGACCAGAAGAGC | This paper | N/A |
| CR3 (Genotyping 3)<br>ATCGCCTTCTTGACGAGTTC | This paper | N/A |
| Homer GFP (Genotyping 1)<br>CCTACGGCGTGCAGTGCTTCAGC | This paper | N/A |
| Homer GFP (Genotyping 2)<br>CGGCGAGCTGCACGCTGCGTCCTC | This paper | N/A |
| zQ175 hdh1 (Genotyping)<br>CATTCATTGCCTTGCTGCTAAG | Laragen | N/A |
| zQ175 hdh2 (Genotyping)<br>CTGAAACGACTTGAGCGACTC | Laragen | N/A |
| zQ175 Neo1 (Genotyping)<br>GATCGGCCATTGAACAAGATG | The Jackson Laboratory | |
| zQ175 Neo2 (Genotyping)<br>AGAGCAGCCGATTGTCTGTTG | The Jackson Laboratory | |
| GAPDH fw (RT-qPCR)<br>AGGTCGGTGTGAACGGATTTG | This paper | N/A |
| GAPDH rev (RT-qPCR)<br>TGTAGACCATGTAGTTGAGGTCA | This paper | N/A |
| P2RY12 mouse fw (RT-qPCR)<br>CACAGAGGGCTTTGGGAACTTA | This paper | N/A |
| P2RY12 mouse rev (RT-qPCR)<br>TGGTCCTGCTTCTGCTGAATC | This paper | N/A |
| Iba-1 mouse fw (RT-qPCR)<br>ATCAACAAGCAATTCCTCGATGA | This paper | N/A |
| Iba-1 mouse rev (RT-qPCR)<br>CAGCATTCGCTTCAAGGACATA | This paper | N/A |
| CR3 human fw (RT-qPCR)<br>GCCTTGACCTTATGTCATGGG | This paper | N/A |
| CR3 human rev (RT-qPCR)<br>CCTGTGCTGTAGTCGCACT | This paper | N/A |
| C3 human fw (RT-qPCR)<br>GGGGAGTCCCATGTACTCTATC | This paper | N/A |
| C3 human rev (RT-qPCR)<br>GGAAGTCGTGGACAGTAACAG | This paper | N/A |
| Recombinant DNA | | |
| pCMV SPORT6 C3 (human) | Open Biosystems/Horizon | Cat# MHS6278-202800305; Clone Id# 40148812 |
| pOTB7 C1QA (human) | Open Biosystems/Horizon | Cat# MHS6278-202832349; Clone Id# 4850418 |
| pOTB7 ENO2 (human) | Open Biosystems/Horizon | Cat# MHS6278-202829522; Clone Id# 3629603 |
| pCMV SPORT 6 C1qa (mouse) | Open Biosystems/Horizon | Cat# MMM1013-202704027; Clone Id# 3987436 |

| | | |
|---|---|---|
| pCMV-SPORT6 ENO2 (mouse) | Open Biosystems/Horizon | Cat# MMM1013-202703673; Clone Id# 3711960 |
| pCMV-SPORT6 C3 (mouse) | Open Biosystems/Horizon | Cat# MMM1013-202768722; Clone Id# 5134713 |
| pULTRA EGFP plasmid | Addgene/Horizon | Cat# 24129 |
| RNAscope probes | | |
| RNAscope® Probe- Mm-C3 | ACD | Cat# 417841; Transcript target region 1821-2197 |
| RNAscope® Probe- Mm-Acta2 | ACD | Cat# 319531; Transcript target region 41-1749 |
| RNAscope® 3-plex Positive Control Probe- Mm | ACD | Cat# 320881 |
| RNAscope® 3-plex Negative Control Probe | ACD | Cat# 320871 |
| Kits | | |
| Alkaline phosphatase conjugation kit | Abcam | Cat# ab102850 |
| FITC conjugation kit | Abcam | Cat# ab102884 |
| TSA staining kit | Perkin Elmer | Cat# NEL0701001KT |
| SuperScript First-Strand Synthesis System for RT-PCR | Thermo Fisher Scientific | Cat# 11904018 |
| Human Hemoglobin subunit alpha ELISA kit | Abcam | Cat# ab133999 |
| Human Complement C3 ELISA kit (for CSF) | Abcam | Cat# ab108823 |
| Human Complement C3 ELISA kit (for plasma) | Abcam | Cat# ab108822 |
| Human Albumin ELISA kit | Abcam | Cat# ab108788 |
| QuantiChrom Total Protein Assay kit | Fisher Scientific | Cat# QTPR-01K |
| BCA protein assay kit | Thermo Fisher Scientific | Cat# 23225 |
| RNEasy Mini kit | Quiagen | Cat# 74104 |
| RNAscope® Target Retrieval Reagents | ACD | Cat# 322000 |
| RNAscope® Fluorescent Multiplex Reagent Kit | ACD | Cat# 320850 |
| Viruses | | |
| AAV1/2 h-syn-EGFP | Addgene | Cat# 50465: RRID: Addgene 50465 |


1. Wang, N., *et al.* Neuronal targets for reducing mutant huntingtin expression to ameliorate disease in a mouse model of Huntington's disease. *Nat Med* **20**, 536-541 (2014).
2. Coxon, A., *et al.* A novel role for the beta 2 integrin CD11b/CD18 in neutrophil apoptosis: a homeostatic mechanism in inflammation. *Immunity* **5**, 653-666 (1996).
3. Ebihara, T., Kawabata, I., Usui, S., Sobue, K. & Okabe, S. Synchronized formation and remodeling of postsynaptic densities: long-term visualization of hippocampal neurons expressing postsynaptic density proteins tagged with green fluorescent protein. *J Neurosci* **23**, 2170-2181 (2003).
